# Lipid bilayer strengthens the cooperative network of membrane proteins

**DOI:** 10.1101/2023.05.30.542905

**Authors:** Shaima Muhammednazaar, Jiaqi Yao, Matthew R. Necelis, Yein C. Park, Zhongtian Shen, Michael D. Bridges, Ruiqiong Guo, Nicole Swope, May S. Rhee, Miyeon Kim, Kelly H. Kim, Wayne L. Hubbell, Karen G. Fleming, Linda Columbus, Seung-gu Kang, Heedeok Hong

## Abstract

Although membrane proteins fold and function in a lipid bilayer constituting cell membranes, their structure and functionality can be recapitulated in diverse amphiphilic assemblies whose compositions deviate from native membranes. It remains unclear how various hydrophobic environments can stabilize membrane proteins and whether lipids play any role therein. Here, using the evolutionary unrelated α-helical and β-barrel membrane proteins of *Escherichia coli*, we find that the hydrophobic thickness and the strength of amphiphile– amphiphile packing are critical environmental determinants of membrane protein stability. Lipid solvation enhances stability by facilitating residue burial in the protein interior and strengthens the cooperative network by promoting the propagation of local structural perturbations. This study demonstrates that lipids not only modulate membrane proteins’ stability but also their response to external stimuli.

## Main Text

The solvent environment plays a pivotal role in folding and function of proteins (*1, 2*). In the case of water-soluble proteins, the hydrophobic effect provides a critical driving force for folding inducing the cohesion of nonpolar residues in the protein interior and the expulsion of solvating water molecules to the bulk aqueous phase (*3*). Involving the collective formation and dismantling of ordered hydrogen (H)-bond networks of water, the solvent effect further mediates cooperativity in folding and allosteric protein–ligand interactions (*1, 4–6*).

In contrast to water-soluble proteins, membrane proteins fold and function in a lipid bilayer, which acts as a hydrophobic solvent-like environment in cells. The folding of helical membrane proteins can be described using the two-stage model (*7*). In stage I, transmembrane (TM) helices are formed across the bilayer. This stage is primarily driven by the hydrophobic effect inducing the burial of nonpolar polypeptide segments and by the favorable formation of backbone H-bonds in the nonpolar environment of the bilayer (*8, 9*). In stage II, the TM helices associate to form a compact native structure. Recent studies show that the denatured state ensemble (DSE) before the compaction is highly dynamic and rich in conformation: the TM helices can flip across the membrane, unfold at the water–membrane interface, or partially associate with one another (*10–14*). Since the hydrophobic effect is weak within the bilayer due to the lack of water, various molecular forces can drive this stage, including interhelical van der Waals (vdW) packing and polar interactions (*15–18*), the backbone and side-chain entropies (*19, 20*), and selective lipid binding (*21*). Membrane properties (e.g., the lateral pressure profile and lipid packing density) and membrane deformation caused by the hydrophobic thickness mismatch between the protein and bilayer are known to influence oligomerization of large membrane proteins or single-spanning TM helices (*22–28*). Nonetheless, it is not well understood how those effects involving the membrane affect the folding of multi-spanning membrane proteins (*29–31*).

The lipid composition of cell membranes is enormously heterogeneous and varies across species, organelles, the inner *vs* outer leaflet of the bilayer, and cellular responses to environmental stresses (*32–34*). While some studies suggest the importance of native lipid composition to the structure and function of membrane proteins, others point out that the fold and functionality are remarkably tolerant to variations in lipid composition and can even be recapitulated in a broad range of artificial amphiphilic assemblies (e.g., micelles, bicelles, nanodiscs, liposomes, and amphiphilic polymers) whose chemical compositions vastly deviate from the native membranes (*34–42*). Still, there are multiple examples where the structure of a given membrane protein is highly sensitive to the choice of an amphiphilic assembly (*43, 44*).

The broad spectrum of membrane proteins’ environmental sensitivity raises questions about what properties of hydrophobic environments are critical to the conformational stability of membrane proteins and whether lipids play any role therein distinguishing them from other types of amphiphiles. While recent studies focus on high-affinity, selectively bound lipids on proteins (*21, 45*), the role of low-affinity, “solvating lipids” in the folding and function of membrane proteins is largely unknown. Here, we explore the properties of hydrophobic environments that impact stage II of membrane protein folding and the effects of lipid solvation on the stability and cooperativity of membrane proteins. Cooperativity links the behaviors of distant sites (*46*) and may underlie the function of membrane proteins (e.g., ion channels, receptors, transporters, and enzymes) by enabling the propagation of physical or chemical stimuli from one site to another. We hypothesize that hydrophobic thickness of a membrane is a minimal requirement to stabilize the proteins’ secondary (stage I) and tertiary structure (stage II) and that additional membrane properties modulate the strengths of the tertiary interactions and cooperativity.

To test this hypothesis, we employed the monomeric six helical-bundle membrane protein GlpG of *E. coli*, a member of the universally conserved rhomboid protease family (*47*). We conducted a comparative analysis on the stability, cooperative network, and solvation dynamics of GlpG in two distinct hydrophobic environments widely used for structural and functional studies of membrane proteins: bicelles, discoidal bilayer fragments edge-stabilized by detergents, and detergent micelles (*48–53*). For the discovery of membrane properties that are critical to proteins’ stability and cooperativity, bicelles are an advantageous model because their physical properties can be continuously tuned by varying the lipid content (*54–58*). In addition, bicelles undergo a fast lipid exchange with one another (*59*), allowing a facile adjustment of the number of solvating lipids in response to large conformational changes of proteins such as folding. We further tested whether the modulation of stability and cooperativity by membrane properties is a generalizable hypothesis, using an evolutionary unrelated β-barrel membrane protein, OmpLA of *E. coli*. Our study reveals the pivotal role of lipid solvation in shaping the folding energy landscape and cooperative network of membrane proteins, which may provide a foundational physical principle that underlies the folding, function, and quality control of membrane proteins.

### Stability enhancement of the helical membrane protein in lipid-enriched bicelles

Measuring thermodynamic stability of a helical membrane protein (Δ*G*°_N-D_: the free energy change from the denatured to the native state in the membrane) is a daunting task due to the inherent difficulty of achieving reversible folding in a lipid environment (*31*). Here, we overcame the challenge using the steric trapping strategy, which capitalizes on the coupling of spontaneous denaturation of a doubly biotinylated protein to the simultaneous binding of two bulky monovalent streptavidin molecules (mSA, 52 kDa) (Fig. 1A for detailed description) (*60–62*). This strategy allows a thermodynamic analysis of stage II of membrane protein folding under native conditions.

**Fig. 1.**
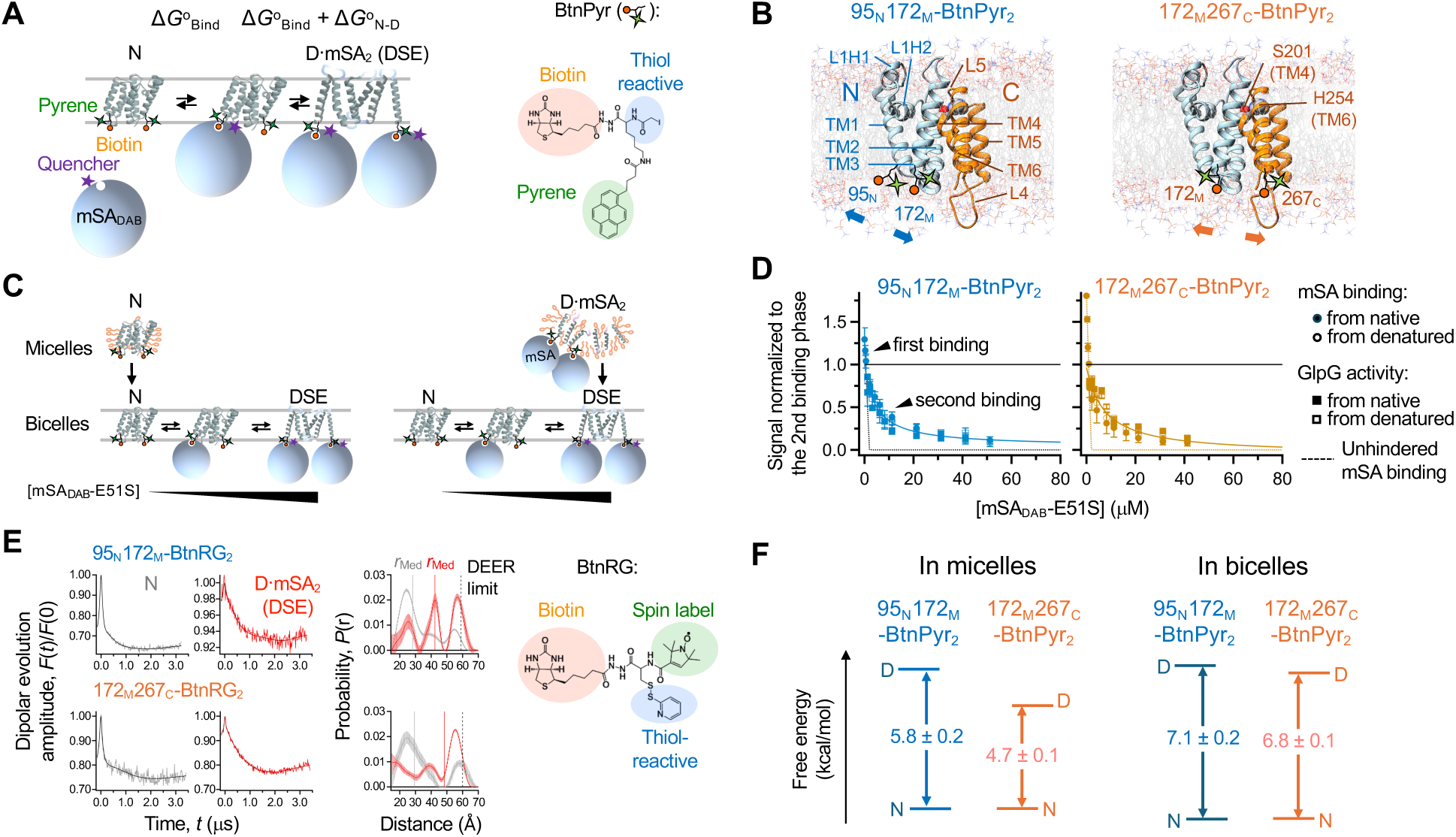
Thermodynamic stability of GlpG using steric trapping. **(A)** Steric trapping scheme. GlpG is labeled with the biotin tags (BtnPyr) at two specific residues, which are close in space in the native state and distant in the amino acid sequence. A first monovalent streptavidin (mSA, the tetramer with only one active biotin-binding site) binds unhindered to either biotin tag (Δ*G*°_Bind_). Due to the steric hindrance with the first bound mSA, a second mSA binds when the tertiary contacts between the biotinylated sites are spontaneously unraveled. The affinity of mSA to the biotin tag can be controlled by amino acid substitution on the active subunit in mSA. The coupling between GlpG denaturation and mSA binding attenuates the second binding (Δ*G*°_Bind_ + Δ*G*°_N-D_). N: native state; D×mSA2: denatured state with two bound mSA; DSE: denatured state ensemble; mSA_DAB_: mSA labeled with dabcyl quencher. **(B)** Structure of GlpG annotated with the secondary structural elements, N- and C-subdomains, the positions of the biotin pair, and the catalytic dyad for proteolysis (Ser201/His254). **(C)** Scheme for testing the reversibility of GlpG folding and mSA binding. **(D)** Binding isotherms between the double biotin variants of GlpG and mSA_DAB_-E51S. GlpG activity for the TM model substrate, LYTM2, is overlayed. **(E)** The subdomain stability of GlpG in DDM micelles (*61*) *vs* DMPC:CHAPS bicelles. Errors denote ± SEM (*N* = 4). **(F)** DEER spectroscopy. Dipolar evolution data were fitted to yield the interspin distances. The error bar at each distance denotes ± SD of the fitted probability. “DEER limit”: the maximal nominal interspin distance detectible with confidence. “*r*_Med_”: the median interspin distance.

Our main amphiphilic assemblies under comparison were the neutral bicelles composed of zwitterionic 1,2-dimyristoyl-sn-glycero-3-phosphocholine (DMPC) and 3-[(3-cholamidopropyl) dimethylammonio]-1-propanesulfonate (CHAPS) (*q* = [DMPC]/[CHAPS] = 1.5) and the neutral micelles of dodecylmaltoside (DDM). Cryo-electron microscopic analysis of bicelles without protein shows uniform discoidal particles (the average diameter, <*d*>_bicelles_ = ∼90 Å; fig. S1) indicating the formation of a lipid-enriched bilayer. DDM micelles have an oblate-spheroidal shape that is more globular and smaller than bicelles (<*d*>_micelles_ = ∼60 Å; fig. S1) (*63*).

We first validated the steric trapping strategy for measuring the stability of GlpG in bicelles. Two double cysteine variants of GlpG were generated and labeled with BtnPyr, the thiol-reactive biotin derivative with fluorescent pyrene (Fig. 1A) (*61*), resulting in the double biotin variants, 95_N_172_M_‒BtnPyr_2_ and 172_M_267_C_‒BtnPyr_2_ (95, 172, and 267: the positions of engineered cysteine residues; N, M, and C: the N-terminal, Middle, and C-terminal helices, respectively, where the cysteine residues are located). Thus, GlpG stability was measured at the N- or C-terminal half (denoted as N- and C-subdomains) depending on the position of the biotin pair (Fig. 1B) (*61*).

To construct the binding isotherm between the double biotin variant of GlpG and mSA, native or sterically denatured GlpG in DDM micelles was transferred to bicelles by dilution at an increasing concentration of mSA_DAB_-E51S (a mSA variant labeled with a dabcyl quencher which has a weaker biotin affinity than WT mSA) (Fig. 1C; fig. S2 and S3). The binding isotherm obtained by quenching of pyrene fluorescence from the BtnPyr labels (Fig. 1A and 1D) exhibits a tight unhindered first binding of mSA_DAB_-E51S followed by an optimally attenuated second binding, the latter of which is expected to be coupled to the denaturation of GlpG. In parallel, the proteolytic activity of GlpG was measured as a folding indicator. The activity of GlpG decreased as the concentration of mSA_DAB_-E51S increased and this inactivation phase was consistent with the second binding phase (Fig. 1D and fig. S4). Regardless of whether GlpG was initially folded or denatured, we obtained the same second binding phase that agreed with the inactivation phase. Therefore, the denaturation of GlpG and the second mSA binding were coupled and reversible, thereby validating the steric trapping scheme (Fig. 1A).

We further characterized the conformation of the denature state obtained by steric trapping. Upon simultaneous binding of two mSA molecules, the double-biotin variants of GlpG became more susceptible to proteolysis by Proteinase K than those without mSA, indicating an increase in conformational flexibility and water accessibility (fig. S5). To measure the degree of compactness of the denatured state, we used our spin-labeled thiol-reactive biotin label (BtnRG) (Fig. 1E) (*61*). When doubly conjugated to GlpG, BtnRG allows both the trapping of the denatured state by mSA and the measurement of interspin distances by double electron-electron resonance spectroscopy (DEER) (*10, 61*). Upon simultaneous binding of mSA, the median interspin distance between the BtnRG labels on GlpG increased from 28 Å to 42 Å for 95_N_172_M_–BtnRG_2_ and from 29 Å to 48 Å for 172_M_267_C_–BtnRG_2_ (Fig. 1E; table S1). Thus, the TM helices were largely separated in the denatured state of GlpG.

Finally, the fitting of the attenuated second binding phases yielded Δ*G*°_N-D,bicelle_^N^ = –7.1 ± 0.2 kcal/mol for N-subdomain and Δ*G*°_N-D,bicelle_^C^ = –6.8 ± 0.1 kcal/mol for C-subdomain in bicelles (Figs. 1D and 1F; fig. S6). In DDM micelles, the two subdomains exhibit different folding properties (Fig. 1F*-left*) (*61*). That is, N-subdomain (Δ*G*°_N-D,micelle_^N^ = –5.8 ± 0.2 kcal/mol), whose disruption leads to global denaturation, is more stable than C-subdomain (Δ*G*°_N-D.micelle_^C^ = –4.7 ± 0.1 kcal/mol) which undergoes subglobal denaturation (*61*). Resultantly, lipid-enriched bicelles impacted GlpG stability in two distinct ways compared to micelles: N- and C-subdomains were stabilized by –1.3 ± 0.3 kcal/mol and –2.1 ± 0.3 kcal/mol, respectively, and the stability of the two subdomains became nearly uniform (|Δ*G*°_N-D,bicelle_^N^ – Δ*G*°_N-D,bicelle_^C^ | = 0.3 ± 0.3 kcal/mol) (Fig. 1F*- right*). The stability that we measured directly under native conditions (∼–12*k*_B_*T*) is larger than that (–6.5*k*_B_*T*) extrapolated to zero force in a molecular tweezer study (*64–66*), probably due to the different conformation and degree of freedom of the denatured states (fig. S7).

### Impacts of the physical properties of amphiphilic assemblies on membrane protein stability

Next, we tracked down the molecular origin of the stabilization of GlpG in lipid-enriched bicelles. As the lipid content (*q*-value) increases in a bicelle, lipid molecules segregate into the center to form a discoidal bilayer while detergent molecules are excluded to the periphery to edge-stabilize the bilayer (*54, 67*). However, the lipid segregation is not complete such that detergents also partition into the bilayer, where the extent of partitioning depends on the lipid content (*54*). Thus, by changing the *q*-value, the physical properties of bicelles can be tuned (*54–58, 68*), offering an opportunity to identify key environmental factors that stabilize a membrane protein.

To this end, we measured GlpG stability in two different types of bicelles, DMPC:CHAPS and DMPC:DHPC (1,2-dihexanoyl-sn-glycero-3-phosphocholine), as a function of lipid content (*q*_eff_: an effective *q*-value) (*67*) (Fig. 2A; fig. S8; Methods). The change in stability was then compared to the changes in various physical properties of bicelles, including the disk thickness (*L*) (by small-angle X-ray scattering) (*54, 63, 69*), the degree of lipid segregation (by the gel–fluid phase transition temperature, *T*_m_) (*54*), and the strength of amphiphile–amphiphile packing (by the generalized polarization, *GP*) (*70, 71*) (Figs. 2B to 2D; fig. S9). All bicelle parameters were measured without GlpG. Thus, the impacts of intrinsic bicelle properties on protein stability were investigated. We note that a fluidic DMPC bilayer provides a close hydrophobic thickness-matching condition for GlpG (27–29 Å for GlpG *vs* 24–28 Å for DMPC) (*72–74*).

**Fig. 2.**
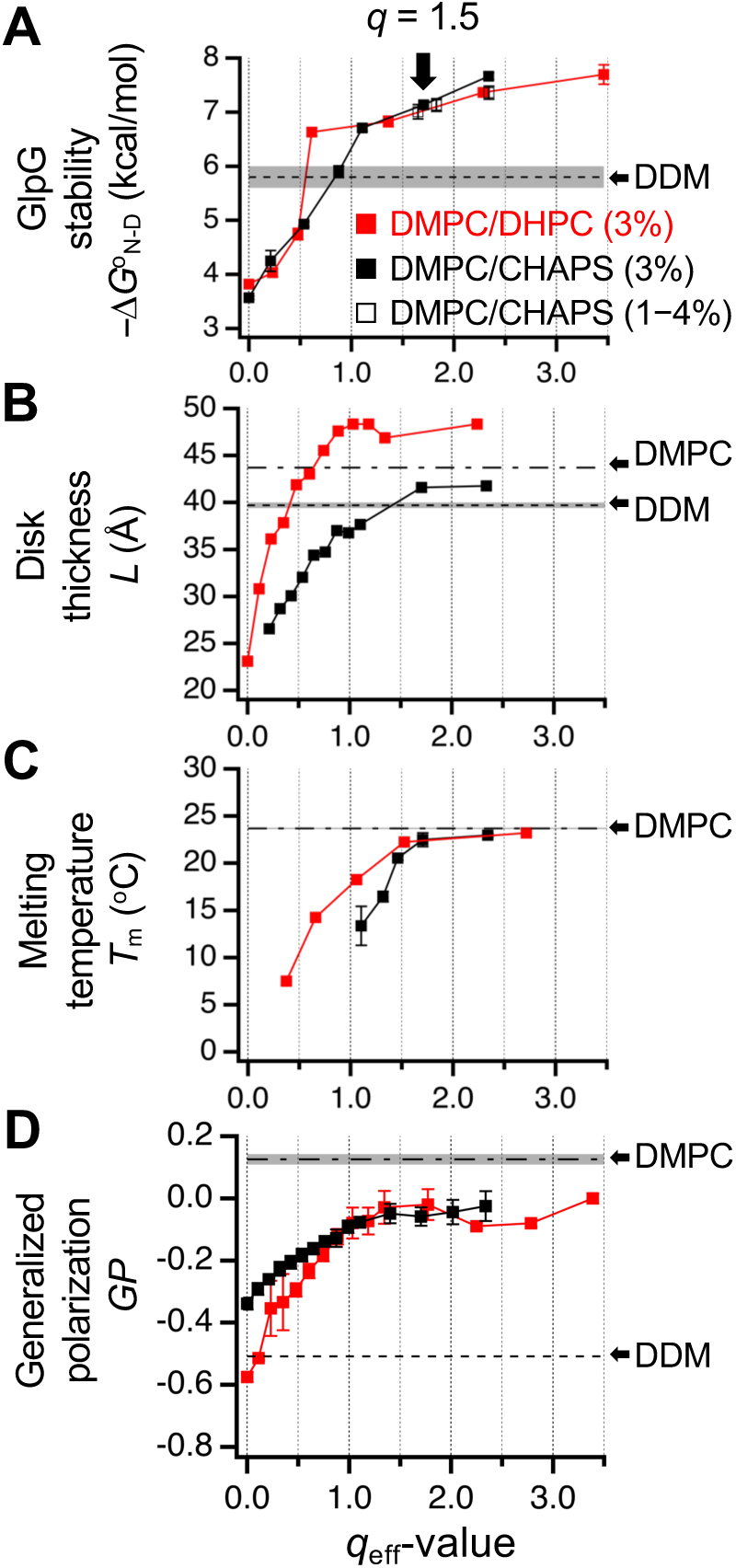
The relationship between the stability of GlpG and the physical properties of bicelles measured as a function of lipid content. The lipid content, *q*_eff_, in DMPC:CHAPS or DMPC:DHPC bicelles was calculated using the fixed critical bicelle concentrations (Eq. 7, Methods). **(A)** Stability (–Δ*G*°_N-D_) of the double-biotin variant of GlpG, 95_N_172_M_–BtnPyr_2_. The *q*_eff_ was varied either by fixing the total amphiphile concentration (DMPC:DHPC or DMPC:CHAPS) at 3 m/v-% or by diluting the bicelle solution at *q* = 1.5 (DMPC:CHAPS) to the final total amphiphile concentrations of 1 m/v-% to 4 m/v-%. **(B)** The bicelle thickness (*L*: the average headgroup–headgroup distance along the short axis of an oblate-ellipsoidal model) measured by small angle X-ray scattering. The thickness of DDM micelles was adapted from Ref. (*63*). **(C)** The gel–fluid phase transition temperature measured by fluorescence anisotropy of diphenylhexatriene (DPH) incorporated into amphiphilic assemblies. The data for DHPC:DMPC was adapted from Ref. (*54*). Errors denote ± SEM (*N* = 3 or 4). **(D)** Generalized polarization (*GP*) of Laurdan incorporated into amphiphilic assemblies.

GlpG stability was substantially lower in CHAPS and DHPC micelles (–Δ*G*°_N-D_^N^ = ∼3.5 kcal/mol at *q*_eff_ = 0) than in DDM micelles (–Δ*G*°_N-D_^N^ = 5.8 kcal/mol) (Fig. 2A). As the lipid content increased, the stability in each bicelle type steeply increased surpassing the stability level in DDM until the *q*_eff_ reached 0.6 in DMPC:DHPC or 1.1 in DMPC:CHAPS. As the *q*_eff_ increased further, the stabilities in the two bicelle types converged and then gradually increased together up to ∼8 kcal/mol. This gradual increase in stability depended only on the lipid content regardless of the features of detergents.

Interestingly, in each bicelle type, the change in GlpG stability as a function of lipid content was highly correlated with the changes in all three parameters (*L*, *T*_m_, and *GP*) (Fig. 2; fig. S10). Similar to stability, those parameters steeply increased in the low *q*_eff_ ranges (≤0.6 in DMPC:DHPC and ≤1.1 in DMPC:CHAPS) (Figs. 2B to 2D) and then saturated or gradually increased close to the corresponding values for DMPC liposomes in the higher *q*_eff_ ranges. Due to the strong correlation, it was obscure which parameter dominantly affected GlpG stability. Nonetheless, we noticed that DDM micelles provided a better thickness-matching (*L* = ∼40 Å) and stabilizing (Δ*G*°_N-D_^N^ = –5.8 kcal/mol) condition for GlpG than the low *q*_eff_ bicelles (Figs. 2A and 2B), indicating a positive correlation between stability and the degree of thickness matching. Although stabilizing the protein, DDM micelles had weak amphiphile packing (*GP* = –0.5) similar to the low *q*_eff_ bicelles (*GP* = –0.6 to –0.3) (Fig. 2D), indicating a weak correlation between stability and the strength of amphiphile packing. Thus, the improved thickness matching between GlpG and the bicelle is likely a dominant factor for the steep increase of stability in the low *q*_eff_ ranges.

In both bicelle types, the transition from the steep to the gradual increase in stability occurred at the *q*_eff_ values where the *T*_m_ exceeded ∼10 °C (∼0.6 for DMPC:DHPC and ∼1.1 for DMPC:CHAPS) (Figs. 2A and 2C). This indicates that the lipid segregation and resultant formation of a bilayer is critical to GlpG stability. While the disk thickness was saturated near the transition *q*_eff_ values (Fig. 2B), the shallow increases in the degree of lipid segregation and the strength of amphiphile packing (∼+0.8 °C/*q*_eff_ for *T*_m_ and ∼+0.02/*q*_eff_ for *GP*, Figs. 2C and 2D) noticeably increased stability, accounting for ∼25% of the total stability change.

Taken together, these results demonstrate the remarkable impact of physical properties of a hydrophobic environment on membrane protein stability. The thickness-matching between an amphiphilic assembly and the native protein has a primary importance to stability. Surprisingly, the strength of amphiphile packing, which does not directly involve an interaction with the protein, provides an additional layer of stability modulation. Detailed chemical features of constituent detergents seems less important in this modulation. The physical parameters of our main bicelles (DMPC:CHAPS at *q* = 1.5) are similar to those of DMPC liposomes (Figs. 2B to 2D). Moreover, the activity of GlpG in lipid-enriched DMPC:CHAPS bicelles surpasses the activity in DDM and DMPC:DHPC bicelles, approaching that in DMPC liposomes (fig. S11). Thus, although not perfect, our main bicelles reasonably mimic the lipid solvation on the protein in the bilayer.

### Facilitation of residue burial in the protein interior by lipid solvation

Our main micelles (DDM) and bicelles (DMPC:CHAPS at *q* = 1.5) have a similar thickness but substantially different strengths of amphiphile packing (Fig. 2). We investigated whether those features of the hydrophobic environments affect the contribution of individual residue interactions to GlpG stability. To this end, 37 residues with various degrees of burial in the protein interior were targeted mainly for large-to-small mutation. When all mutation-induced stability changes (ΔΔ*G*°_N-D,WT-Mut_) measured at N- and C-subdomains (Fig. 3A-*left*; fig. S12; tables S2 and S3) were plotted, the ΔΔ*G*°_N-D,WT-Mut_ values in micelles *vs* bicelles displayed linear correlation with a slope close to 1 (*m* = 1.1 ± 0.1). This may indicate that individual residue interactions make similar contributions to stability in the two distinct hydrophobic environments.

**Fig. 3.**
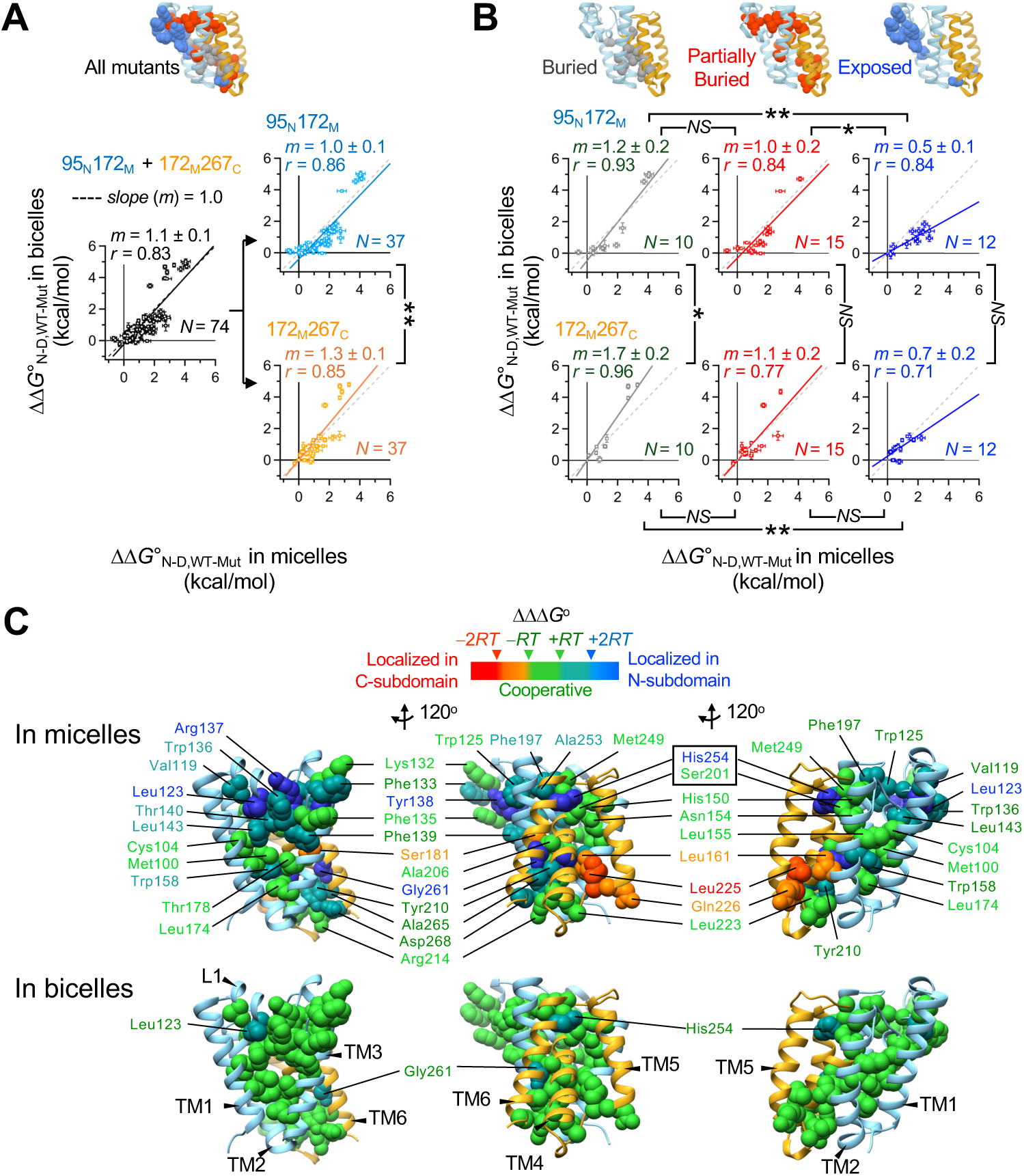
Mutation-induced stability changes (ΔΔ*G*°_N-D,WT-Mut_’s) and cooperativity profiling of GlpG in micelles *vs* bicelles. **(A)** (*Left*) All ΔΔ*G*°_N-D,WT-Mut_ values measured at N- (95_N_172_M_) and C-subdomains (172_M_267_C_) in micelles *vs* bicelles. (*Right*) ΔΔ*G*°_N-D,WT-Mut_ values depending on the location of the biotin pair. **(B)** ΔΔ*G*°_N-D,WT-Mut_ values depending on the location of the biotin pair and on the degree of burial of the mutated residues. “Buried”: *f*_ASA_, the fraction of solvent-accessible residue surface area = 0; “Partially buried”: 0 <*f*_ASA_ ≤0.1; “Exposed”: *f*_ASA_ >0.1). Errors denote ± SD from fitting. The statistical significance of difference in correlation slope (*m*) was evaluated using Chow’s test (*NS*: *p* >0.05; *: *p* ≤0.05; **: *p* ≤0.005). The *r* denotes Pearson’s coefficient. **(C)** Cooperativity profiles mapped on GlpG structure. The color code of each residue’s cooperativity profile: “cooperative” (green, ΙΔΔΔ*G*Ι≤ *RT* = 0.6 kcal/mol), “moderately localized in N-subdomain” (tin, 2*RT*≥ ΔΔΔ*G*> *RT*), “localized in N-subdomain” (blue, ΔΔΔ*G*> 2*RT*), “moderately localized in C-subdomain” (orange, –*RT*> ΔΔΔ*G*≥ –2*RT*), and “localized in C-subdomain” (red, –2*RT*> ΔΔΔ*G*). The cooperativity profiles of 20 residues in micelles have previously been assigned (*61*) (table S3).

However, the mutational impacts displayed different environmental sensitivities depending on the subdomain at which the stability was measured. That is, while mutations led to a similar degree of destabilization of N-subdomain in micelles and bicelles (*m* = 1.0 ± 0.1) (Fig. 3A-*right*), the same mutations induced the larger destabilization of C-subdomain in bicelles than in micelles (*m* = 1.3 ± 0.1) (*p*<0.005 from Chow’s test (*75*), Methods). An in-depth analysis of mutational impacts on stability based on the degree of burial of mutated residues provided a clue as to the origin of the environmental sensitivity. When the residues completely buried in the protein interior were mutated (Fig. 3B-*left*), the fitted slopes exceeded one (*m* = 1.2 ± 0.2 in N-subdomain and *m* = 1.7 ± 0.2 in C-subdomain). This tendency was more pronounced in C-subdomain than in N-subdomain (*m* = 1.7 *vs* 1.2) regardless of the position of mutation. This result implies that buried residues more favorably contribute to stability in the lipid environment than in micelles.

The different environmental sensitivity of mutational impacts observed at the two subdomains may be attributed to distinct stabilizing motifs. The stability of C-subdomain primarily arises from the extended backbone–backbone contact between the conserved Gly-zipper motifs (Gly–xxx–Gly–xxx–Gly: Gly can be Ala or Ser, and x is any residue) in TM4 and TM6 (fig. S13; Fig. 1B) (*61, 76–78*). Apparently, the more dehydrated interior of bicelles (Fig. 2D) led to the larger stabilization of the weakly polar backbone contacts between the Gly-zipper motifs (*79*) than in micelles, thereby enhancing the stability of C-subdomain (by –2.0 kcal/mol, Fig. 1F; *m* = 1.7, Fig. 3B). On the other hand, the stability of N-subdomain mainly relies on the extensive vdW contacts between large aromatic or aliphatic residues (Trp, Phe, Met, Leu, Val, Cys, and Ala) (fig. S13). This type of packing likewise induced the larger stabilization of N-subdomain in bicelles than in micelles, but to a lesser extent (by –1.2 kcal/mol, Fig. 1F; *m* = 1.2, Fig. 3B).

As the degree of exposure of mutated residues increased, the correlation slope progressively decreased to *m* = 0.5 to 0.7 (Fig. 3B) indicating that bicelles attenuated mutational impacts on stability compared to micelles. It is surprising that the perturbation of residue interactions at the protein surface differently impacts protein stability in different hydrophobic environments. This result implies that direct residue−amphiphile interactions are as important as residue−residue interactions in the protein interior to stabilizing a protein fold. The lower degree of destabilization by mutations at the lipid-contacting protein surface can be explained by either of two scenarios: *1)* lipids more favorably compensate for the structural defects created by surface mutations than detergents, or *2)* lipids provide a more adaptable solvation environment such that the protein stability in bicelles is less sensitive to the surface mutations than in micelles. Our molecular dynamics (MD) simulation supports the latter (see below).

### Strengthening of the cooperative network of the helical membrane protein by lipid solvation

Are the observed lipid effects on GlpG stability only local to the specific region under investigation (i.e., the subdomain or the site of mutation) or do they globally impact the residue interaction network? To address this question, we employed our cooperativity profiling analysis allowing us to identify whether a given residue is engaged in local or cooperative interactions with its surrounding (*61*). This experimental approach measures the degree of spatial propagation of local structural perturbation induced by a point mutation, which is quantified by the differential effect of the mutation on the stability of the two subdomains (i.e., ΔΔΔ*G*° = ΔΔ*G*°_WT-Mut_^N^ − ΔΔG°_WT-Mut_^C^) (*61*). We used four regular cutoff values, ΔΔΔ*G*° = –2*RT*, –*RT*, +*RT*, and +2*RT* (*R*: gas constant; *T*: absolute temperature) to define the cooperativity profile of each residue (Fig. 3C).

The cooperativity profiles mapped on the structure of GlpG unveiled distinct types of residue interactions, classified as “cooperative” (a mutation similarly destabilizes the two subdomains), “localized” (a mutation preferentially destabilizes the subdomain bearing the mutation), and “overpropagated” (a mutation on one subdomain induces the larger destabilization of the other). In micelles, the cooperative interactions were clustered in multiple regions (Fig. 3C): the packing core near the bilayer center (Met100, Cys104, Leu174 and Thr178) (*61*), and the narrow water channel (Ser201, Met249, His150 and Asn154) connected to the catalytic dyad (*80*). Additionally, many residues (Ala253, His254, Tyr260, Gly261, Ala265 and Asp268) at the conserved TM4/TM6 interface harboring the catalytic dyad Ser201/His254, respectively, were engaged in overpropagated interactions (*61*).

Strikingly, the cooperativity map shows a substantially different pattern in lipid-enriched bicelles (Fig. 3C). Most of the localized and overpropagated interactions in micelles turned into cooperative interactions in bicelles. Resultantly, nearly the entire set of residue-packed regions formed a single cooperative unit. The use of the narrower cutoff values (ΔΔΔ*G* = –*RT*, –1/2*RT*, +1/2*RT*, and +*RT*) yielded the cooperative clusters resembling those in micelles with the regular cutoffs (fig. S14). Thus, the cooperativity profiles in micelles were partially preserved in bicelles.

This result reveals that the lipid environment stabilizes the intraprotein interaction network more tightly than micelles, thereby facilitating the propagation of a structural perturbation throughout the protein. Thus, the observed lipid effects (i.e., the stabilization of the subdomains, the near-uniform subdomain stability, and the favorable residue burial) likely stemmed from the globally strengthened cooperative network of the protein. GlpG catalyzes proteolysis through the coordinated motions of multiple structural segments (TM4, TM6, L4, and L5) upon substrate binding (*81*). Remarkably, DMPC:CHAPS bicelles elicited a five-fold increase in GlpG activity relative to DDM micelles for both membrane-bound and water-soluble substrates (figs. S11 and S15) (*82*), which may be attributed to the augmented cooperativity in the lipid environment.

### Inefficient solvation of the helical membrane protein by lipids

To understand the molecular basis underlying the environmental dependence of GlpG stability and cooperativity, we carried out all-atom MD simulations of the GlpG–bilayer and GlpG–micelle complexes as well as the micelles alone up to 2.3 μs. In the micelle simulations, we chose two aggregation numbers of DDM (*N*_A,DDM_) per protein, 120 (DDM120) and 150 (DDM150) within the experimental range (90 to 150) (fig. S16) (*63, 69, 83*). Although an analysis of protein stability requires the investigation of both the native and denatured states, modeling and efficient sampling of the denatured states in atomistic detail are challenging for membrane proteins. Thus, our simulation focused on the native structure of GlpG.

After the conformational equilibration of the protein and amphiphiles was reached during simulation (Figs. 4A to 4C), we analyzed the solvation dynamics of amphiphiles (Fig. 4D) by calculating the time-dependent contact autocorrelation for the protein–amphiphile and amphiphile–amphiphile interactions (Fig. 4E, Methods) (*84*). The autocorrelation values decayed to <1% in all simulations, indicating that the intermolecular interactions involving amphiphiles were largely equilibrated during simulation and that DMPC mainly acted as “solvating lipids” transiently interacting with the protein. We further obtained the residence times of amphiphiles on the protein as well as on themselves (*1*_R_: the time at which the amplitude of contact autocorrelation reached 1/*e* of its initial value). Overall, lipids or detergents resided longer on GlpG (80 ns to 90 ns) than on themselves (20 ns to 40 ns), indicating a preference of amphiphiles for the protein.

**Fig. 4.**
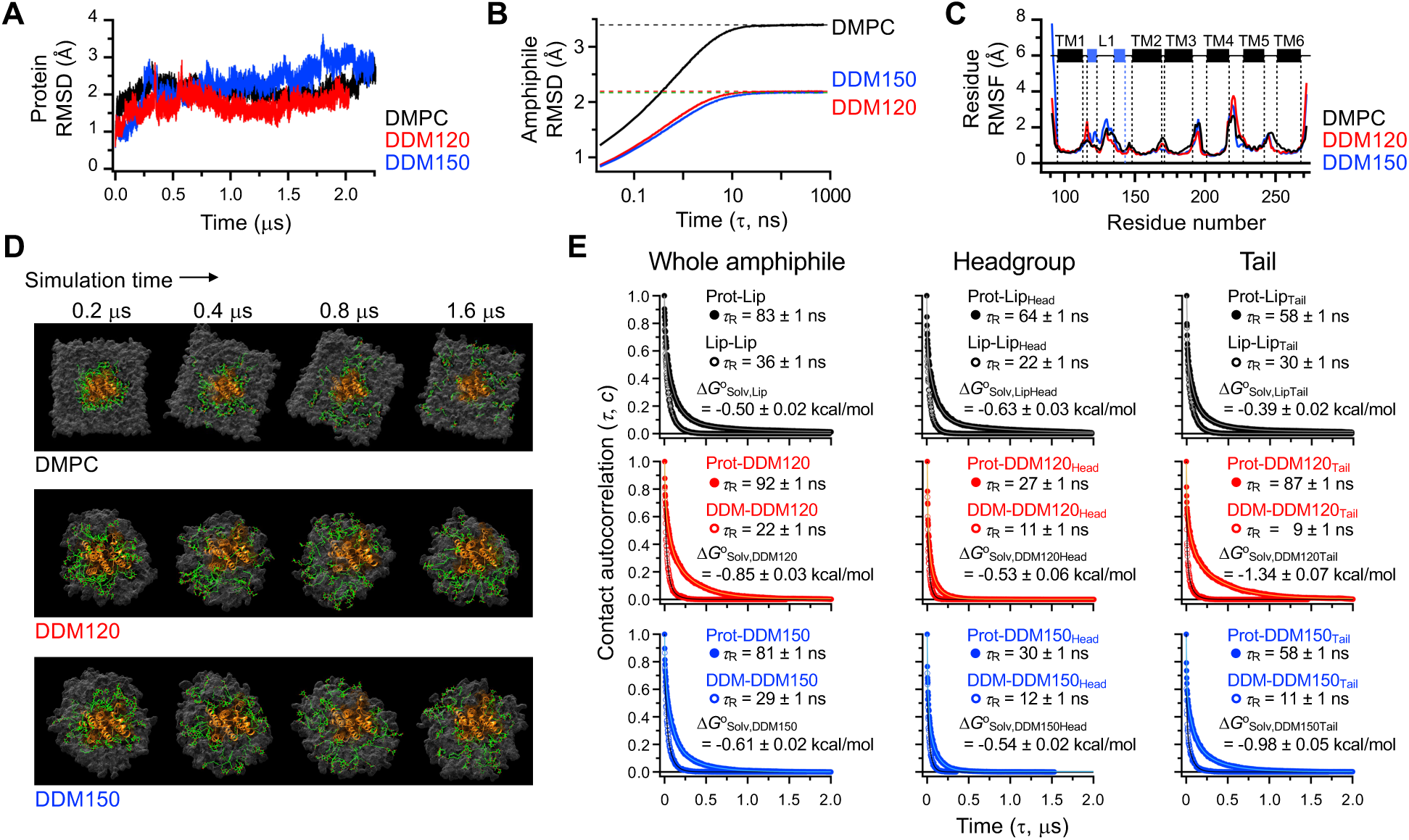
All-atom MD simulation of GlpG in the lipid bilayer and micelles. **(A)** The RMSD of the backbone heavy atoms. **(B)** The RMSD (*τ*) of lipid or detergent conformation in the bulk amphiphilic assemblies as a function of time lag *τ*. **(C)** The residue RMSF in each environment. DDM120: *N*_A,DDM_ = 120; DDM150: *N*_A,DDM_ = 150. *N*_A,DDM_: the aggregation number of DDM molecules in each micelle model. **(D)** Structural snapshots displaying the solvation dynamics of amphiphiles on GlpG in the DMPC bilayer and DDM micelles. 40 lipid or detergent molecules (green) in the first solvation shell of GlpG (orange) at the simulation time, *t* = 0.2 μs, are tracked as a function of time. In all snapshots, the viewing angle and orientation of GlpG (i.e., from the extracellular side) are fixed. **(E)** The contact autocorrelation on time for measuring the residence time (*1*_R_) of lipid or detergent molecules on GlpG or on themselves (*left*). The contacts were also separately monitored for the headgroup (*middle*) or tail (*right*) regions of amphiphiles. Each data was fitted to a triple-exponential decay function (solid lines) (table S4). Errors denote ± SD from fitting.

Based on the residence times and the equilibration of amphiphile interactions, we were able to quantify the solvation free energy (Δ*G*°_Solv_ = –*RT*·ln[*1*_R,protein-amphiphile_/*1*_R,amphiphile-amphiphile_]) per amphiphile molecule by scaling the average lifetime of amphiphile–protein contacts with the average lifetime of amphiphile–amphiphile contacts in the bulk (Fig. 4E; table S4). We expected that a lipid molecule with double hydrocarbon tails would interact more strongly with GlpG than a detergent with a single tail. However, lipids interacted with the protein more weakly than detergents (Δ*G*°_Solv,Lip_ = –0.50 ± 0.02 kcal/mol *vs* Δ*G*°_Solv,DDM120_ = –0.85 ± 0.03 kcal/mol and Δ*G*°_Solv,DDM150_ = –0.61 ± 0.02 kcal/mol) (Fig. 4E). This weaker lipid interaction stemmed from the longer *1*_R_ of lipid–lipid contacts and the comparable or shorter *1*_R_ of protein–lipid contacts than the respective *1*_R_ values of detergents. In a micelle, the increase of *N*_A,DDM_ from 120 to 150 (i.e., an increase in micellar volume) led to the weakening of detergent interaction with the protein probably due to the increased detergent-mixing entropy and the stronger interaction between detergents in the larger micelle where detergent molecules were tightly packed (fig. S16). Interestingly, the solvation on the protein was similarly driven by the headgroup and tails for lipids whereas primarily by the tail for detergents (Fig. 4E).

The MD simulations indicate that the bilayer, which is regarded as a natural solvent for membrane proteins, acts as a poorer solvent for the protein than nonnative micelles. The relatively weak lipid–protein interaction was mainly due to the lipid–lipid interactions being stronger than the detergent–detergent interactions (Fig. 2D), facilitating the dissociation of lipids from the protein (Fig. 4E). Thus, it is a plausible mechanism that the inefficient lipid solvation allows the intraprotein interactions to outweigh the lipid–protein interactions, leading to the stabilization of the compact native state (Figs. 1 to 3). That is, the residue interactions needed for stabilizing the protein’s native structure are strengthened due to the weak lipid–protein interaction that would otherwise favorably solvate the denatured state.

### Impacts of membrane properties on the folding of a β-barrel membrane protein

Finally, we tested whether the observed modulation of stability and cooperativity by membrane properties can be a working hypothesis for the folding of other types of membrane proteins. Accordingly, we employed a β-barrel membrane protein, OmpLA, which is evolutionarily unrelated to GlpG. The reversible folding of OmpLA has been established using guanidine hydrochloride (GdnHCl) as a denaturant in a 1,2-dilauroyl-sn-glycero-3-phosphocholine (DC12PC) bilayer (*85–87*). Here, we achieved the reversible folding in a 1,2-diundecanoyl-sn-glycero-3-phosphocholine (DC11PC) bilayer (fig. S17). In the unperturbed states, the DC11PC bilayer has a smaller hydrophobic thickness than the DC12PC bilayer (∼18 Å *vs* ∼20 Å) (*88*).

We first investigated how the change in bilayer thickness affects the conformational dynamics of OmpLA and lipids using all-atom MD simulations. OmpLA in the DC11PC bilayer displays larger backbone fluctuations (RMSD: ∼2 Å *vs* ∼4 Å) and a broader distribution of the tilt angles relative to the membrane normal (*θ* : 0 ° to 20° *vs* 0° to 30°) than in the DC12PC bilayer during the simulation (∼100 ns) (Figs. 5A and 5B). Moreover, compared to the DC12PC bilayer, the time-averaging of lipid conformation in the DC11PC bilayer shows a stiffening of the aliphatic chains of the solvating lipids, leading to the noticeable thickening defect in the extracellular leaflet around the protein (Fig. 5C).

**Fig. 5.**
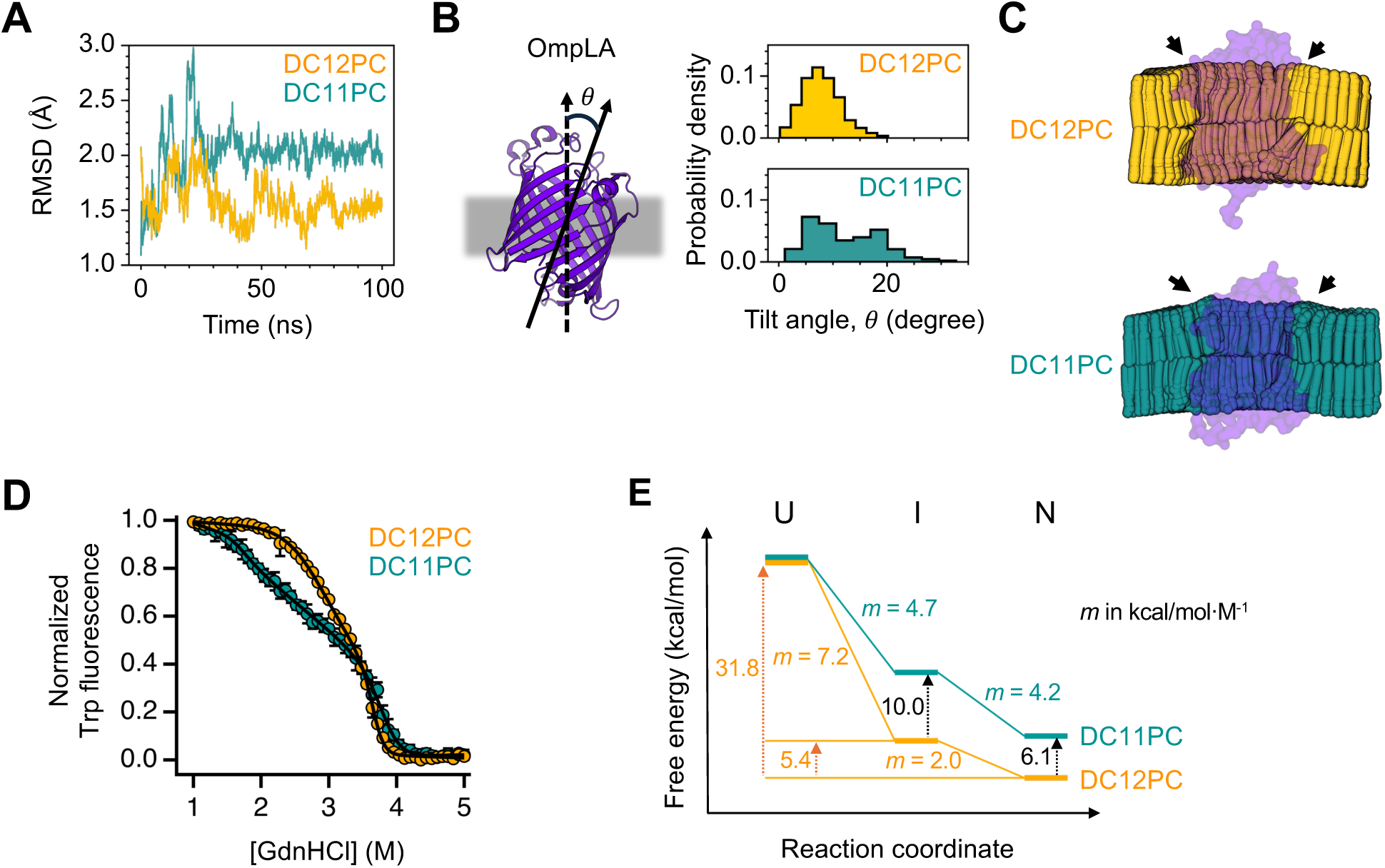
All-atom MD simulations and the stability of the β-barrel membrane protein, OmpLA, in DC12PC and DC11PC bilayers. **(A)** The RMSD of the backbone C_α_ atoms. **(B)** The distributions of the tilt angle (*θ*) between the molecular axis of OmpLA and the membrane normal. **(C)** The time-averaged lipid configurations sampled in the simulations. Black arrows indicate the bilayer thickening defects. **(D)** The folding titrations of OmpLA. Errors denote ± SEM (*N* = 3). The GdnHCl-titration data were fitted to a three-state equilibrium folding model (Eq. 18, Methods). **(E)** The folding free energy landscape of OmpLA. The free energy level of each state and the *m*-value for each transition were obtained from the fitting of the GdnHCl-titration data in Fig. 5D. The unfolded state (U), which partition to the aqueous phase, was used as a reference state.

The equilibrium folding data were fitted to a three-state model with the native (N), intermediate (I), and unfolded (U) states (*86*) (Fig. 5D; table S5). In the DC12PC bilayer, the transition free energies of OmpLA without GdnHCl were Δ*G*°_N-I,l,w_ = –5.4 ± 0.5 kcal/mol between the N and I states, both of which were formed in the membrane, and Δ*G*°_I-U,l,w_ = –26.4 ± 0.1 kcal/mol between the I state in the membrane and the U state in the aqueous phase, yielding the total free energy change of Δ*G*°_N-U,l,w_ = –31.8 ± 0.6 kcal/mol (Fig. 5E) (*86*). Compared to the DC12PC bilayer, the DC11PC bilayer substantially destabilized both the N and I states by 6.1 kcal/mol and 10.0 kcal/mol, respectively (Fig. 5E). The degree of surface exposure of protein during an unfolding transition is translated into the degree of cooperativity of the transition (*89*). When the surface exposure of OmpLA was evaluated by the *m*-value (*d*(Δ*G*°_unfold_)/*d*[GdnHCl]) (Fig. 5E) (*89*), a major portion of the protein surface area was exposed during the I-to-U transition in the DC12PC bilayer (*m*_I-U_ = 7.2 kcal/mol·M^-1^ *vs m*_N-I_ = 2.0 kcal/mol·M^-1^ for N-to-I). In contrast, a similar surface area was exposed during the N–to–I and I–to–U transitions (*m*_N-I_ and *m*_I-U_ = ∼4.5 kcal/mol·M^-1^) in the DC11PC bilayer.

Taken together, compared to the hydrophobic mismatching condition in the DC11PC bilayer, the hydrophobic matching between OmpLA and the DC12PC bilayer led to the stabilization of the N and I states in the membrane. The hydrophobic matching also induced the smaller degree of surface exposure of the I state in the membrane and the larger cooperativity of the unfolding transition involving the transfer of the I state from the membrane to the aqueous phase. This result indicates that the modulation of protein stability and cooperativity by the physical properties of a bilayer can be a common principle to both β-barrel and α-helical membrane proteins.

## Discussion and Perspectives

Here, we demonstrated the profound impact of hydrophobic environments on the folding and cooperativity of membrane proteins. Compared to micelles, lipid-enriched bicelles increase protein stability by promoting the residue burial in the protein interior and by strengthening the cooperative network. This enhancement of stability and cooperativity is linked to the formation of a lipid bilayer in a bicelle, involving the improved hydrophobic matching with the protein, the increased strength of amphiphile–amphiphile packing, and the weakening of the solvation interaction between the protein and amphiphiles, compared to micelles. These physical properties conferred by lipids may provide the formative forces for the folding energy landscape and the cooperative network of membrane proteins, unique from other types of amphiphiles. Thus, while the fold of membrane proteins is encoded by the amino acid sequence, the energetics of intraprotein interactions depend on the properties of the hydrophobic environment solvating the proteins.

Previously, we have shown that the bilayer induces contraction of the denatured state of GlpG allowing partial association of the TM helices (*10*). In this study, we observed that the lipid environment facilitates the burial of residues in the protein interior, enhancing stability. These results point to a “lipophobic effect” (i.e., an aversity of protein for lipids) in the membrane analogous to the hydrophobic effect in water. That is, the lipid environment tends to induce compaction of the polypeptide chains inserted in the membrane but not their collapse (*10*). Our MD simulation predicts that the solvation free energy of lipids on the protein (Δ*G*°_Solv_ = ∼–0.5 kcal/mol per lipid molecule) is comparable to the thermal energy (∼0.6 kcal/mol). Thus, protein interactions encoded by the amino acid sequence, which are either global or local, intra- or intermolecular, specific or nonspecific, can drive compaction if their strengths surpass thermal fluctuations.

Theories suggest that the lipophobic effect stems from perturbations of lipid conformation (e.g., the immobilization, stretching, and compression of lipid molecules) around proteins relative to the bulk lipids (*19, 90–92*). The release of the perturbed lipids to the bulk fluidic bilayer is entropically favorable, thereby driving the association of proteins to reduce the total lipid-accessible protein surface area. While this entropic effect is expected to exist for the folding of membrane proteins in various amphiphilic assemblies, our comparative study implies that lipids have a larger ability to drive folding than detergents. Although quantitative analysis of the entropic effects of amphiphiles is beyond the scope of this study, the correlation between GlpG stability and the strength of amphiphile–amphiphile packing suggests an additional physical mechanism: the large membrane tension caused by the relatively strong lipid–lipid interaction in the bilayer drives the compaction of the expanded denatured state to the native state by facilitating the release of the solvating lipids on the denatured state to the lipid–lipid packing network in the bulk.

The hydrophobic thickness mismatch between the bilayer and an embedded protein (or a single TM helix) has been recognized as an important driving force for oligomerization of large membrane proteins (*23, 25, 26*) or single-spanning TM helices (*27, 28*). Our study presents the first experimental evidence that the same physical force can modulate the stability of multi-spanning membrane proteins including both α-helical and β-barrel types. This modulatory effect by hydrophobic environments may be general to other types of amphiphilic assemblies, not limited to the lipid bilayer. For example, micelles are regarded as a highly dynamic assembly undergoing shape fluctuation, chain disordering, and an exchange of detergent molecules with the bulk water (*93*). Hence, one might expect that micellar shapes could readily be adapted to various structural features of membrane proteins. However, we show that if the intrinsic hydrophobic thickness of a micelle does not match with that of a protein, an adjustment of the micelle’s thickness to the protein’s incurs an energetic cost, leading to destabilization of the protein (Fig. 2). Thus, the physical characterization of amphiphilic assemblies will be beneficial efforts to the optimization of membrane protein stability for structural and functional studies (*54, 63, 69, 94, 95*).

The structures of GlpG crystallized in detergents and bicelles are essentially identical (RMSD: 0.65 Å) (*96*), indicating that GlpG has a robust structural fold and the key intraprotein interactions that stabilize the fold are preserved in the two hydrophobic environments. If the fold of a membrane protein is not stable enough, micelles and lipid environments may induce different structures due to the distinct physical constraints on the protein. Such examples have been thoroughly discussed in the previous literatures (*43*).

The cell membranes are composed of a diverse set of lipids with different headgroups, aliphatic chain lengths, and degrees of chain unsaturation, which collectively confer a distinct global or local hydrophobic thickness on each membrane (*32, 35, 97*). An X-ray scattering study indicate that the average hydrophobic thickness of an organelle membrane containing proteins significantly deviate from those depleted of proteins by up to ∼5 Å (*97*). This implies that the hydrophobic thicknesses of membrane proteins are not naturally matched with those of the membranes, consistently inducing strains on the membranes (*97*). Recent lipidomic, MD simulation, and protein conformational studies point out that certain types of lipids can preferentially partition on the membrane proteins modulating their oligomerization, activity, and conformational equilibrium (*23, 98–100*). Considering the negative impact of the hydrophobic mismatch on membrane protein stability observed in this study, it is plausible to hypothesize that the local membrane deformation by hydrophobic mismatch can be relieved to an extent through the preferential partitioning of hydrophobically matching lipids to the protein surface in the heterogeneous cell membranes (*23*).

In this study, the bilayer thickness and the strength of lipid packing do not exceed those of the DMPC bilayer. Thus, it is an open question, how the two competing forces (i.e., the thickness mismatching as a destabilizing force and the increased lipid packing as a stabilizing force) would impact protein stability if the bilayer thickness further increases. For an investigation of broader environmental effects induced by various types of lipids, the establishment of a detergent-free lipid bilayer system for stability measurements will be necessary. We interpreted the amphiphile and mutational effects on protein stability mainly in the context of the native structure of the protein. However, it is an essential future task to investigate how lipid and water molecules solvate the denatured states and how they impact the folding energetics of membrane proteins.

Finally, the lipid-induced enhancement of cooperativity both sheds light and cast shadows on the function, folding, and quality control of membrane proteins. For function, many membrane proteins such as ion channels, transporters, receptors, and enzymes require conformational changes (e.g., helix tilting, rotation, domain/subdomain rearrangement, etc.) upon external stimuli, which span the entire length of the protein (*101–103*). Our results indicate that the bilayer is a conducive medium to an efficient transmission of local structural perturbation throughout the protein, which can facilitate such conformational changes in a cooperative manner, thereby benefitting protein function.

In contrast, the efficient propagation of a local structural perturbation through the strengthened cooperative network can render the conformational integrity of membrane proteins particularly vulnerable to missense mutations in cells. Most of disease-causing mutations on proteins are known to be detrimental to protein stability rather than to directly disrupt active-site residues (*104–107*). Interestingly, the mapping of disease-causing mutations on the structures of G-protein coupled receptors, ion channels, and transporters shows a strong bias of finding disease mutations toward the residues in the TM regions rather than those in the extramembraneous regions (*108*). In the TM regions, this bias is even more pronounced for the residues buried in the protein interior than for the residues exposed to lipids (*108*). Our results showing the amplification of destabilizing effects by internal mutations on GlpG and the strengthened cooperative network in the bilayer environment may provide a physical basis for explaining the biased distribution of disease mutations in the interiors of membrane proteins.

## Acknowledgments

We thank the Michigan State University RTSF Cryo-EM Facility and the IBM Thomas J. Watson Research Center for their support.

## Author contributions

Conceptualization: SM, RG, KGF, LC, SGK, HH

Methodology: SM, RG, MRN, MDB, KHK, WLH, KGF, LC, SGK, HH

Investigation: SM, JY, MRN, YCP, ZS, MDB, RG, NS, MSR, MK, KHK, WLH, KGF, LC, S-gK, HH

Funding acquisition: KHK, WLH, KGF, LC, S-gK, HH

Project administration: KGF, LC, S-gK, HH

Supervision: KHK, WLH, KGF, LC, S-gK, HH

Writing – original draft: SM, MRN, YCP, ZS, MDB, MK, KHK, WLH, KGF, LC, S-gK, HH

Writing – review & editing: SM, MRN, YCP, ZS, MDB, MK, KHK, WLH, KGF, LC, S-gK, HH

## Data and materials availability

All data are available in the main text or the supplementary materials.

## Supplementary Materials

Materials and Methods Figs. S1 to S16 Tables S1 to S5 References

## References

1. Y. Levy, J. N. Onuchic, Water mediation in protein folding and molecular recognition. Annu Rev Biophys Biomol Struct 35, 389–415 (2006).

2. M. C. Bellissent-Funel et al., Water Determines the Structure and Dynamics of Proteins. Chem Rev 116, 7673–7697 (2016).

3. K. A. Dill, Dominant forces in protein folding. Biochemistry 29, 7133–7155 (1990).

4. H. S. Chan, K. A. Dill, Polymer principles in protein structure and stability. Annu Rev Biophys Biophys Chem 20, 447–490 (1991).

5. L. Jiang et al., Real-time monitoring of hydrophobic aggregation reveals a critical role of cooperativity in hydrophobic effect. Nat Commun 8, 15639 (2017).

6. A. G. Salvay, J. R. Grigera, M. F. Colombo, The role of hydration on the mechanism of allosteric regulation: in situ measurements of the oxygen-linked kinetics of water binding to hemoglobin. Biophys J 84, 564–570 (2003).

7. J. L. Popot, D. M. Engelman, Membrane protein folding and oligomerization: the two-stage model. Biochemistry 29, 4031–4037 (1990).

8. T. Hessa et al., Recognition of transmembrane helices by the endoplasmic reticulum translocon. Nature 433, 377–381 (2005).

9. Z. Cao, J. M. Hutchison, C. R. Sanders, J. U. Bowie, Backbone hydrogen bond strengths can vary widely in transmembrane helices. J Am Chem Soc 139, 10742–10749 (2017).

10. K. A. Gaffney et al., Lipid bilayer induces contraction of the denatured state ensemble of a helical-bundle membrane protein. Proc Natl Acad Sci U S A 119, (2022).

11. M. Seurig, M. Ek, G. von Heijne, N. Fluman, Dynamic membrane topology in an unassembled membrane protein. Nat Chem Biol 15, 945–948 (2019).

12. R. C. Van Lehn, B. Zhang, T. F. Miller, 3rd, Regulation of multispanning membrane protein topology via post-translational annealing. Elife 4, (2015).

13. H. Vitrac, D. M. MacLean, V. Jayaraman, M. Bogdanov, W. Dowhan, Dynamic membrane protein topological switching upon changes in phospholipid environment. Proc Natl Acad Sci U S A 112, 13874–13879 (2015).

14. N. B. Woodall, S. Hadley, Y. Yin, J. U. Bowie, Complete topology inversion can be part of normal membrane protein biogenesis. Protein Sci 26, 824–833 (2017).

15. N. H. Joh et al., Modest stabilization by most hydrogen-bonded side-chain interactions in membrane proteins. Nature 453, 1266–1270 (2008).

16. N. H. Joh, A. Oberai, D. Yang, J. P. Whitelegge, J. U. Bowie, Similar energetic contributions of packing in the core of membrane and water-soluble proteins. J Am Chem Soc 131, 10846–10847 (2009).

17. M. Mravic et al., Packing of apolar side chains enables accurate design of highly stable membrane proteins. Science 363, 1418–1423 (2019).

18. K. G. Fleming, D. M. Engelman, Specificity in transmembrane helix-helix interactions can define a hierarchy of stability for sequence variants. Proc Natl Acad Sci U S A 98, 14340–14344 (2001).

19. K. Corin, J. U. Bowie, How physical forces drive the process of helical membrane protein folding. EMBO Rep 23, e53025 (2022).

20. E. S. O’Brien et al., Membrane Proteins Have Distinct Fast Internal Motion and Residual Conformational Entropy. Angew Chem Int Ed Engl 59, 11108–11114 (2020).

21. A. Laganowsky et al., Membrane proteins bind lipids selectively to modulate their structure and function. Nature 510, 172–175 (2014).

22. S. Ballweg et al., Regulation of lipid saturation without sensing membrane fluidity. Nat Commun 11, 756 (2020).

23. R. Chadda et al., Membrane transporter dimerization driven by differential lipid solvation energetics of dissociated and associated states. Elife 10, (2021).

24. H. Hong, J. U. Bowie, Dramatic destabilization of transmembrane helix interactions by features of natural membrane environments. J Am Chem Soc 133, 11389–11398 (2011).

25. Y. Jiang et al., Membrane-mediated protein interactions drive membrane protein organization. Nat Commun 13, 7373 (2022).

26. X. Periole, T. Huber, S. J. Marrink, T. P. Sakmar, G protein-coupled receptors self-assemble in dynamics simulations of model bilayers. J Am Chem Soc 129, 10126–10132 (2007).

27. L. V. Schafer et al., Lipid packing drives the segregation of transmembrane helices into disordered lipid domains in model membranes. Proc Natl Acad Sci U S A 108, 1343–1348 (2011).

28. S. Schick et al., Assembly of the m2 tetramer is strongly modulated by lipid chain length. Biophys J 99, 1810–1817 (2010).

29. H. Hong, L. K. Tamm, Elastic coupling of integral membrane protein stability to lipid bilayer forces. Proc Natl Acad Sci U S A 101, 4065–4070 (2004).

30. M. R. Sanders, H. E. Findlay, P. J. Booth, Lipid bilayer composition modulates the unfolding free energy of a knotted alpha-helical membrane protein. Proc Natl Acad Sci U S A 115, E1799–E1808 (2018).

31. J. U. Bowie, Solving the membrane protein folding problem. Nature 438, 581–589 (2005).

32. T. Harayama, H. Riezman, Understanding the diversity of membrane lipid composition. Nat Rev Mol Cell Biol 19, 281–296 (2018).

33. J. H. Lorent et al., Plasma membranes are asymmetric in lipid unsaturation, packing and protein shape. Nat Chem Biol 16, 644–652 (2020).

34. G. van Meer, D. R. Voelker, G. W. Feigenson, Membrane lipids: where they are and how they behave. Nat Rev Mol Cell Biol 9, 112–124 (2008).

35. W. Dowhan, Molecular basis for membrane phospholipid diversity: why are there so many lipids? Annu Rev Biochem 66, 199–232 (1997).

36. T. Gonen et al., Lipid-protein interactions in double-layered two-dimensional AQP0 crystals. Nature 438, 633–638 (2005).

37. K. Matsumoto, Dispensable nature of phosphatidylglycerol in Escherichia coli: dual roles of anionic phospholipids. Mol Microbiol 39, 1427–1433 (2001).

38. A. Oberai, Y. Ihm, S. Kim, J. U. Bowie, A limited universe of membrane protein families and folds. Protein Sci 15, 1723–1734 (2006).

39. J. L. Popot, D. M. Engelman, Membranes Do Not Tell Proteins How To Fold. Biochemistry 55, 5–18 (2016).

40. C. R. Sanders, K. F. Mittendorf, Tolerance to changes in membrane lipid composition as a selected trait of membrane proteins. Biochemistry 50, 7858–7867 (2011).

41. M. Wikstrom et al., Lipid-engineered Escherichia coli membranes reveal critical lipid headgroup size for protein function. J Biol Chem 284, 954–965 (2009).

42. J. Xie, M. Bogdanov, P. Heacock, W. Dowhan, Phosphatidylethanolamine and monoglucosyldiacylglycerol are interchangeable in supporting topogenesis and function of the polytopic membrane protein lactose permease. J Biol Chem 281, 19172–19178 (2006).

43. H. X. Zhou, T. A. Cross, Influences of membrane mimetic environments on membrane protein structures. Annu Rev Biophys 42, 361–392 (2013).

44. H. X. Zhou, T. A. Cross, Modeling the membrane environment has implications for membrane protein structure and function: influenza A M2 protein. Protein Sci 22, 381–394 (2013).

45. M. Landreh, M. T. Marty, J. Gault, C. V. Robinson, A sliding selectivity scale for lipid binding to membrane proteins. Curr Opin Struct Biol 39, 54–60 (2016).

46. V. J. Hilser, D. Dowdy, T. G. Oas, E. Freire, The structural distribution of cooperative interactions in proteins: analysis of the native state ensemble. Proc Natl Acad Sci U S A 95, 9903–9908 (1998).

47. M. Freeman, The rhomboid-like superfamily: molecular mechanisms and biological roles. Annu Rev Cell Dev Biol 30, 235–254 (2014).

48. U. H. Durr, M. Gildenberg, A. Ramamoorthy, The magic of bicelles lights up membrane protein structure. Chem Rev 112, 6054–6074 (2012).

49. S. Faham, J. U. Bowie, Bicelle crystallization: a new method for crystallizing membrane proteins yields a monomeric bacteriorhodopsin structure. J Mol Biol 316, 1–6 (2002).

50. R. M. Garavito, S. Ferguson-Miller, Detergents as tools in membrane biochemistry. J Biol Chem 276, 32403–32406 (2001).

51. S. Niebling et al., Biophysical Screening Pipeline for Cryo-EM Grid Preparation of Membrane Proteins. Front Mol Biosci 9, 882288 (2022).

52. G. Ratkeviciute, B. F. Cooper, T. J. Knowles, Methods for the solubilisation of membrane proteins: the micelle-aneous world of membrane protein solubilisation. Biochem Soc Trans 49, 1763–1777 (2021).

53. C. R. Sanders, 2nd, J. H. Prestegard, Magnetically orientable phospholipid bilayers containing small amounts of a bile salt analogue, CHAPSO. Biophys J 58, 447–460 (1990).

54. T. A. Caldwell et al., Low- q Bicelles Are Mixed Micelles. J Phys Chem Lett 9, 4469-4473 (2018).

55. M. Beaugrand et al., Lipid concentration and molar ratio boundaries for the use of isotropic bicelles. Langmuir 30, 6162–6170 (2014).

56. R. D. Giudice et al., Expanding the Toolbox for Bicelle-Forming Surfactant-Lipid Mixtures. Molecules 27, (2022).

57. K. S. Mineev, K. D. Nadezhdin, S. A. Goncharuk, A. S. Arseniev, Characterization of Small Isotropic Bicelles with Various Compositions. Langmuir 32, 6624–6637 (2016).

58. H. G. Mortensen, G. V. Jensen, S. K. Hansen, T. Vosegaard, J. S. Pedersen, Structure of Phospholipid Mixed Micelles (Bicelles) Studied by Small-Angle X-ray Scattering. Langmuir 34, 14597–14607 (2018).

59. Y. Yang et al., Folding-Degradation Relationship of a Membrane Protein Mediated by the Universally Conserved ATP-Dependent Protease FtsH. J Am Chem Soc 140, 4656–4665 (2018).

60. Y. C. Chang, J. U. Bowie, Measuring membrane protein stability under native conditions. Proc Natl Acad Sci U S A 111, 219–224 (2014).

61. R. Guo et al., Steric trapping reveals a cooperativity network in the intramembrane protease GlpG. Nat Chem Biol 12, 353–360 (2016).

62. J. Yao, H. Hong, Steric trapping strategy for studying the folding of helical membrane proteins. Methods 225, 1–12 (2024).

63. R. C. Oliver et al., Dependence of micelle size and shape on detergent alkyl chain length and head group. PLoS One 8, e62488 (2013).

64. D. Min, R. E. Jefferson, J. U. Bowie, T. Y. Yoon, Mapping the energy landscape for second-stage folding of a single membrane protein. Nat Chem Biol 11, 981–987 (2015).

65. W. Lu, N. P. Schafer, P. G. Wolynes, Energy landscape underlying spontaneous insertion and folding of an alpha-helical transmembrane protein into a bilayer. Nat Commun 9, 4949 (2018).

66. H. K. Choi et al., Watching helical membrane proteins fold reveals a common N-to-C-terminal folding pathway. Science 366, 1150–1156 (2019).

67. K. J. Glover et al., Structural evaluation of phospholipid bicelles for solution-state studies of membrane-associated biomolecules. Biophys J 81, 2163–2171 (2001).

68. T. C. Gruenhagen, J. J. Ziarek, J. P. Schlebach, Bicelle size modulates the rate of bacteriorhodopsin folding. Protein Sci 27, 1109–1112 (2018).

69. J. Lipfert, L. Columbus, V. B. Chu, S. A. Lesley, S. Doniach, Size and shape of detergent micelles determined by small-angle X-ray scattering. J Phys Chem B 111, 12427–12438 (2007).

70. T. Parasassi, G. De Stasio, G. Ravagnan, R. M. Rusch, E. Gratton, Quantitation of lipid phases in phospholipid vesicles by the generalized polarization of Laurdan fluorescence. Biophys J 60, 179–189 (1991).

71. H. Orlikowska-Rzeznik, E. Krok, M. Chattopadhyay, A. Lester, L. Piatkowski, Laurdan Discerns Lipid Membrane Hydration and Cholesterol Content. J Phys Chem B 127, 3382–3391 (2023).

72. O. Engberg et al., Rhomboid-catalyzed intramembrane proteolysis requires hydrophobic matching with the surrounding lipid bilayer. Sci Adv 8, eabq8303 (2022).

73. M. A. Lomize, I. D. Pogozheva, H. Joo, H. I. Mosberg, A. L. Lomize, OPM database and PPM web server: resources for positioning of proteins in membranes. Nucleic Acids Res 40, D370–376 (2012).

74. J. Eisenblatter, R. Winter, Pressure effects on the structure and phase behavior of DMPC-gramicidin lipid bilayers: a synchrotron SAXS and 2H-NMR spectroscopy study. Biophys J 90, 956–966 (2006).

75. G. C. Chow, Tests of Equality Between Sets of Coefficients in Two Linear Regressions. Econometrica 28, 591–605 (1960).

76. S. Kim et al., Transmembrane glycine zippers: physiological and pathological roles in membrane proteins. Proc Natl Acad Sci U S A 102, 14278–14283 (2005).

77. Y. Wang, Y. Zhang, Y. Ha, Crystal structure of a rhomboid family intramembrane protease. Nature 444, 179–180 (2006).

78. R. P. Baker, S. Urban, Architectural and thermodynamic principles underlying intramembrane protease function. Nat Chem Biol 8, 759–768 (2012).

79. S. M. Anderson, B. K. Mueller, E. J. Lange, A. Senes, Combination of Calpha-H Hydrogen Bonds and van der Waals Packing Modulates the Stability of GxxxG-Mediated Dimers in Membranes. J Am Chem Soc 139, 15774–15783 (2017).

80. Y. Zhou, S. M. Moin, S. Urban, Y. Zhang, An internal water-retention site in the rhomboid intramembrane protease GlpG ensures catalytic efficiency. Structure 20, 1255–1263 (2012).

81. S. Cho, R. P. Baker, M. Ji, S. Urban, Ten catalytic snapshots of rhomboid intramembrane proteolysis from gate opening to peptide release. Nat Struct Mol Biol 26, 910–918 (2019).

82. E. Arutyunova et al., An internally quenched peptide as a new model substrate for rhomboid intramembrane proteases. Biol Chem 399, 1389–1397 (2018).

83. D. M. Kruger, S. C. L. Kamerlin, Micelle Maker: An Online Tool for Generating Equilibrated Micelles as Direct Input for Molecular Dynamics Simulations. ACS Omega 2, 4524–4530 (2017).

84. R. M. Brunne, E. Liepinsh, G. Otting, K. Wuthrich, W. F. van Gunsteren, Hydration of proteins. A comparison of experimental residence times of water molecules solvating the bovine pancreatic trypsin inhibitor with theoretical model calculations. J Mol Biol 231, 1040–1048 (1993).

85. S. K. McDonald, K. G. Fleming, Aromatic Side Chain Water-to-Lipid Transfer Free Energies Show a Depth Dependence across the Membrane Normal. J Am Chem Soc 138, 7946–7950 (2016).

86. C. P. Moon, K. G. Fleming, Side-chain hydrophobicity scale derived from transmembrane protein folding into lipid bilayers. Proc Natl Acad Sci U S A 108, 10174–10177 (2011).

87. C. P. Moon, N. R. Zaccai, P. J. Fleming, D. Gessmann, K. G. Fleming, Membrane protein thermodynamic stability may serve as the energy sink for sorting in the periplasm. Proc Natl Acad Sci U S A 110, 4285–4290 (2013).

88. L. B. Li, I. Vorobyov, T. W. Allen, The role of membrane thickness in charged protein-lipid interactions. Biochim Biophys Acta 1818, 135–145 (2012).

89. J. M. Scholtz, G. R. Grimsley, C. N. Pace, Solvent denaturation of proteins and interpretations of the m value. Methods Enzymol 466, 549–565 (2009).

90. V. Helms, Attraction within the membrane. Forces behind transmembrane protein folding and supramolecular complex assembly. EMBO Rep 3, 1133–1138 (2002).

91. P. Lague, M. J. Zuckermann, B. Roux, Lipid-mediated interactions between intrinsic membrane proteins: a theoretical study based on integral equations. Biophys J 79, 2867–2879 (2000).

92. J. P. Duneau, J. Khao, J. N. Sturgis, Lipid perturbation by membrane proteins and the lipophobic effect. Biochim Biophys Acta Biomembr 1859, 126–134 (2017).

93. M. T. Ivanovic, M. R. Hermann, M. Wojcik, J. Perez, J. S. Hub, Small-Angle X-ray Scattering Curves of Detergent Micelles: Effects of Asymmetry, Shape Fluctuations, Disorder, and Atomic Details. J Phys Chem Lett 11, 945–951 (2020).

94. M. Kieber et al., The Fluidity of Phosphocholine and Maltoside Micelles and the Effect of CHAPS. Biophys J 116, 1682–1691 (2019).

95. R. C. Oliver et al., Tuning micelle dimensions and properties with binary surfactant mixtures. Langmuir 30, 13353–13361 (2014).

96. K. R. Vinothkumar, Structure of rhomboid protease in a lipid environment. J Mol Biol 407, 232-247 (2011).

97. K. Mitra, I. Ubarretxena-Belandia, T. Taguchi, G. Warren, D. M. Engelman, Modulation of the bilayer thickness of exocytic pathway membranes by membrane proteins rather than cholesterol. Proc Natl Acad Sci U S A 101, 4083–4088 (2004).

98. H. Sawczyc et al., Lipid-polymer nanoparticles to probe the native-like environment of intramembrane rhomboid protease GlpG and its activity. Nat Commun 15, 7533 (2024).

99. C. Martens et al., Lipids modulate the conformational dynamics of a secondary multidrug transporter. Nat Struct Mol Biol 23, 744–751 (2016).

100. G. Zhang et al., Identifying Membrane Protein-Lipid Interactions with Lipidomic Lipid Exchange-Mass Spectrometry. J Am Chem Soc 145, 20859–20867 (2023).

101. D. Drew, O. Boudker, Shared Molecular Mechanisms of Membrane Transporters. Annu Rev Biochem 85, 543–572 (2016).

102. E. E. Matthews et al., Thrombopoietin receptor activation: transmembrane helix dimerization, rotation, and allosteric modulation. FASEB J 25, 2234–2244 (2011).

103. N. R. Latorraca, A. J. Venkatakrishnan, R. O. Dror, GPCR Dynamics: Structures in Motion. Chem Rev 117, 139–155 (2017).

104. S. V. Molinski et al., Comprehensive mapping of cystic fibrosis mutations to CFTR protein identifies mutation clusters and molecular docking predicts corrector binding site. Proteins 86, 833–843 (2018).

105. R. R. Kopito, D. Ron, Conformational disease. Nat Cell Biol 2, E207–209 (2000).

106. C. R. Sanders, J. K. Myers, Disease-related misassembly of membrane proteins. Annu Rev Biophys Biomol Struct 33, 25–51 (2004).

107. J. P. Schlebach et al., Conformational Stability and Pathogenic Misfolding of the Integral Membrane Protein PMP22. J Am Chem Soc 137, 8758–8768 (2015).

108. A. Oberai, N. H. Joh, F. K. Pettit, J. U. Bowie, Structural imperatives impose diverse evolutionary constraints on helical membrane proteins. Proc Natl Acad Sci U S A 106, 17747–17750 (2009).

