## Supplementary Materials for "Lipid bilayer strengthens the cooperative network of membrane proteins"

Shaima Muhammednazaar *et al.*

**The PDF file includes:**

Materials and Methods
Figs. S1 to S16
Tables S1 to S5
References

### Materials and Methods

**Expression and purification of GlpG.** *E. coli* BL21(DE3)-RP cells were transformed with pET21a vector encoding the transmembrane (TM) domain (residues 87–276) of GlpG (*1*). The cells in Luria-Bertani (LB) broth were grown at 37°C until $OD_{600nm}$ reached 0.9. Protein expression was induced at 0.5 mM Isopropyl β-D-thiogalactopyranoside (IPTG) followed by overnight culture at 15 °C. Resuspension of the total membrane pellet was solubilized by the addition of 0.7 w/v-% *n*-dodecyl-β-D-maltoside (DDM), 1 mM Tris-(2-carboxyethyl)phosphine (TCEP), 0.25 mM phenylmethylsulfonyl fluoride (PMSF), 200 mM NaCl, 50 mM 2-amino-2-(hydroxymethyl)propane-1,3-diol;dihydrochloride (Tris-HCl) buffer (pH 8.0). GlpG was purified using nickel-nitrilotriacetic acid (Ni-NTA, Qiagen) affinity chromatography in 50 mM Tris-HCl buffer (pH 8.0, 200 mM NaCl, 0.1 w/v-% DDM).

**Biotinylation of GlpG.** 50 μM of the double-cysteine variant of GlpG (P95C/G172C or G172C/V267C) was incubated with 0.5 mM TCEP for 2 h at 25 °C. The reaction was incubated overnight at 25 °C upon addition of a 40-molar excess of BtnPyr-IA in dimethyl sulfoxide (DMSO). Excess free labels were removed by washing GlpG bound to Ni-NTA resin with 0.1 w/v-% DDM and further by dialysis. Labeling efficiency was determined by measuring the absorbance of pyrene ($\varepsilon_{346nm}$ = 42,000 $M^{-1}cm^{-1}$) and protein concentration by 660 nm assay (Bio-Rad). An SDS-PAGE shift assay was carried out on ice without sample heating for single mSA-bound, double mSA-bound, and no mSA-bound GlpG by mixing 10 μL of 5 μM GlpG with 10 μL of the SDS sample buffer followed by adding 10 μL of 25 μM mSA-WT (each step incubated for 30 min).

**Preparation of monovalent streptavidin.** Detailed procedures were described previously (*1, 2*). Streptavidin (active or inactive) encoded in pET21a vector was expressed in an inclusion body form in *E. coli* BL21(DE3)RP cells. To label mSA with a thiol-reactive dabcyl quencher (dabcyl-maleimide, AnaSpec), Tyr83 near the biotin-binding pocket in the active subunit was substituted with cysteine. Active subunit: wild-type streptavidin or weaker biotin affinity variants (W79M, S45A, S27A, and E51S) with a C-terminal $His_6$ tag; Inactive subunits: the triple mutant (N23A/S27D/S45A) without $His_6$ tag (*2*).

**Expression and purification of GlpG substrate SN–LYTM2.** The procedures for expression and purification of the second TM segment of *E. coli* lactose permease fused to the C-terminus of staphylococcus nuclease (SN–LYTM2) were described previously (*1*). In LYTM2, the residue at a five residue-upstream of the scissile bond was mutated to cysteine to conjugate the thiol-reactive, environment-sensitive fluorophore, iodoacetyl-7-nitrobenz-2-oxa-1,3-diazol (IA–NBD amide, Setareh Biotech). The initial slope of the NBD fluorescence change *vs* time represented proteolytic activity of GlpG. NBD fluorescence was monitored on a SpectraMax M5e plate reader (Molecular Devices) with $\lambda_{Ex}$ = 485 nm and $\lambda_{Em}$ = 535 nm.

**Cryo-electron microscopy of bicelles and micelles.** 3 w/v-% DMPC: CHAPS bicelles ($q$ = 1.5) and w/v-3% DDM micelles were prepared in 40 mM HEPES buffer (pH 7.5, 40 mM KCl) without GlpG. Cryo-EM grids were frozen using a Vitrobot Mark IV (ThermoFisher). Briefly, 3.5 μL of each sample was applied to a glow-discharged Quantifoil Cu 1.2/1.3 holey carbon 200-mesh grid. The grid was blotted for 3.5 s prior to plunge freezing in liquid ethane. Cryo-EM images were recorded on a Talos Arctica (ThermoFisher) operated at 200 kV and equipped with a Falcon 3EC direct electron detector camera. Images were recorded in counting mode using EPU software at a nominal magnification of ×92,000 (1.12 Å/pixel), with a defocus of −2.5 mm. Micrographs were collected as single-frame images with a total exposure time of 1.5 s and a total dose of 30 electrons/Å$^2$. A total of 6,538 particles from 10 images were auto-picked and extracted into 192×192-pixel boxes. The particles were then subjected to 2D classification using cryoSPARC into 50 classes. The diameters of the bicelles and micelles in each class average were measured to obtain their size distributions.

**Preparation of native and sterically denatured GlpG in micelles.** 20 μM of the double-biotin variants of GlpG ($95_N172_M$–BtnPyr$_2$ or $172_M267_C$–BtnPyr$_2$) was incubated with 2.4 molar excess of mSA$_{DAB}$-E51S in 20 mM HEPES buffer (pH 7.5, 40 mM KCl, 0.5 mM DTT, 5 mM DDM) at 25 °C. Denaturation was monitored every 24 h using GlpG activity as a folding indicator. For $172_M267_C$–BtnPyr$_2$, maximum denaturation was reached in 24 h. For $95_N172_M$–BtnPyr$_2$, 8 mM SDS was added to facilitate the denaturation and incubated for 5 h.

**Fluorescence quenching assay to measure incorporation of GlpG into bicelles.** As a positive control for full incorporation of GlpG in bicelles, DMPC was mixed with dabcyl-1-palmitoyl-2-oleoyl-sn-glycero-3-phosphoethanolamine (dabcyl-POPE) (Avanti polar lipids) at the molar ratio of 99.5:0.5 in chloroform in a glass tube and dried under a stream of nitrogen. After further dried in vacuum for 4 h, the lipid mixture was solubilized in 500 mL of 20 mM HEPES buffer (pH 7.5), 5 w/v-% β-octylglucoside (Anatrace) at the final lipid concentration of 7.5 w/v-%. GlpG variant $95_N172_M$–BtnPyr$_2$ or $172_M267_C$–BtnPyr$_2$ in DDM was added to the resuspension and incubated on ice for 30 min. Biobeads (Bio-Rad) were added to remove the detergents in three steps (for each step, 0.2 g/mL of wet Biobeads for 6 h to 12 h at 25 °C). Resulting proteoliposomes were extruded through a 200 nm pore-size membrane. The total lipid concentration was measured using an organic phosphate assay. Based on the lipid concentration, CHAPS was added to form bicelles of $q$ = 1.5. The protein concentration was measured using a 660 nm assay. As a negative control for no incorporation, water-soluble mSA-E51S/Y83C labeled with *N*-(1-pyrene)maleimide (ThermoFisher) was used. To prepare an experimental sample, native ("N") or sterically denatured GlpG ("D·mSA$_2$") in DDM was directly injected to the solution of the bicelles containing dabcyl-POPE (20 mM HEPES buffer, pH 7.5, 40 mM KCl). In the control and experimental samples, the final concentrations of pyrene labels, DDM, and bicelles were adjusted to 1 mM, 5 mM, and 3 w/v-% ([DMPC] + [CHAPS] = 46 mM), respectively. After incubation of the mixtures at 25 °C for 24 h, pyrene fluorescence was measured with $\lambda_{Ex}$ = 345 nm and $\lambda_{Em}$ = 390 nm. The degree of quenching, which was related to the degree of GlpG incorporation into bicelles, was determined by the equation, $[F_{\text{Negative control}} - F_{\text{Experiment}}]/[F_{\text{Negative control}} - F_{\text{Positive control}}]$ ($F$: fluorescence intensity of pyrene).

**Measuring the biotin affinity of mSA variants in bicelles.** mSA variant with a weaker biotin-binding affinity, mSA$_{DAB}$-W79M (FRET acceptor) was titrated to 100 nM of the single-cysteine variants of GlpG singly labeled with BtnPyr (FRET donor) at P95C, G172C, or V267C in 20 mM HEPES buffer (pH 7.5, 3 w/v-% DMPC:CHAPS bicelles, 40 mM KCl, 0.5 mM DTT). Pyrene fluorescence was measured with $\lambda_{Ex}$ = 345 nm and $\lambda_{Em}$ = 390 nm. After 24 h, to dissociate bound mSA, excess free biotin was added to 2 mM and incubated for another 24 h. The measured pyrene fluorescence serves as a background. Background-subtracted data were fitted to **Eq. 1** to obtain $K_{d,biotin}$ of mSA$_{DAB}$-W79M in bicelles (*1*).

$$F = A1 \cdot \frac{\left(P_T + [mSA] + K_{d,biotin}\right) - \sqrt{\left(P_T + [mSA] + K_{d,biotin}\right)^2 - 4P_T \cdot [mSA]}}{2P_T} + A2$$

**Eq. 1**

, where $F$: the measured fluorescence intensity; $P_T$: the total GlpG concentration; [mSA]: the total mSA concentration; $K_{d,biotin}$: the dissociation constant of $mSA_{DAB}$ from biotin; $A1$ the total net change in fluorescence; $A2$: the fluorescence level without $mSA_{DAB}$. The $K_{d,biotin}$ of a stronger biotin-binding mSA variants (W79M, S45A, or S27A) was measured by a FRET-based competition assay. 1 μM G172C–BtnPyr was pre-equilibrated with a 2- or 5-times molar excess of a dabcyl-labeled mSA variant for 3 h at 25 °C (the quenched state). Next, a weaker biotin-affinity mSA variant without the dabcyl label was titrated against the quenched state. Dequenching of pyrene fluorescence was measured. Once equilibrium was reached (24 h to 48 h), A final 2 mM biotin was added to dissociate bound mSA and further equilibrated for 7 h to 24 h. The fluorescence data served as a background signal. Background-subtracted data were fitted to **Eq. 2** (*3*).

$$F = A1 \cdot \frac{-\left[P_T + [mSA] + \frac{K_{unlabel}}{K_{label}} \cdot \left(C_T - P_T\right)\right] + \sqrt{\left(P_T + [mSA] + \frac{K_{unlabel}}{K_{label}} \cdot \left(C_T - P_T\right)\right)^2 + 4P_T \cdot [mSA] \cdot \frac{K_{unlabel}}{K_{label}}}}{2P_T \cdot \frac{K_{unlabel}}{K_{label}}} + A2$$

**Eq. 2**

, where $K_{unlabel}$: the dissociation constant of mSA without dabcyl; $K_{label}$: the dissociation constant for $mSA_{DAB}$. Fitted values include $A1$, $A2$, and $K_{unlabel}$ or $K_{label}$. For $mSA_{DAB}$-E51S, 1.5 μM G172C–BtnPyr was first titrated with various concentrations of mSA-S27A without dabcyl, and pyrene fluorescence was measured. This signal served as a background. Then, 2 μM $mSA_{DAB}$-E51S was added, and quenching of pyrene fluorescence was measured. After reaching an equilibrium (48 h), the background-subtracted data were fitted to **Eq. 2**.

**Testing the folding reversibility of GlpG.** Native or sterically denatured GlpG in DDM micelles was directly injected into DMPC/CHAPS bicelles ($q$ = 1.5, 3 w/v-%) at various concentrations of $mSA_{Dab}$-E51S in 20 mM HEPES buffer (pH 7.5, 40 mM KCl, 0.5 mM DTT) to initiate denaturation and refolding at 25 °C, respectively. The final concentrations of GlpG, DDM, DMPC, and CHAPS were 0.5 μM, 0.1 mM to 0.2 mM, 28 mM, and 18 mM, respectively. Thus, there was one DDM molecule in every 225 to 450 DMPC and CHAPS molecules in bicelles. To monitor mSA binding, pyrene fluorescence was measured with $\lambda_{Ex}$ = 345 nm and $\lambda_{Em}$ = 390 nm every 24 h until an equilibrium was reached (48 h to 72 h). In parallel, GlpG activity as a folding indicator was measured at a 20-times molar excess of SN–LYTM2 incorporated in bicelles.

**Construction of binding isotherms to determine $\Delta G^o_{N\text{-}D}$ of GlpG.** GlpG ($95_N172_M$–$BtnPyr_2$ or $172_M267_C$–$BtnPyr_2$) in DDM was added to the bicellar solutions at various concentrations of $mSA_{DAB}$ in 20 mM HEPES buffer (pH 7.5, 3 w/v-% bicelles, 40 mM KCl, 1 mM DTT). The final concentrations of GlpG, DDM, DMPC, and CHAPS were 1.0 μM, 0.2 mM to 0.4 mM, 28 mM, and 18 mM, respectively. Thus, there was one DDM molecule in every 112.5 to 225 DMPC and CHAPS in bicelles. Depending on the stability of GlpG mutant, multiple mSA variants with a weaker biotin affinity ($mSA_{DAB}$-W79M, -S45A, -S27A, or -E51S) were screened until an optimal second binding phase in the range from 0 μM to 60 μM [mSA] was obtained. The titrated samples were transferred to a 96 well plate and incubated at 25°C. After the equilibrium was reached, the binding was measured by quenching of pyrene fluorescence with $\lambda_{Ex}$ = 345 nm and $\lambda_{Em}$ = 390 nm. Data were averaged from three fluorescence readings.

**Fitting of the second binding phase to obtain D$G^o_{N\text{-}D}$ of GlpG.** The attenuated second binding of mSA was fitted to the equation derived from the following scheme (*1, 4*):

$$N\cdot mSA \overset{K_D}{\rightleftarrows} D\cdot mSA \;\; where\; K_D = \frac{[D\cdot mSA]}{[N\cdot mSA]}$$

$$D\cdot mSA + mSA \underset{K_{d,\,biotin}}{\rightleftarrows} D\cdot 2mSA \;\; where\; K_{d,\,biotin} = \frac{[D\cdot mSA][mSA]}{[D\cdot 2mSA]} \qquad \textbf{Eq. 3}$$

The fitting equation was:

$$F = \frac{1}{\left[1 + \left(K_{d,\,biotin} + \frac{K_{d,\,biotin}}{K_D}\right)\cdot \frac{1}{[mSA]}\right]}\cdot (F_\infty - F_\circ) + F_\circ \qquad \textbf{Eq. 4}$$

$$\Delta G^o_{N-D} = -RT\cdot \ln\left(\frac{1}{K_D}\right) \qquad \textbf{Eq. 5}$$

, where $F$: the measured fluorescence intensity; $F_o$ and $F_\infty$: the fluorescence intensities from BtnPyr conjugated to GlpG at [mSA] = 0 and at [mSA] = ∞, respectively; [mSA]: the total mSA concentration; $K_{d,biotin}$: the unhindered biotin affinity of mSA; $K_D$: the equilibrium constant for denaturation of GlpG.

**Proteinase K digestion of native and sterically denatured GlpG.** Native or sterically denatured GlpG doubly labeled with BtnRG (*1*) ($95_N172_M$–$BtnRG_2$ or $172_M267_C$–$BtnRG_2$) was directly injected into DMPC:CHAPS bicelles (3 w/v-%, $q$ = 1.5) in 20 mM HEPES buffer (pH 7.5, 40 mM KCl, 5 mM DDM) at the final concentrations of 5 μM GlpG and 25 μM mSA-WT. After incubation at 25 °C for 24 h, Proteinase K was added to the final concentration of 3.4 mg/mL. The samples were withdrawn at each time point followed by the addition of 10 mM PMSF to quench proteolysis. 10 mM DTT was added and incubated for 1 h to dissociate bound mSA-WT from biotinylated GlpG by cleaving the disulfide bond. SDS-PAGE was run on ice.

**DEER for native and sterically denatured GlpG.** To label the double-cysteine variants (95C172C and 172C267C) of GlpG with the thiol-reactive paramagnetic biotin derivative, BtnRG, the GlpG stock in DDM was diluted to ~50 μM in 1.0 w/v-% DDM, 50 mM TrisHCl (pH 8.0, 200 mM NaCl), and incubated with a 2 mM TCEP-HCl for 2 h at room temperature. BtnRG in DMSO was added at a 40-times molar excess of GlpG to the mixture during gentle vortexing. Labeling reaction was incubated at room temperature overnight in the dark with gentle stirring. After labeling with BtnPyr-IA, excess free labels were removed by extensive washing of the proteins bound to $Ni^{2+}$-NTA affinity resin using 0.1 w/v-% DDM, 50 mM TrisHCl (pH 8.0), 200 mM NaCl solution. After elution with 500 mM imidazole buffer, imidazole was removed by twice running desalting column (Bio-Rad) equilibrated with 0.1 w/v-% DDM, 50 mM TrisHCl (pH 8.0), 200 mM NaCl. The samples were concentrated using a centrifugal concentrator (Millipore, MWCO = 30 kD). The final protein concentration was determined using $OD_{280nm}$, The labeling efficiency of GlpG was determined using SDS-PAGE gel shift assay (in the absence of reducing agents) in the presence and absence of mSA-WT as previously described (*1*). To obtain the sterically denatured state in DDM micelles, 120 μL of GlpG variant $95_N172_M$–$BtnRG_2$ or $172_M267_C$–$BtnRG_2$ (25 μM) was incubated with a 5-times molar excess of mSA-WT in 40 mM DDM, 20 mM HEPES (pH 7.5, 40 mM NaCl) at room temperature until the degree of denaturation reached a maximum (typically, for three days) as measured by proteolytic activity of GlpG against the model substrate SN–LYTM2. Then, native (i.e., without mSA) and sterically denatured GlpG samples were transferred to 3 w/v-% DMPC:CHAPS bicelles. The samples were finally concentrated using a centrifugal concentrator unit (MWCO = 10 kD) to 50−100 μM as measured by $OD_{280nm}$. Glycerol was added to the final concentration of 10 v/v-% for cryo-protection. Four-pulse DEER data were collected on a Q-band Bruker ELEXSYS 580 spectrometer using a 150 W amplifier and an E5106400 cavity resonator (Bruker Biospin). Samples were loaded into quartz capillaries and flash frozen in liquid nitrogen prior to data collection at 50 K. The interspin distances were determined from fits to the background-corrected dipolar evolution data using the model-free, non-negative Tikhonov regularization algorithm on the LongDistances program (https://www.biochemistry.ucla.edu/Faculty/Hubbell/software.html).

**Fluorescence anisotropy of bicelles and liposomes.** 3.0 w/v-% DMPC:CHAPS bicelles ($q$ = 0.05, 0.50, 1.19, 1.26, 1.50, and 2.0 with the total amphiphile concentrations of 46 mM to 48 mM) and DMPC liposomes (3 w/v-%, 44 mM) were prepared with diphenylhexatriene (DPH, the final concentration of 12 μM) incorporated into the amphiphilic assemblies. The final volume of each sample was 1.0 mL in 20 mM HEPES buffer (pH 7.5, 40 mM KCl). The conditions of prepared stocks were 1 mL of 80 mg/ml DMPC in chloroform, 1 ml of 0.25 mg/ml DPH in chloroform, and 1 mL of 25 w/v-% CHAPS in 40 mM HEPES buffer (pH 7.5, 40 mM KCl). Proper volumes of the DMPC and DPH stocks were mixed to each glass test tube (13×100 mm) and dried under a gentle stream of $N_2$ gas. The resulting DMPC/DPH films were further dried under vacuum overnight. The samples were then hydrated by the addition of 40 mM HEPES buffer (pH 7.5, 40 mM KCl) and resuspended by vortexing. Finally, a proper volume of the CHAPS stock was added followed by vortexing and bath-sonication (30 min at 25 °C) to form the final bicelle solution. To prepare DMPC liposomes, the hydrated lipid/DPH resuspension was extruded 21 times through a 0.2 μm pore-size polycarbonate membrane (Whatman) and stored at 4 °C. Fluorescence anisotropy was measured using a 3×3 mm quartz cuvette on a Jasco FP-8350 spectrofluorometer at $\lambda_{Ex}$ = 350 nm and $\lambda_{Em}$ = 430 nm with the excitation and emission slit widths of 5 nm (for liposomes, the excitation slit width was set to 2.5 nm). The fluorescence intensity was measured with the excitation and emission polarizers aligned in parallel and then vertical direction to each other. The $G$-factor was measured as a ratio of light scattering intensity ($\lambda_{Ex}$ = 450 nm and $\lambda_{Em}$ = 450 nm) measured with parallel to that with vertical polarizer alignments. At each temperature, anisotropy values were measured with the same sample three times and averaged. Melting curves were generated for DMPC:CHAPS bicelles and DMPC liposomes by measuring temperature-dependent fluorescence anisotropy of DPH over the range from 3 °C to 42.5 °C. Each melting curve was fitted to a sigmoid function (Igor Pro 6.4, WaveMetrics) and the fitted inflection point was taken as a melting temperature ($T_m$).

**Bicelle preparation for SAXS and Laurdan fluorescence.** DMPC (*di*-14:0):CHAPS and DMPC:DHPC (*di*-6:0) bicelles were formed with mixtures of DMPC (Avanti Polar Lipids), CHAPS (Anatrace), or DHPC (Avanti Polar Lipids). The bicelles were prepared from $q$ values (molar ratio of lipid to detergent) of 0.1 to 1.0 in increments of 0.1 and from $q$ = 1.0 to 2.0 in increments of 0.25 (fluorescence) and 0.5 (small-angle X-ray scattering, SAXS). All samples for fluorescence were prepared at 3 w/v-% total amphiphile in phosphate buffer (137 mM NaCl, 2.7 mM KCl, 8 mM $Na_2HPO_4$, 2 mM $KH_2PO_4$, pH 7.4; PBS). Samples prepared for SAXS were prepared at 6 w/v-% total amphiphile in PBS. DMPC and Laurdan in a powder form were dissolved into chloroform stocks before mixing to a molar ratio of 1:400 (Laurdan:amphiphile). Excess chloroform was evaporated with $N_2$ gas and subsequently desiccated under vacuum for >16 h. CHAPS and DHPC detergents in a powder form were dissolved in PBS, and subsequently added to the dried lipid film at amounts to achieve specific $q$-values. After samples were brought to a final volume of 0.25 mL, they were vortexed for 60 s. Samples were subjected to three freeze-thaw cycles between liquid $N_2$ and 40°C, with vortexing for 60 s after each thaw.

**SAXS collection and data processing.** Synchrotron SAXS data were measured with the 12-ID-C beamline at the Advanced Photon Source of the Argonne National Laboratory. Samples were loaded into quartz capillary tubes (2.0 mm OD) with 0.01 mm wall thickness (Charles Supper) for data collection. The incident photon energy was 12 keV (wavelength = 1 Å), and the sample-to-detector distance was adjusted to provide a scattering wave-vector range $Q$ of $0.006 < Q < 1.079$ Å$^{-1}$. A mosaic X-ray charge-coupled device detector was used to acquire images with typical exposure times of 1.0 s. A temperature-controlled sample holder with 13 capillary slots was used to record data at 25 °C. SAXS measurements with filtered buffer were collected in parallel for background subtraction. 2D scattering data were radially averaged upon acquisition to give the measured scattering intensity $I(Q)$ as a function of $Q$ (Å$^{-1}$). SAXS profiles [$I(Q)$ vs. $Q$ (Å$^{-1}$)] from matched buffer conditions were subtracted from sample scattering profiles using SasView (Doucet, M., et al. SasView version 4.2.2. Available from: http://www.sasview.org). $Q$-values at the second maxima of the scattering profile were used to calculate the dominant headgroup to headgroup distance across the short dimension, $L$, of lipid-detergent assemblies according to the relationship $Q = 2p/L$.

**Laurdan generalized polarization.** Fluorescence for DMPC:CHAPS bicelles doped with a 1:400 molar ratio of Laurdan:amphiphile was measured using a Spectromax ID5 Plate Reader (Molecular Devices). Experiments were conducted in a 96-well costar black transparent bottom plate with a final sample volume of 200 µL. Laurdan was excited at 375 nm, and emissions were recorded from 400 nm to 550 nm in increments of 1 nm. Laurdan generalized polarization ($GP$) was calculated using the following equation (*5*):

$$GP = \frac{I_{440nm} - I_{490nm}}{I_{440nm} + I_{490nm}} \quad \textbf{Eq. 6}$$

**Calculation of effective *q*-values ($q_{\mathrm{eff}}$).** The actual lipid-to-detergent ratio in a bicelle was calculated based on the following equation (*6*):

$$q_{eff} = \frac{[lipid]_{bicelle}}{[detergent]_{bicelle}} = \frac{[lipid]_{total}}{[detergent]_{total} - CBC} \quad \textbf{Eq. 7}$$

, where $[\mathrm{detergent}]_{\mathrm{bicelle}}$ denotes the concentration of detergents that partition into the bicellar phase and *CBC* stands for the critical bicelle concentration denoting the free detergent concentration ($[\mathrm{detergent}]_{\mathrm{free}}$) that partitions into the aqueous phases. The $q_{\mathrm{eff}}$ was calculated in two ways. Primarily, the $[\mathrm{detergent}]_{\mathrm{free}}$ was fixed to $[\mathrm{DHPC}]_{\mathrm{free}}$ = 6.5 mM or $[\mathrm{CHAPS}]_{\mathrm{free}}$ = 2.5 mM as experimentally derived at high $q$ values ($q$ >0.75 for DMPC:DHPC and $q_{\mathrm{eff}}$ >1.0 for DMPC:CHAPS) (*6, 7*). The $q_{\mathrm{eff}}$ was also calculated under the assumption that lipids and detergents are ideally mixed (*8*) (**Eq. 8**; **fig. S10**):

$$\frac{1}{CBC} = \frac{\chi_{lipid}}{CMC_{lipid}} + \frac{\chi_{detergent}}{CMC_{detergent}} \quad \textbf{Eq. 8}$$

, where $c_{\text{lipid}}$ and $c_{\text{detergent}}$ represent the mol fractions of lipid and detergent, respectively. $CMC_{\text{lipid}}$ and $CMC_{\text{detergent}}$ denote the critical micelle concentrations of lipid and detergent, respectively (DMPC: 6 nM; DHPC: 15 mM; CHAPS: 6 mM from https://avantilipids.com/tech-support/physical-properties/cmcs).

**Statistical Analysis.** Chow's test (*9*) (**Fig. 3**) evaluates whether the true coefficients (i.e., the slopes) in two linear regressions on different subgroups (i.e., the datasets with different degrees of burial of mutated residues) are equal. It tests whether the independent variables (i.e., the mutation-induced stability changes in micelles) have different impacts (i.e., the mutation-induced stability changes in bicelles) on the different subgroups of the population. The *F*-statistic (*F*) in Chow's test is defined as $F = [(S_C - (S_1 + S_2))/k]/[(S_1 + S_2)/(N_1 + N_2 - 2\bullet k)]$, where $S_C$, $S_1$, and $S_2$ denote the sum of squared residuals for the combined subgroups, subgroup 1, and subgroup 2, respectively. $N_1$ and $N_2$ designate the number of points in subgroup 1 and subgroup 2, respectively. $k$ and $(N_1 + N_2 - 2\bullet k)$ indicate the first (the number of datasets, $k = 2$) and second degrees of freedom, respectively. The *p*-value is calculated using the *F*-statistic and the degrees of freedom, where $p = 1 - f.dist(F\text{-statistic}, k, N_1 + N_2 - 2\bullet k, \text{TRUE})$. On the 5% significance level, the null hypothesis cannot be rejected if $p>0.05$.

**Cooperativity profiling.** Specific residue interaction is perturbed by a single point mutation in the background of the double-biotin variants $95_N172_M$–$BtnPyr_2$ ($\Delta G^o_{N\text{-}D}{}^N$) or $172_M267_C$–$BtnPyr_2$ ($\Delta G^o_{N\text{-}D}{}^C$), which is set as 'WT'. Then, the stability change induced by the same mutation was measured by steric trapping for each WT background ($\Delta\Delta G^o_{N\text{-}D,WT\text{-}Mut}{}^N = \Delta G^o_{N\text{-}D,WT}{}^N - \Delta G^o_{N\text{-}D,Mut}{}^N$ or $\Delta\Delta G^o_{N\text{-}D,WT\text{-}Mut}{}^C = \Delta G^o_{N\text{-}D,WT}{}^C - \Delta G^o_{N\text{-}D,Mut}{}^C$). Then, the differential effect of the mutation on the stability of the two subdomains is quantified as follows (*1*):

$$\Delta\Delta\Delta G = \left[\Delta G^o_{N-D,WT}{}^N - \Delta G^o_{N-D,Mut}{}^N\right] - \left[\Delta G^o_{N-D,WT}{}^C - \Delta G^o_{N-D,Mut}{}^C\right]$$
$$= \Delta\Delta G^o_{N-D,WT-Mut}{}^N - \Delta\Delta G^o_{N-D,WT-Mut}{}^C \quad \textbf{Eq. 9}$$

We apply four cut-off values, $\Delta\Delta\Delta G = -2RT, -RT, RT$ and $2RT$ ($R$: gas constant and $T$: absolute temperature) to resolve the degree of cooperativity of each residue interaction. For a given $\Delta\Delta\Delta G$ value, we assign the cooperativity profile as follows. $+2RT<\Delta\Delta\Delta G$: highly localized in N-subdomain; $+RT<\Delta\Delta\Delta G\leq+2RT$: moderately localized in N-subdomain; $-RT\leq\Delta\Delta\Delta G\leq+RT$: cooperative; $-2RT\leq\Delta\Delta\Delta G\leq-RT$: moderately localized in C-subdomain; $\Delta\Delta\Delta G<-2RT$: highly localized in C-subdomain.

**MD simulation of GlpG.** MD simulation setups were based on the crystal structure of *E. coli* GlpG (PDB code: 2IC8) (*10*). The bicelle was approximated to a lipid bilayer composed of 315 DMPC molecules, which was constructed using the CHARMM-GUI membrane builder (*11*). Two micellar systems were built with 120 (DDM120) and 150 (DDM150) DDM molecules per micelle, modeled by symmetrically enclosing the TM domain of GlpG with DDM molecules (*12*). Each of the GlpG–bilayer and GlpG–micelle composite systems were immersed in the TIP3P water solvent, followed by charge neutralization and ionization with 150 mM NaCl. Each system was composed of >90,000 atoms in a 115×115×89 Å$^3$ box. Independently, we prepared DDM120 and DDM150 micelles without GlpG as controls. All inter- and intramolecular interactions were enumerated under the CHARMM36 force field (*13*). The nonbonding van der Waals and short-range electrostatic interactions were treated with a typical cutoff distance of 12 Å, while the long-range electrostatic contributions were evaluated with the particle-mesh Ewald method. All simulations were carried out using GROMACS software (*14*) parallelized in the GPU-accelerated IBM Power8 machine. Each system was first subject to 10,000 steps of conjugate gradient energy minimization to remove any unfavorable atomic crash with lipids and GlpG, which were restrained to preserve their conformation and relative positions. The systems were pre-equilibrated along six scheduled steps as gradually removing the external restraints until no constraints. The simulations were proceeded with a 2 fs timestep in the semi-isotropic isobaric and isothermal ensemble of 1 atm and 310 K, where the pressure and temperature were controlled by Parrinello-Rahman barostat and Nosé-Hoover thermostat, respectively. The pressure was decoupled between the *xy*-plane and the z-axis, so the membrane normal fluctuated independently from the isotropic lateral motions (*xy*-plane).

**Assessing the equilibration of protein and amphiphiles.** The equilibration of GlpG conformation was examined by calculating the RMSD's of all heavy atoms referenced to the crystal structure. Regarding the equilibration of amphiphile conformation, we assessed the time-autocorrelated RMSD ($\tau$) as a function of the time lag, $\tau$ by averaging over all lipid or detergent molecules in the bulk as follows:

$$RMSD(\tau) = \frac{1}{N_L} \sum_{i=1}^{N_L} < RMSD_i(t, t+\tau) >_t \quad \textbf{Eq. 10}$$

, where $N_L$: the number of the lipid or detergent molecules; $<RMSD_i\,(t,\ t+\tau)>_t$: the heavy-atom RMSD between the *i*-th amphiphile's conformations at the time *t* and $t+\tau$, averaged over all available time *t*'s. The bulk lipid molecules were selected from the ones not in contact with the protein over the analysis period while the bulk detergent molecules from the control micelles without GlpG. Prior to the RMSD calculation, the amphiphiles under comparison at each *t* and $t+\tau$ were structurally aligned with each other by transrotating the heavy-atom conformations.

For the solvation dynamics of interfacial amphiphiles on protein, we analyzed how fast the lipid or detergent would dissociate from the protein by measuring the residence time ($\tau_R$) from the autocorrelation function on time, $c(\tau)$ for the amphiphile heavy atom within 5 Å from GlpG as follows:

$$c(\tau) = \frac{1}{Nc} \sum_{i=1}^{Nc} < c_i(t, t+\tau) >_t \quad \textbf{Eq. 11}$$

, where $N_c$ is the number of contact events, and a single contact event is defined as a consecutive contact of an amphiphile with no non-contacting time gap longer than the amphiphile relaxation time measured above. The autocorrelation function at the time $\tau$ of the *i*-th contact event, $<c_i\,(t,\ t+\tau)>_t$ is defined by $<q_i(t)\bullet q_i(t+\tau)/q_i^2(t)>_t$ the normalized product of heavy atom contact numbers of an amphiphile in the *i*-th contact event, $q_i(t)$ and $q_i(t+\tau)$ at two time moments (*t* and $t+\tau$), averaged over the time *t*.

**Residence time of amphiphiles.** The $\tau_R$ was defined as the time when the amplitude of contact autocorrelation reached 1/*e* of the initial value without any assumption on the dissociation mechanisms based on **Eq. 11**. Dissociation dynamics of an amphiphile from GlpG was assessed as a whole or parts (i.e., head group and tail). The tail of DMPC was defined as the atoms in two aliphatic chains (C22–C214 and C32–C314) while that of DDM included all carbons in the dodecyl chain (C1–C12), thus leaving the rest as the head group. As a control, we assessed self-dissociation of an amphiphile from another amphiphile, where the pairs were selected from the molecules in contact with each other in the bulk. DMPC molecules were selected from ones of no explicit protein contact in the bilayer, and DDM molecules from the micelles without GlpG. In each system, the residence time for self-dissociation was served as a reference for evaluating the relative preference toward protein.

**Solvation free energy ($\Delta G^{o}_{Solv}$) of amphiphiles on GlpG.** The average residence time ($\tau_{R,P\cdot L}$) of amphiphiles interacting with the protein was computed by analyzing the time-autocorrelation function for each amphiphile contacting the transmembrane domain of GlpG. The analysis is extended over all peripheral area of the TM domain of GlpG, encapsulating the entire network of possible amphiphile-protein binding events:

$$\begin{aligned} P\cdot L_{(1)} &\xrightarrow{k_{off,P\cdot L_{(1)}}} P + L_{(1)} \\ P\cdot L_{(2)} &\xrightarrow{k_{off,P\cdot L_{(2)}}} P + L_{(2)} \\ &\vdots \\ P\cdot L_{(i)} &\xrightarrow{k_{off,P\cdot L_{(i)}}} P + L_{(i)} \\ &\vdots \\ P\cdot L_{(N_c)} &\xrightarrow{k_{off,P\cdot L_{(N_c)}}} P + L_{(N_c)} \end{aligned} \qquad \textbf{Eq. 12}$$

Here, $P\cdot L_{(i)}$ represents the amphiphile-protein complex at the site ($i$) on the protein, and $L_{(i)}$ represents the amphiphile at the same site. The corresponding dissociation rate constant, $k_{off,\,P\cdot L(i)}$, was obtained based on the relationship, $k_{off,P\cdot L(i)} = 1/(\tau_{R,P\cdot L(i)})$, where $\tau_{R,P\cdot L(i)}$ denotes the residence time of an amphiphile $L_{(i)}$ at site ($i$) obtained from the contact autocorrelation function for the amphiphile using **Eq. 11**. Meanwhile, the dissociation reactions of amphiphile-amphiphile complexes in the bulk was considered.

$$L\cdot L_{(j)} \xrightarrow{k_{off,L\cdot L_{(j)}}} L_{(j)} + L_{(j)} \qquad \textbf{Eq. 13}$$

Here, $L\cdot L_{(i)}$ denotes the $j$-th amphiphile-amphiphile complex found in the bulk where $j = 1, \ldots, N_{L\cdot L}$ with the corresponding dissociation constant, $k_{off,L\cdot L(i)} = 1/(\tau_{R,L\cdot L(i)})$, where $\tau_{R,L\cdot L(i)}$ is the residence time of the amphiphile–amphiphile complex. The averaged dissociation rate constant $k_{off,L\cdot L(i)}$ was obtained from the residence time $\tau_{R,L\cdot L}$, which was averaged over the contact correlation function analysis (**Eq. 11**) of all available amphiphile-amphiphile complexes.

Thus, the solvation free energy, $\Delta G^{o}_{Solv(i)}$ of an amphiphile at a specific site $i$ on GlpG can be obtained using the dissociation rate constants through the following relationship:

$$\Delta G^{0}_{Solv(i)} = -RT \ln \frac{k_{off,L\cdot L}}{k_{off,P\cdot L_{(i)}}} = -RT \ln \frac{\tau_{R,P\cdot L_{(i)}}}{\tau_{R,L\cdot L}} \qquad \textbf{Eq. 14}$$

However, our current interest lies in the interactions that a protein has with amphiphile molecules across all possible sites rather than at specific sites. Thus, the averaged solvation free energy $\Delta G^{o}_{Solv}$ is evaluated by incorporating all site-specific solvation free energies $\Delta G^{o}_{Solv(i)}$ as follows:

$$\Delta G^{0}_{Solv} = -RT \ln\left(\frac{1}{N_c} \sum_i e^{-\Delta G^{0}_{Solv(i)}/RT}\right) \quad \textbf{Eq. 15}$$

Here, $N_c$ is the number of potential interaction sites. This formulation accounts for the thermal fluctuations and the ensemble of possible amphiphile positions on the protein, resulting in a comprehensive description of $\Delta G^{o}_{Solv}$. The summation over different sites allows for the incorporation of the extensive network of interactions and the competition between amphiphile–amphiphile and amphiphile–protein associations, not merely the exchange at a single site. This is linked to the dissociation reactions of amphiphiles occurring at various sites on the protein through the following.

$$\begin{aligned}
\Delta G^{0}_{Solv} &= -RT \ln\left(\frac{1}{N_c} \sum_i e^{-\Delta G^{0}_{Solv(i)}/RT}\right) \\
&= -RT \ln\left(\frac{1}{N_c} \sum_i \frac{k_{off,L\cdot L}}{k_{off,P\cdot L_{(i)}}}\right) \\
&= -RT \ln\left(\frac{1}{N_c} \sum_i \frac{\tau_{R,P\cdot L_{(i)}}}{\tau_{R,L\cdot L}}\right) \\
&= -RT \ln\left(\frac{1}{\tau_{R,L\cdot L}} \frac{\sum_i \tau_{R,P\cdot L_{(i)}}}{N_c}\right) \\
&= -RT \ln \frac{\tau_{R,P\cdot L}}{\tau_{R,L\cdot L}} \\
&= -RT \ln \frac{k_{off,L\cdot L}}{k_{off,P\cdot L}} \\
&= -RT \ln K_{Solv}
\end{aligned} \quad \textbf{Eq. 16}$$

Notably, the average residence time of the amphiphiles, $\frac{\sum_i \tau_{R,P\cdot L_{(i)}}}{N_c}$ was obtained through the contact correlation analysis (**Eq. 11**) of the amphiphile-protein interactions. In summary, this analysis accounts for the residence time of all amphiphiles interacting with the protein ($\tau_{R,P\cdot L}$) and compares it with that of amphiphiles interacting with each other ($\tau_{R,L\cdot L}$), as expressed in the following equation:

$$\Delta G^{0}_{Assoc} = -RT \ln K_{Assoc} = -RT \ln \frac{k_{off,L\cdot L}}{k_{off,P\cdot L}} = -RT \ln \frac{\tau_{R,P\cdot L}}{\tau_{R,L\cdot L}} \quad \textbf{Eq. 17}$$

Thus, $\Delta G^{o}_{Solv}$ is not merely representative of an exchange energy at a single site but encapsulates the solvation free energy for the protein–amphiphile interaction at all potential interaction sites presented by the protein.

**Expression and purification of OmpLA.** We expressed and purified OmpLA following established protocols (*15, 16*). The HMS *E. coli* cells with OmpLA plasmid was inoculated in 500 ml LB media. At $OD_{600nm}$ = 1.0, the cells were induced with 100 mM IPTG. After 6 h expression, the cells were harvested and lysed in 30 mL lysis buffer (50 mM Tris, 40 mM EDTA, pH=8.0) by a Emusiflex pressure homogenizer. 35 mL Brij was added to 35 mL cell lysate before centrifuge. The pellet was collected after spinning and washed 3 times with wash buffer (10 mM Tris, 1mM EDTA, pH=8.0). The purified inclusion body was aliquoted and stored at –20 °C.

**Folding of OmpLA.** OmpLA was folded into 1,2-diundecanoyl-sn-glycero-3-phosphocholine (DC11PC) or 1,2-dilauroyl-sn-glycero-3-phosphocholine (DC12PC) vesicles following established protocols (*15, 16*). The inclusion body of OmpLA was dissolved in 8 M Guanidine-hydrochloride (GdnHCl) with 100 mM citrate buffer (pH = 3.8). The protein concentration was adjusted to 100 mM after centrifugation (0.1 mm) and filtering of the supernatant with unfolded OmpLA to remove any possible undissolved aggregates. The OmpLA stock was first diluted to 6 μM in 2.5 M GdnHCl with 1.4 mM sulfobetaine (SB)-14 using citrate buffer (100 mM citrate, pH = 3.8). The diluted OmpLA was further drop-wisely diluted to 2 μM in 1 M (folding direction) or 5 M (unfolding direction) GdnHCl, in the presence of 4 mM D12PC or DC11PC vesicles (prepared by extrusion in citrate buffer). The dilutions were performed on a heat plate while stirring at 400 rpm, 42 °C. After overnight incubation at 37 °C, the second dilution was finally diluted to a range of GdnHCl concentrations from 1 M to 5 M and incubated for another 40 h at 37 °C. The fraction of populations was determined by the intrinsic fluorescence of lipid-facing Trp residues.

**Fitting titration data.** At least three replicates of each lipid type were collected. The fluorescence data was normalized and averaged before fitting. The normalized fluorescence data was fitted to a three-state model as described previously (*15, 16*),

$$Y_{obs} = \frac{Y_N + Y_I K_{N-I} + Y_U K_{N-I} K_{I-U}}{1 + K_{N-I} + K_{N-I} K_{I-U}} \qquad \textbf{Eq. 18}$$

, where $Y_{obs}$ is the observable, which is the normalized fluorescence; $Y_N$, $Y_I$, and $Y_U$ are the baselines for the native (N), intermediate (I) and unfolded (U) states ($Y_X = I_X + S_X \cdot [D]$, where X = N, I, or U. $I_X$ and $S_X$ are the intercept and slope of the baseline, respectively. [D] is the concentration of the denaturant GdnHCl); $K_{N\text{-}I}$ and $K_{I\text{-}U}$ are the equilibrium constants of the N to I and I to U transitions, respectively. By expressing the free energy change of each transition as a linear function, the equilibrium constants can be written as,

$$K_{N-I} = \exp\left(\frac{-(\Delta G^{\circ}_{N-I,l,w} - m_{N-I}[D])}{RT}\right) \qquad \textbf{Eq. 19}$$

$$K_{I-U} = \exp\left(\frac{-(\Delta G^{\circ}_{I-U,l,w} - m_{I-U}[D])}{RT}\right) \qquad \textbf{Eq. 20}$$

, where $\Delta G^{o}_{N\text{-}I,l,w}$ and $\Delta G^{o}_{I\text{-}U,l,w}$ are the transition free energies in water; $R$ is the gas constant; $T$ is the temperature; $m_{N\text{-}I}$ and $m_{I\text{-}U}$ are the dependence of the transition energies on denaturant concentrations. For the titrations with DC12PC, we used fixed $m$ values previously determined by global fitting on WT and many variants, $m_{N\text{-}I}$ = 2.0 kcal·mol$^{-1}$·M$^{-1}$ and $m_{I\text{-}U}$ = 7.2 kcal·mol$^{-1}$·M$^{-1}$. For the titrations with DC11PC, we floated both $m$ values during fitting.

**MD simulations of OmpLA.** We performed all-atom MD simulation with the CHARMM36m force field on OmpLA using NAMD. We made the initiation files using CHARM-GUI with a similar setup as previously described (*15, 17-19*). Briefly, we built the system of OmpLA (PDB code: 1QD5) (*20*) embedded in DC12PC or DC11PC bilayer (~75 lipid molecules at each leaflet) with 0.1 M KCl at 1 atm, 37 °C. After the six initial equilibration steps, we continued to equilibrate the system for another 50 ns of production run. A total of at least 150 ns run was performed for both DC12PC and DC11PC systems. We collected the data from the last 100 ns simulations for analysis. The simulation trajectories of OmpLA and lipids were analyzed with MDAnalysis (*21*) and MOSAICS (*22*). Specifically, the "membrane protein tilt angle" and "average lipid conformation" tools were used from MOSAICS. For the average lipid conformation analysis, the membrane was projected onto a grid lattice around the protein. The coordinates of every lipid molecule that visited a certain lattice point were time-averaged over the trajectory. The resulting visualization included non-physical structures due to the averaging-out of the rotational dynamics of lipids, but it represented the overall position and shape of the lipids in the system.

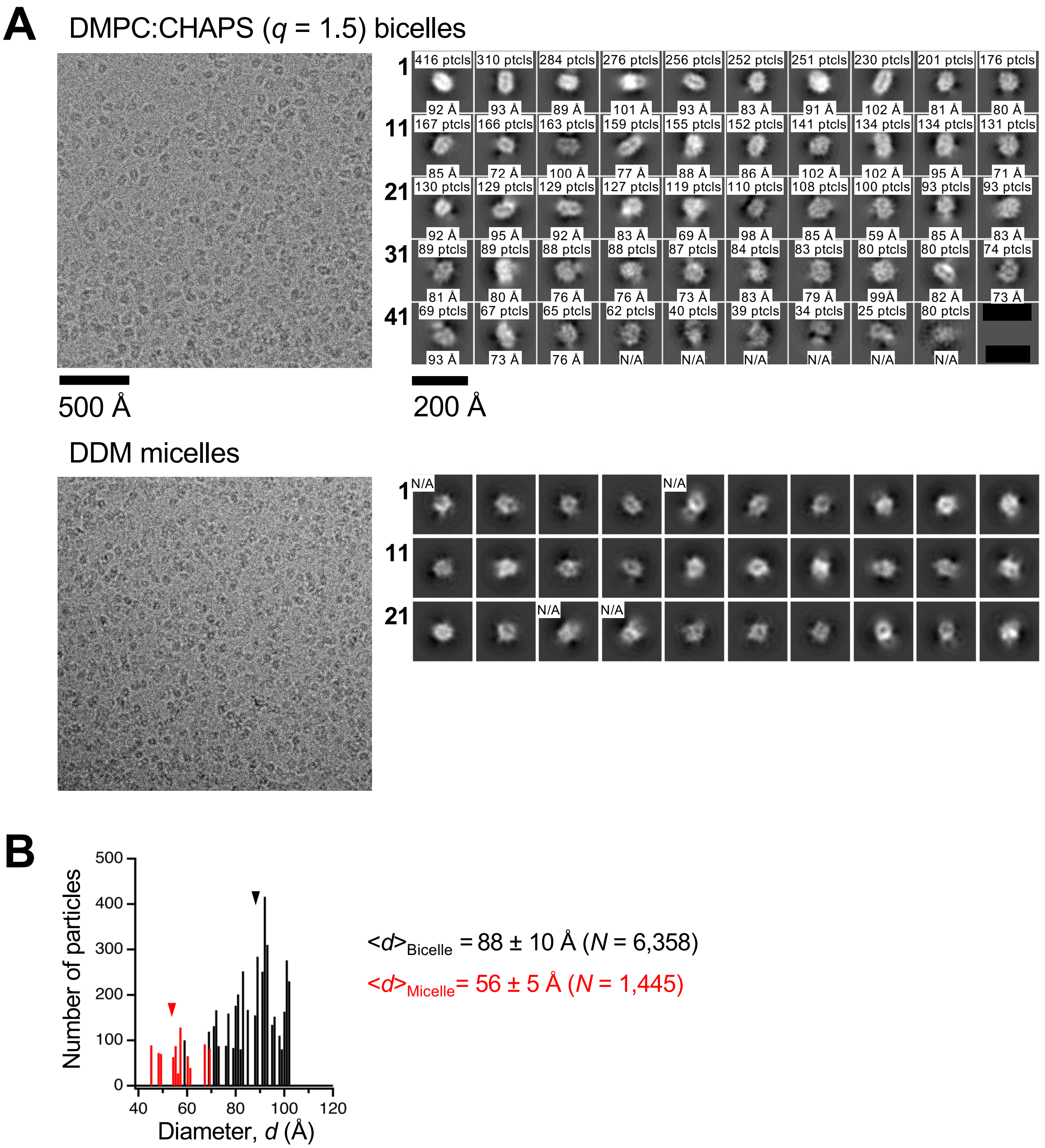


**Fig. S1.** Cryo-EM study of bicelles and micelles. **(A)** Selected raw images (*left*) and 2D class averages (*right*). For bicelles (*top*), a total of forty-nine 2D-class average images were obtained from $N$ = 6,358 particles. For micelles, a total of thirty 2D-class averages were obtained from $N$ = 1,445 particles. Bicelles and micelles were formed in 20 mM HEPES buffer (pH 7.5, 40 mM KCl, and 1 mM DTT). In each 2D class average panel, the maximal particle (ptcl) lengths were shown. The maximal particle length was not determined for the panels marked with asterisks due to the blurring of particle borders or the overlapping of multiple micelles. **(B)** Size analysis based on 2D class averages. Errors denote the average ± SD. The mean diameter values for micelles ($\langle d \rangle_{\text{micelles}}$) and bicelles ($\langle d \rangle_{\text{bicelles}}$) are marked as arrowheads.

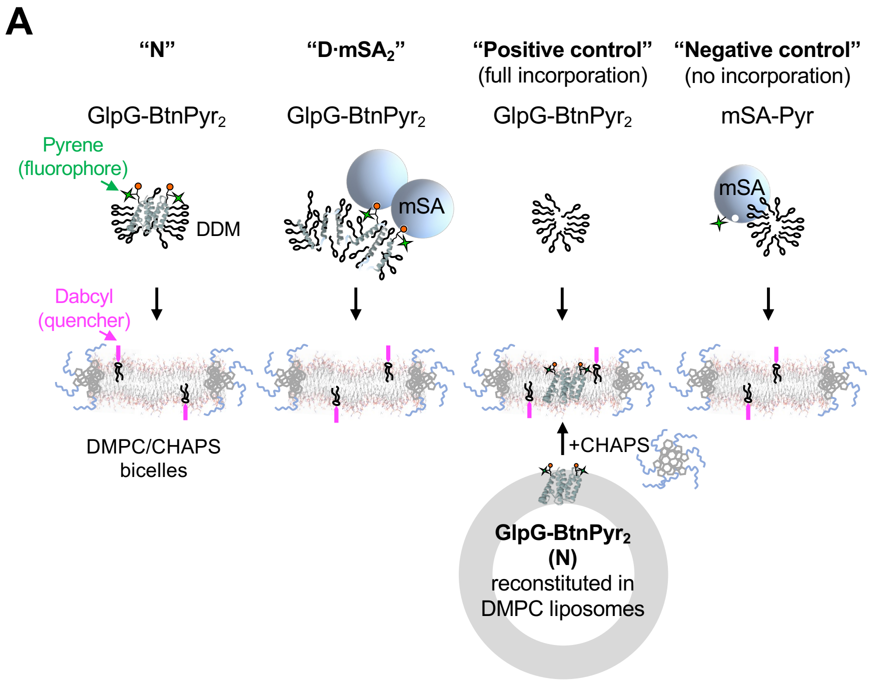


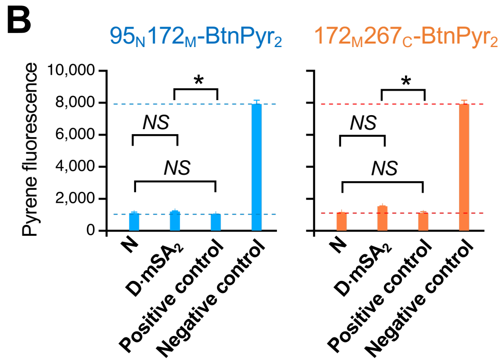


C

| Construct | Incorporation ratio (%) | |
|---|---|---|
| | **N** injected | **D·mSA₂** injected |
| 95N172M-BtnPyr2 | 99.1 ± 1.1 | 97.6 ± 0.6 |
| 172M267C-BtnPyr2 | 99.5 ± 1.3 | 93.7 ± 1.3 |

**Fig. S2. Incorporation of native and sterically denatured GlpG into bicelles. (A)** Schematic description of a fluorescence quenching assay to measure the incorporation of native ("N": the folded double-biotin variants, GlpG-BtnPyr$_2$) and sterically denatured GlpG ("D·mSA$_2$") in DDM micelles to bicelles (3 w/v-%). As a "positive control" representing full incorporation of the protein, GlpG-BtnPyr$_2$ was reconstituted in DMPC liposomes first and then the proteoliposomes were solubilized by CHAPS to form bicelles. As a "negative control" representing no incorporation, water-soluble mSA-Y83C variant labeled with thiol-reactive pyrene-maleimide was added to bicelles. In the samples for both "positive" and "negative" controls, DDM, whose final concentration is comparable to the "N" and "D·mSA$_2$" samples (0.01 w/v-% to 0.02 w/v-%), was added. **(B)** The assay result. Incorporation of GlpG-BtnPyr$_2$ to bicelles induced quenching of pyrene fluorescence. Error bars denote ± SEM. ($N = 3$). The $p$-values for the student $t$-test are shown (*NS*: $p$>0.05; *: $p$<0.05). (*c*) Incorporation ratios of native and sterically denatured GlpG from the micellar to the bicellar phase. The incorporation ratio = [$F$ ("negative control") – $F$ ("N")]/[$F$ ("negative control") – $F$ ("positive control")]. $F$ indicates the fluorescence intensity at 390 nm in each sample. Errors denote ± SEM. ($N = 3$).

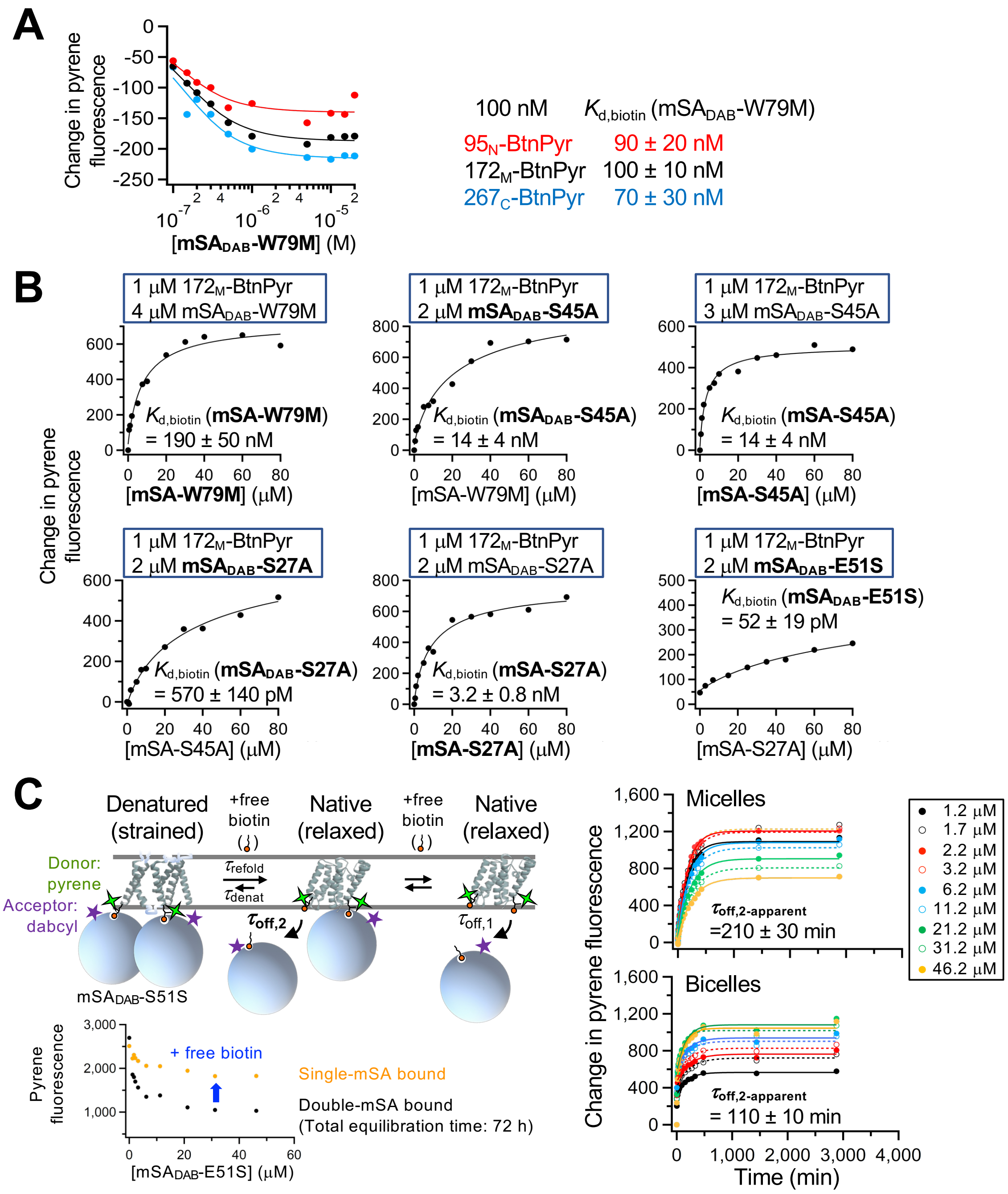


**Fig. S3. Characterization of binding of mSA variants to BtnPyr on GlpG. (A)** Binding isotherms between a weak binding variant mSA-W79M labeled with the dabcyl quencher ($mSA_{DAB}$-W79M) and GlpG variants with a single biotin label at three different positions. The data were fitted to **Eq. 1 (Methods)** to determine $K_{d,biotin}$. Errors denote ± SD from fitting. **(B)** Competition assay results to determine $K_{d,biotin}$ between high-affinity mSA variants and a single-biotin variant of GlpG ($172_M$-BtnPyr: BtnPyr conjugated to G172C in GlpG). The data were fitted to **Eq. 2**. Errors denote ± SD from fitting. **(C)** Measuring the lifetime of the second bound $mSA_{DAB}$-S51S ($\tau_{off,2}$) using dequenching of pyrene fluorescence. Errors denote ± SEM from fitting ($N = 9$).

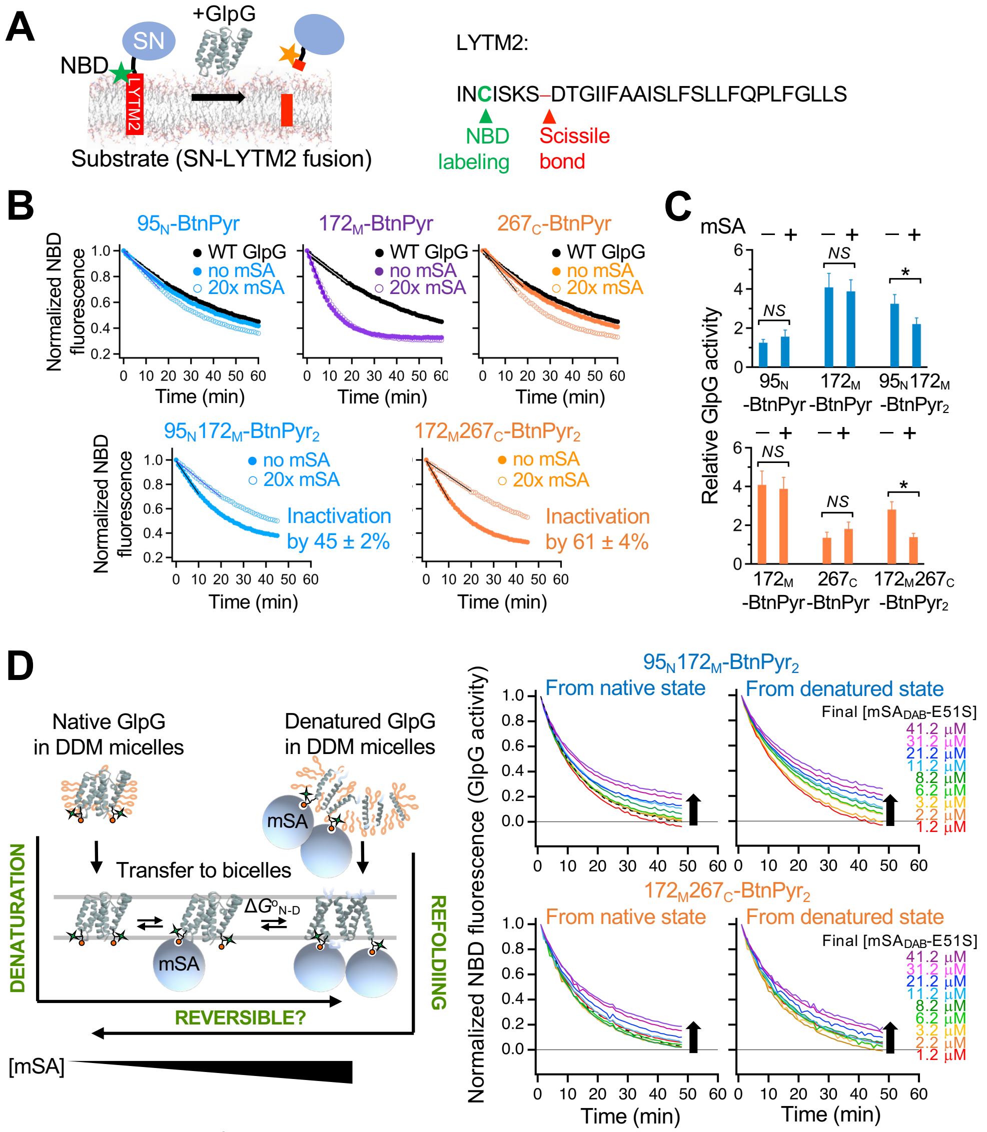


**Fig. S4. Activity assays to measure GlpG denaturation by steric trapping. (A)** Assay to measure the proteolytic activity of GlpG using the transmembrane model substrate LYTM2 (SN: staphylococcal nuclease fusion). The five-residue upstream residue (Cys) from the scissile bond was labeled with NBD fluorophore. The cleavage of LYTM2 by GlpG induces quenching of NBD fluorescence. **(B, C)** Inactivation (i.e., denaturation) occurs when binding of mSA to the double biotin variants is saturated, not when mSA binds to the individual single-biotin variants. The *p*-values were obtained from the student *t*-test (*NS*: $p>0.05$; *: $p<0.05$). The incomplete inactivation is due to the incomplete double labeling of biotin (**fig. S5C**). Errors denote ± SEM ($N = 3$). (**D**) (*Left*) Testing the reversibility of GlpG folding. Refolding and denaturation of GlpG were monitored by GlpG activity at an increasing concentration of $mSA_{DAB}$-S51S in bicelles. (*Right*) The assay results.

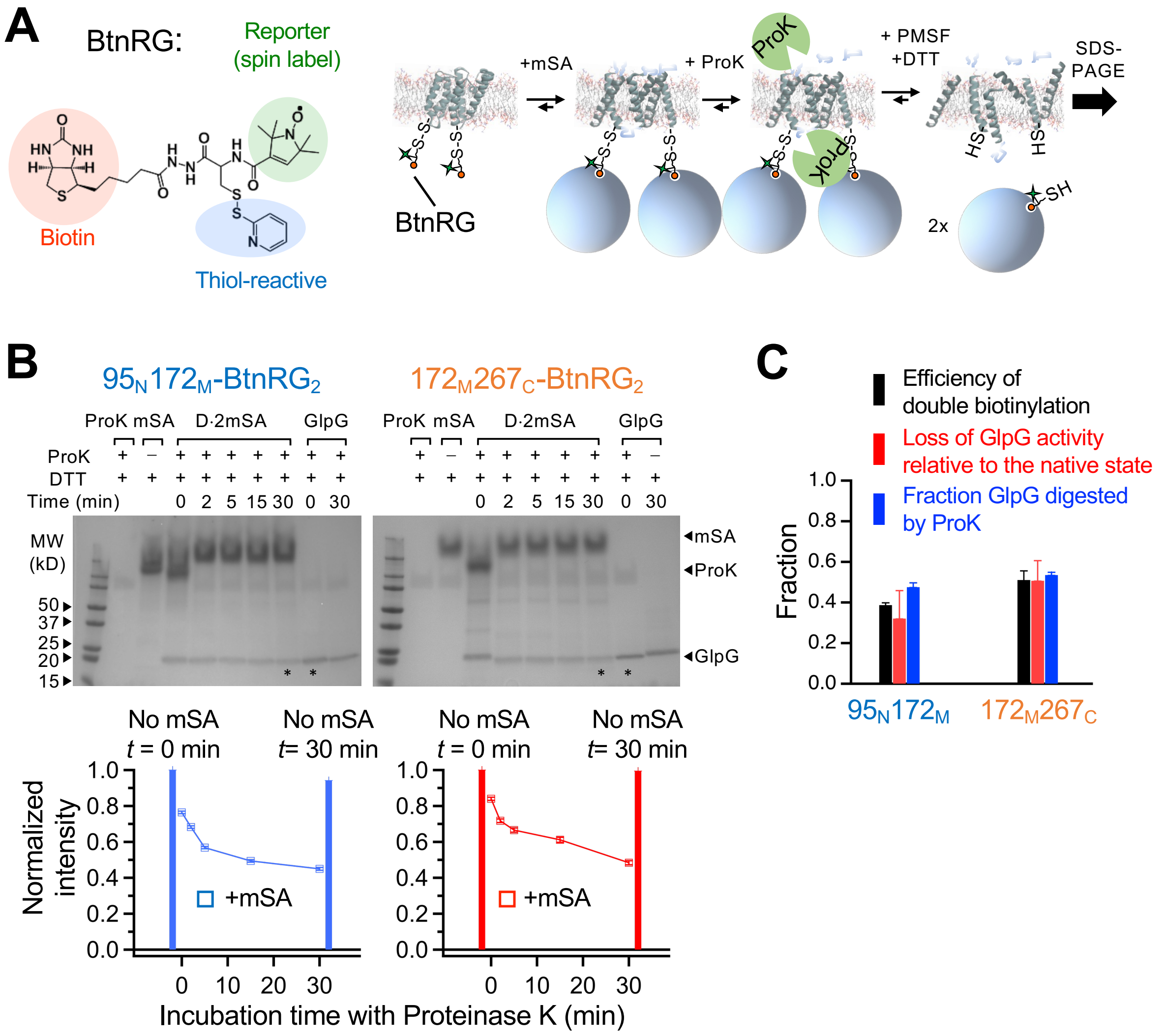


**Fig. S5. Denaturation of GlpG induced by steric trapping. (A)** (*Left*) Structure of the reversible thiol-reactive biotin label (BtnRG-TP). The thiopyridine group reacts with the thiol group in a cysteine residue to form a disulfide linkage. (*Right*) Strategy to detect sterically denatured GlpG by proteolysis. GlpG labeled with BtnRG is denatured by steric trapping. Denatured GlpG reacts with ProK for various incubation times. At each incubation time, the protease inhibitor PMSF and the reducing agent DTT are added to inactivate ProK and to break the linkage between GlpG and the biotin label bound with mSA, respectively. The final products are analyzed by SDS-PAGE. **(B)** (*Top*) Selective digestion of sterically denatured GlpG monitored by SDS-PAGE as a function of incubation time with ProK. (*Bottom*) The band intensities of GlpG on the SDS-PAGE gels were analyzed by the ImageJ program. As controls, the intensities of GlpG without mSA and with ProK at time 0 and 30 min are shown. **(C)** Correlation between the efficiency of double biotinylation, the activity loss induced by steric trapping, and the digestion by ProK. The incomplete digestion in the presence of excess mSA is due to the incomplete double-biotin labeling of GlpG. Error bars denote ± SEM ($N = 3$).

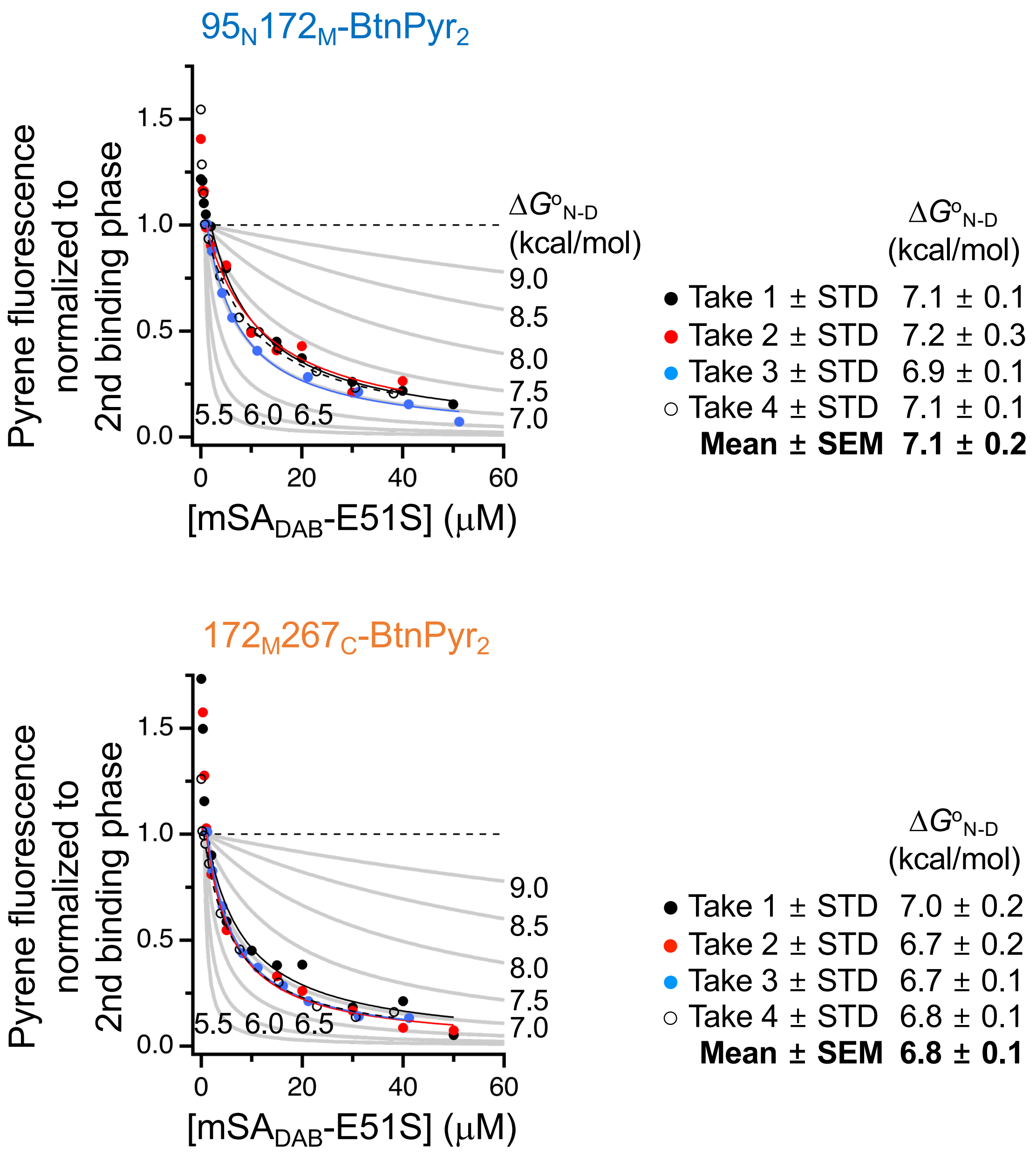


**Fig. S6. Precision of the stability measurements of GlpG using steric trapping.** Binding isotherms from four independent measurements (i.e., three different preparations of GlpG, mSA, and their labeled products) in 3 w/v-% DMPC:CHAPS bicelles ($q$ = 1.5) at 25 °C. The double-biotin variant of GlpG ($95_N172_M$–$BtnPyr_2$ or $172_M267_C$–$BtnPyr_2$) at 1 μM was titrated with an increasing concentration of $mSA_{DAB}$-E51S. Binding was monitored by pyrene fluorescence. In each isotherm, fluorescence intensities were normalized by the amplitude of the attenuated second binding phase.

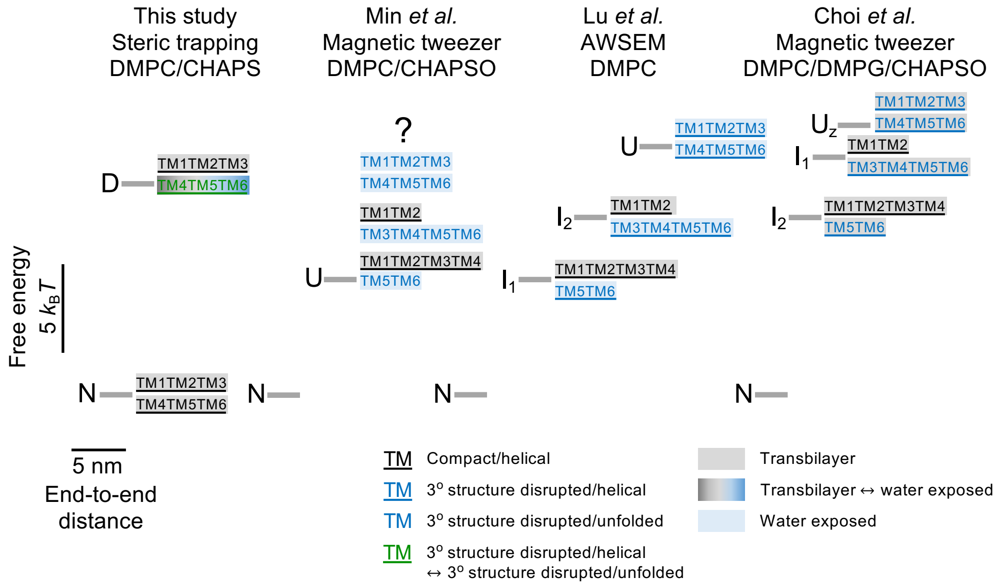


**Fig. S7. Comparison of the free energy landscapes of GlpG folding in the bilayer measured with various methods.** The positions of the free energy level (in $k_BT$) and the degree of compactness (the end-to-end distance between the N- and C-termini) of each state are scaled relative to the native state ("N"). The features of the conformation of each state are described regarding the tertiary interaction, ("compact" vs "3° structure disrupted"), the secondary structure ("helical" *vs* "unfolded"), or burial in the membrane ("transbilayer" *vs* "exposed to water"). "D": denatured state; "I": intermediate state; "U": unfolded state. The stability of GlpG that we determined directly under native condition ($\Delta G^o_{N\text{-}D}$ = ~–12$k_BT$, "This study") in bicelles is larger than that from the single-molecule magnetic tweezer study in the same neutral bicelles (–6.5$k_BT$, "Min *et al.*") (*23*). In the latter, $\Delta G^o_{N\text{-}D}$ is obtained by extrapolating the unfolding and refolding rates measured in two distinct force ranges (12 pN to 30 pN and 2 pN to 7 pN, respectively) to zero force. In the higher force range, GlpG unfolds via a single cooperative step or multiple steps with one or two intermediates to the fully stretched coil (*23*). In the lower force range, the conformation of the starting unfolded state prior to refolding is not defined ("?"). A computational study ("Lu et al. AWSEM") predicts that the unfolded state at low force is $I_1$ (TM1–TM4 folded) (*24*). Notably, our $\Delta G^o_{N\text{-}D}$ is similar to the free energy difference between the native state and $I_2$ (TM1–TM2 folded) from the same simulation (–10$k_BT$) (*24*) as well as to that between the native state and $I_1$ (TM1–TM2 folded) from the more recent tweezer study in the negatively charged bicelles (–13$k_BT$) at low force ("Choi et al.") (*25*). The end-to-end distances for "N" and "D" states in "This study" were taken from our previous work measured in the negatively charged bicelles (DMPC:DMPG:CHAPS) using DEER (*26*). In this study, we have shown that while both N- (TM1–TM3) and C- (TM4–TM6) subdomains in sterically denatured GlpG are expanded relative to the native state (*26*). The degree of compactness of N-subdomain from DEER is close to the collapse limit and that of C-subdomain is close to the full-expansion limit, resembling the conformations of $I_2$ from "Lu et al." and of $I_1$ from "Choi et al." (*24, 25*). Thus, we reason that the stability discrepancy is attributed to the different conformation of the denatured state in the steric trapping ("This study") vs magnetic tweezer ("Min et al.") studies.

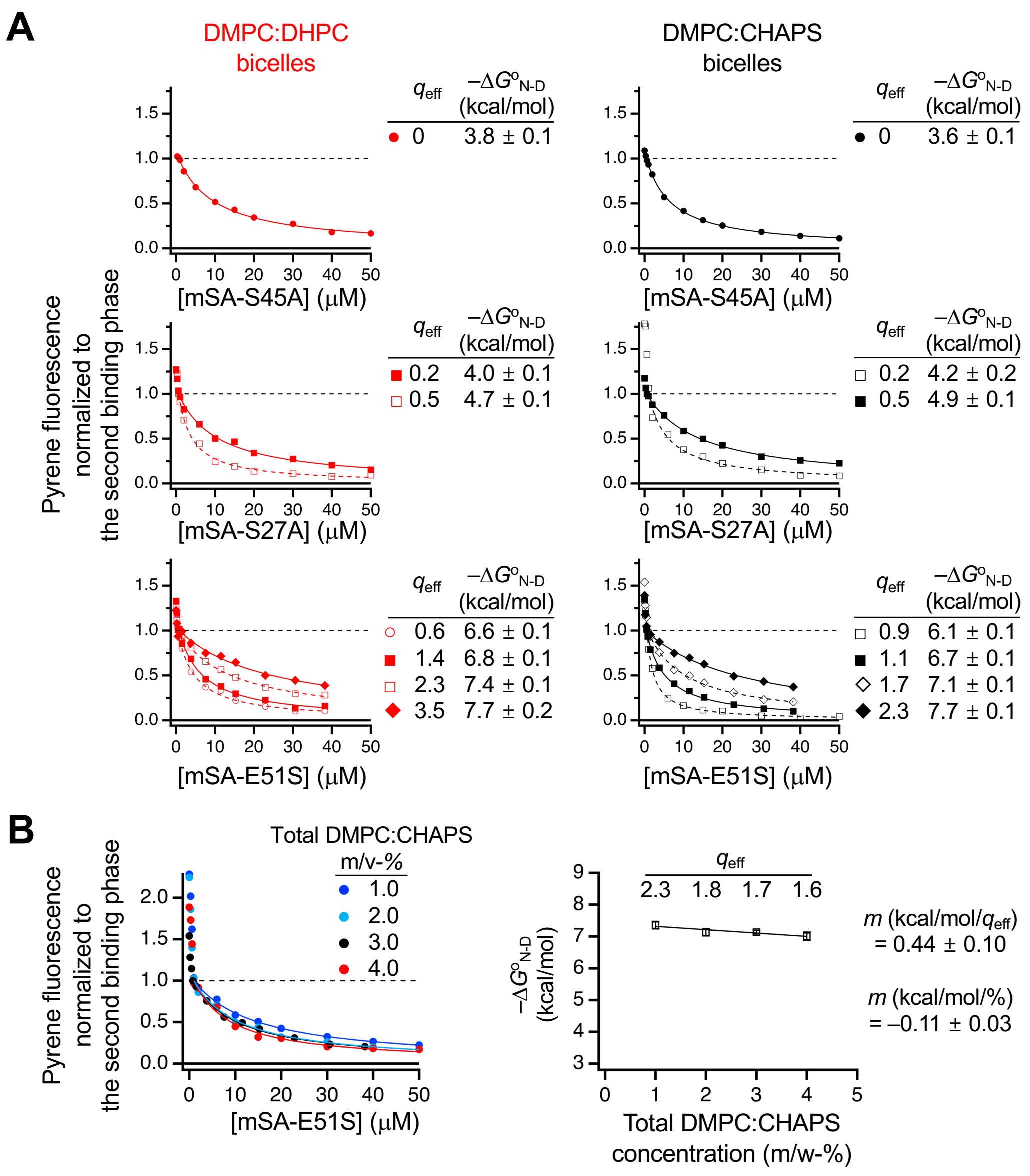


**Fig. S8. GlpG stability at an increasing lipid content in bicelles. (A)** Binding isotherm data for obtaining $\Delta G^o_{N\text{-}D}$ of GlpG ($95_N172_M$–$BtnPyr_2$) at various $q_{eff}$-values in the two types of bicelles (DMPC:DHPC-*left* and DMPC:CHAPS-*right*). The total amphiphilic concentration was fixed to 3 w/v-%. Errors denote ± SD from fitting. **(B)** (*Left*) Binding isotherms for obtaining $\Delta G^o_{D\text{-}N}$ of GlpG ($95_N172_M$–$BtnPyr_2$) at various w/v-% concentrations of the total amphiphiles (DMPC and CHAPS). The *q*-value ([DMPC]/[CHAPS]) was fixed to 1.5. (*Right*) The dependence of GlpG stability ($-\Delta G^o_{N\text{-}D}$) on the total amphiphile concentration (DMPC and CHAPS in w/v-%) at the fixed *q*-value (= 1.5). The $q_{eff}$ was calculated using the fixed free CHAPS concentration ($[CHAPS]_{free}$ = 2.2 mM) (**Eq. 7**). The "*m*" represents the slope of $-\Delta G^o_{N\text{-}D}$ as a function of $q_{eff}$ or w/v-% obtained from linear fitting. Errors denote ± SD from fitting.

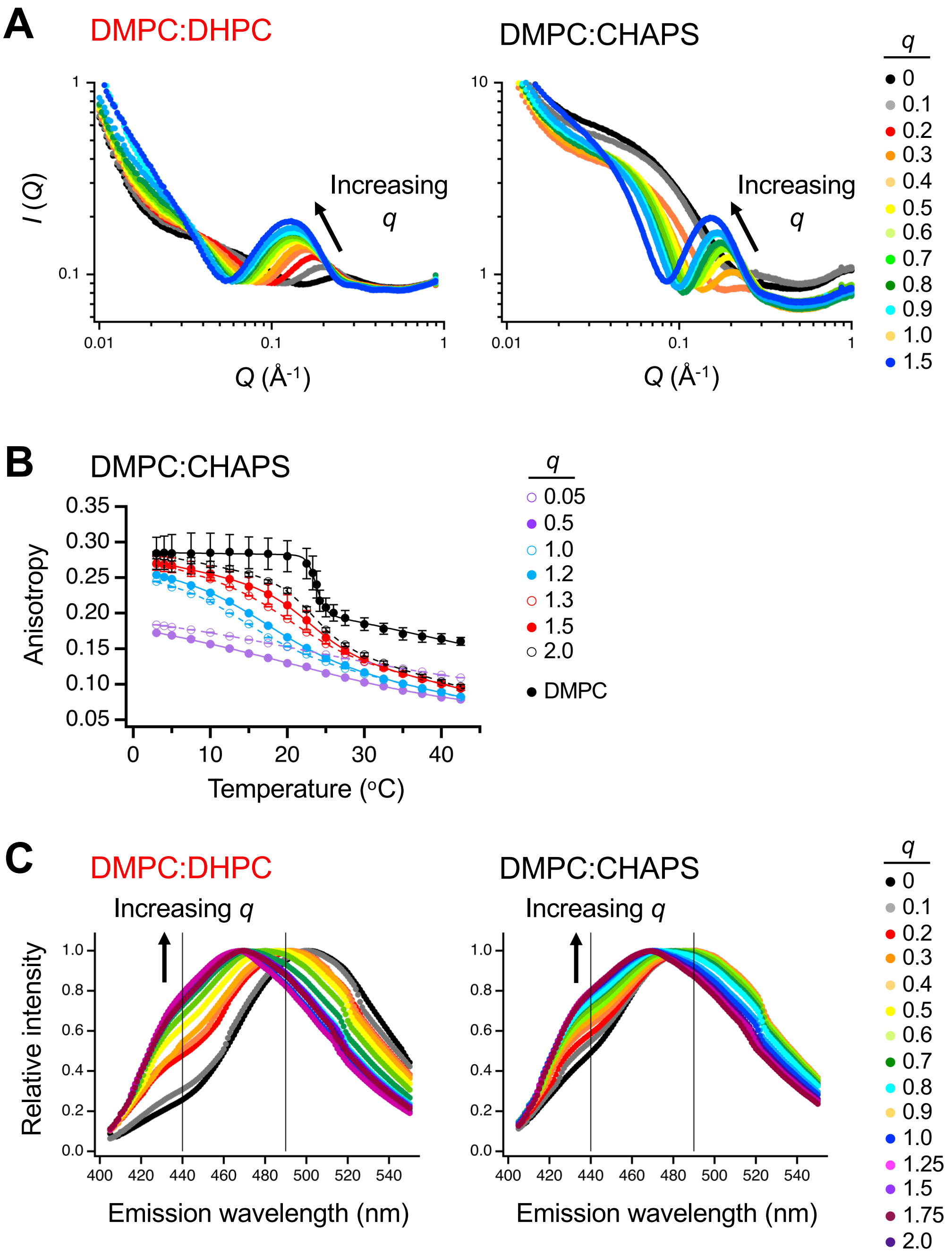


**Fig. S9. Characterization of bicelles using SAXS, fluorescence anisotropy, and Laurdan fluorescence. (A)** SAXS scattering profiles for DMPC:DHPC and DMPC:CHAPS bicelles. **(B)** Temperature-dependent fluorescence anisotropy of DPH incorporated into DMPC:CHAPS bicelles and large unilamellar DMPC liposomes. **(C)** Fluorescence spectra of Laurdan incorporated in DMPC:DHPC and DMPC:CHAPS bicelles. The vertical lines indicate the emission wavelengths of 440 nm and 490 nm, whose intensities are used for calculating generalized polarization (***GP*; Eq. 6**).

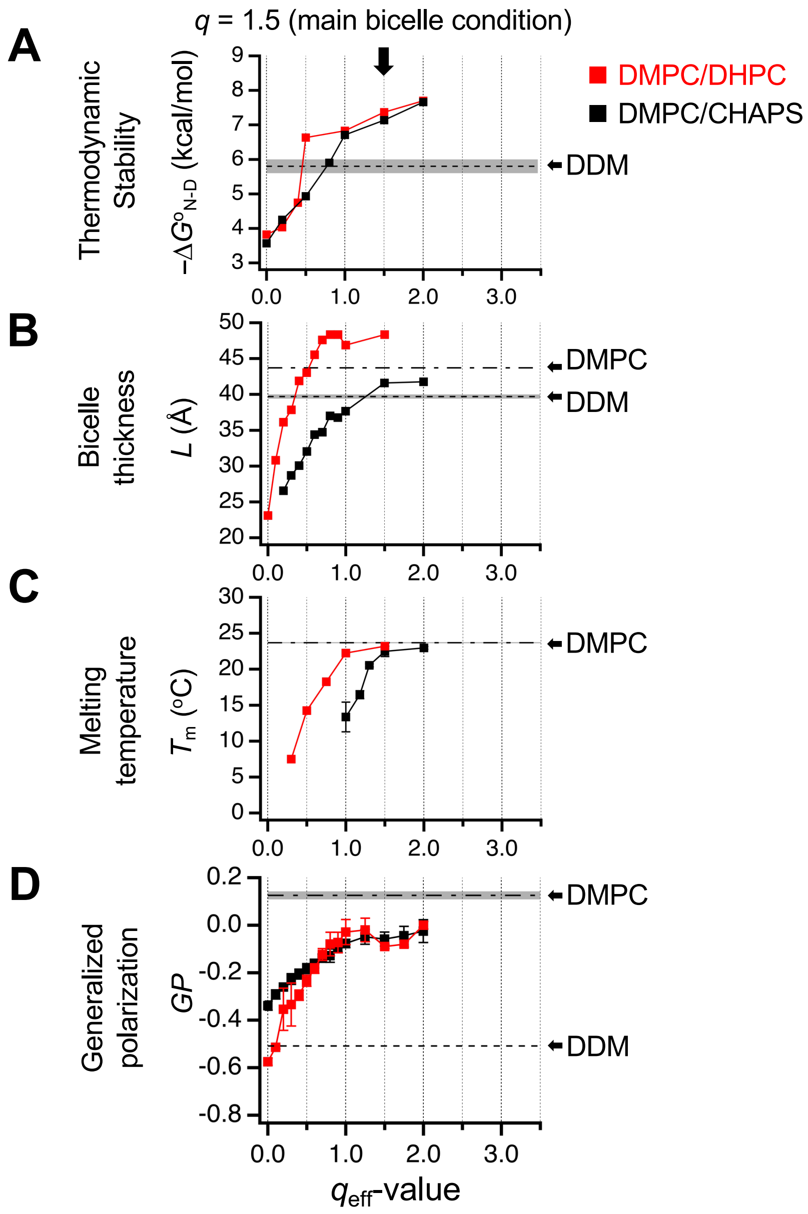


**Fig. S10. GlpG stability and physical parameters of bicelles measured as a function of the lipid content ($q_{eff}$) in DMPC:CHAPS and DMPC:DHPC bicelles.** Here, $q_{eff}$ was calculated using **Eq. 8** under the assumption that lipid and detergents are ideally mixed. **(A)** Thermodynamic stability ($-\Delta G^{o}_{N\text{-}D}$) of the double cysteine variant of GlpG, $95_{N}172_{M}$–BtnPyr$_2$. **(B)** The disk thickness (*L*) measured by SAXS. The thickness of DDM micelles was adapted from Ref. (*27*). **(C)** The gel–fluid phase transition temperature measured by fluorescence anisotropy of DPH incorporated in bicelles and DMPC liposomes. The data for DHPC:DMPC was adapted from Ref. (*28*). **(D)** Generalized polarization (*GP*) of Laurdan fluorescence for measuring the degree of hydration (i.e., the amphiphile–amphiphile packing strength).

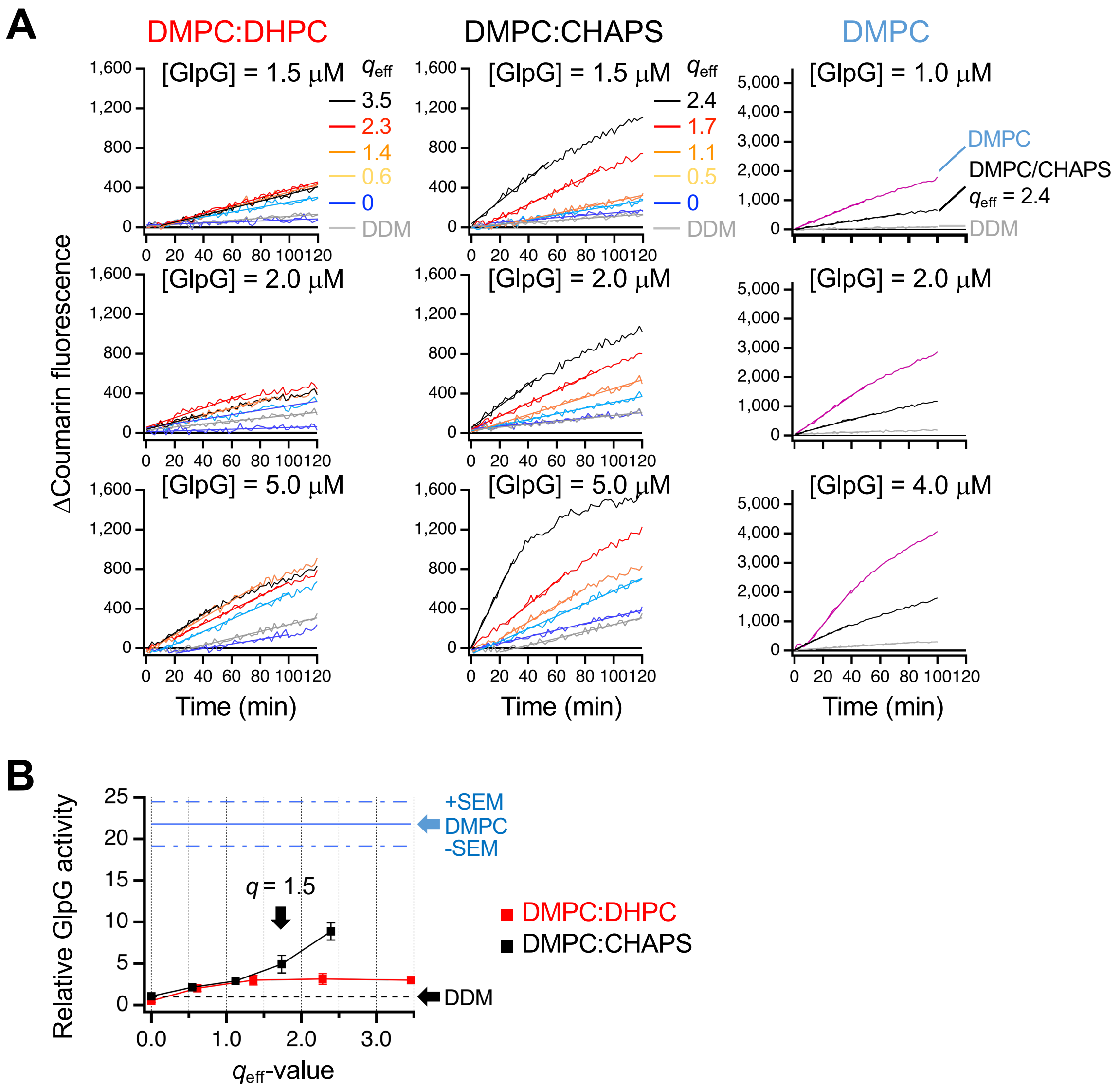


**Fig. S11. GlpG activity at an increasing lipid content in bicelles. (A)** Proteolytic activity of GlpG ($95_{N}172_{M}$–BtnPyr$_2$) measured as a function of effective $q$-value ($q_{eff}$) in two types of bicelles (DMPC:CHAPS and DMPC:DHPC; 3 w/v-%). An internally quenched water-soluble peptide, mca-RPKPYAv/WM-K(dnp), was used as a model substrate (mca: 7-methoxycoumarin; dnp: dinitrophenol; v: norvaline; "/": the scissile peptide bond). GlpG activity was also measured in DDM (3 w/v-%) micelles and DMPC (3 w/v-%) liposomes as references. **(B)** The activity in bicelles is represented by relative activity to that in DDM micelles. The $q_{eff}$-value of the main bicelles in this study ($q$ = 1.5) is marked with a block arrow. The relative activity of GlpG reconstituted in pure DMPC liposomes is shown as a horizontal reference line. Errors denote ± SEM ($N$ = 3).

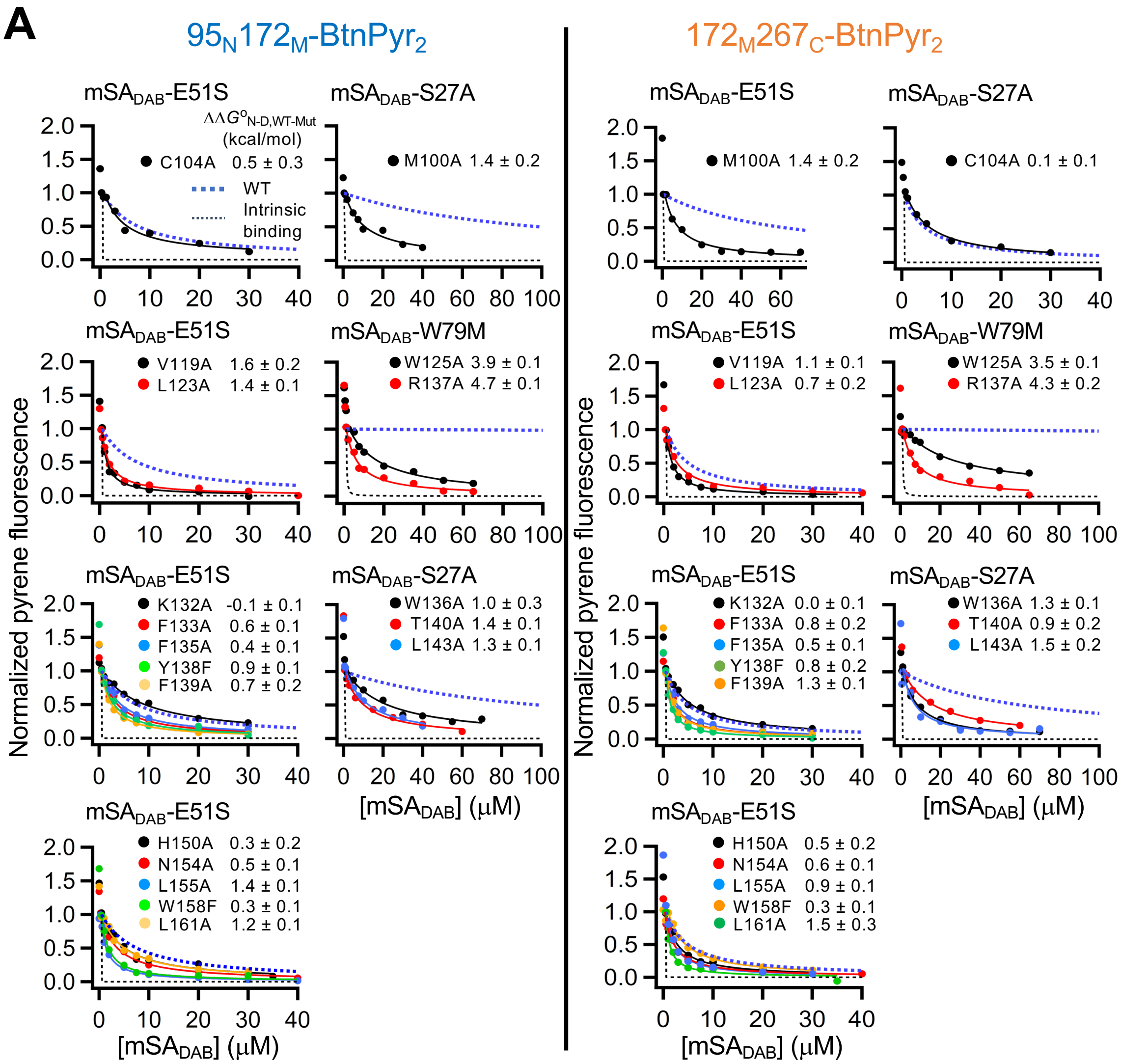


**Fig. S12. Binding isotherms between the double-biotin variant of GlpG and mSA to determine $\Delta G^{o}_{N\text{-}D}$ of GlpG in DMPC:CHAPS bicelles using steric trapping.** Binding was measured by quenching of pyrene fluorescence from the BtnPyr labels on GlpG by the dabcyl quencher conjugated to mSA ($mSA_{DAB}$). In each plot, the fluorescence intensity was normalized to the intensity change of the second binding phase. The data for WT GlpG and the predicted unhindered binding (i.e., the intrinsic binding of mSA to the biotin labels) are also shown as the blue and black dashed lines, respectively. The difference stability between WT and mutant ($\Delta\Delta G^{o}_{N\text{-}D,WT\text{-}Mut} = \Delta G^{o}_{N\text{-}D,WT} - \Delta G^{o}_{N\text{-}D,Mut}$) is shown. An mSA variant was chosen using the criteria that the attenuated second binding phase was observed up to 40 μM or 80 μM. *The more attenuated second binding indicates the higher stability.* **(A)** Binding isotherms for the variants bearing a mutation on the segments TM1, L1 and TM2 of GlpG. Errors denote ± SD from fitting. (Continued in the next page)

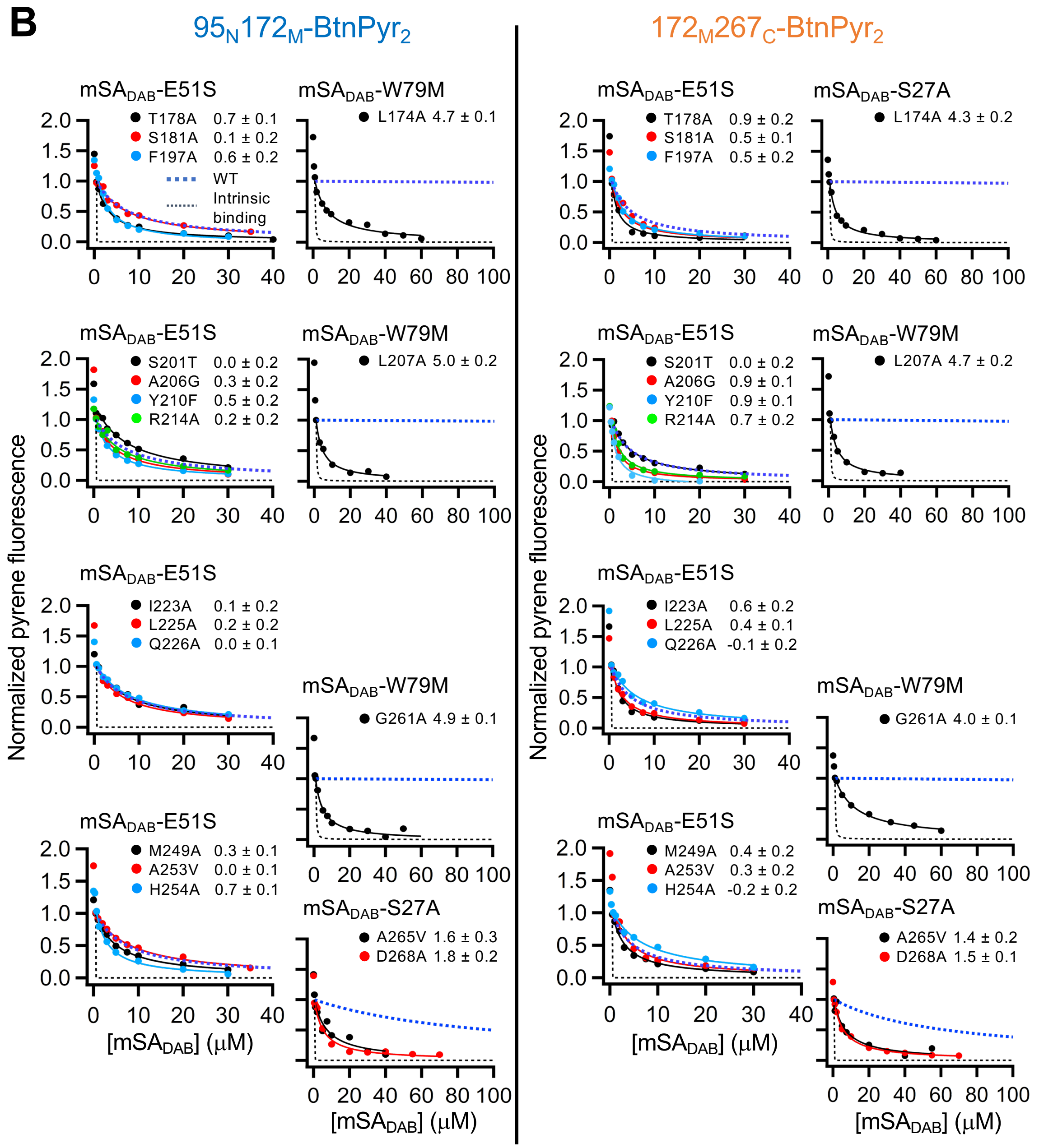


**Fig. S12.** (Continued from the previous page) **(B)** Binding isotherms for the variants bearing a mutation on the segments TM3, TM4, TM5 and TM6 of GlpG. Errors denote ± SD from fitting.

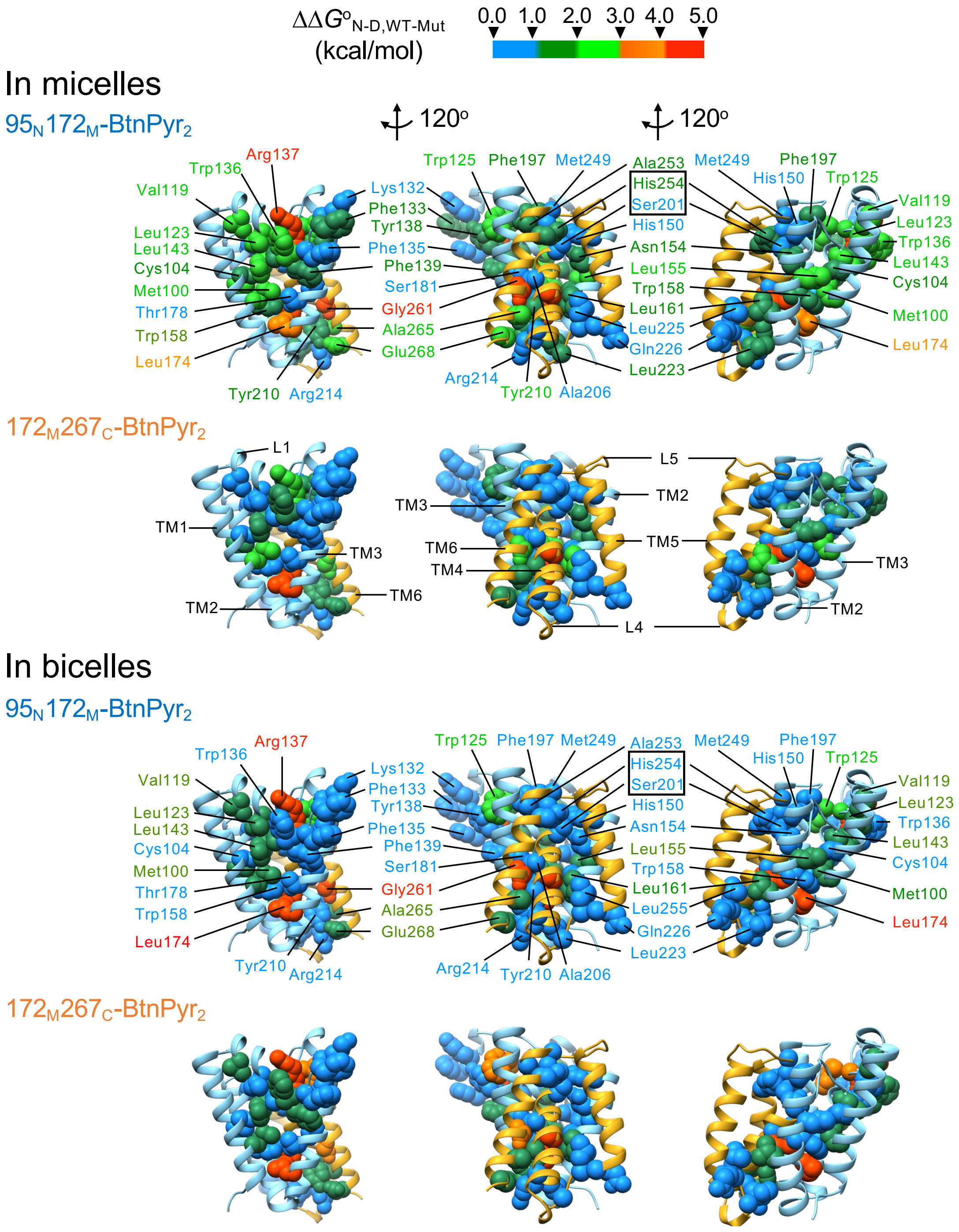


**Fig. S13. Mapping of the mutation-induced stability changes onto GlpG structure.** The mutation-induced stability changes ($\Delta\Delta G^{o}_{\text{N-D,WT-Mut}} = \Delta G^{o}_{\text{N-D,WT}} - \Delta G^{o}_{\text{N-D,Mut}}$) measured at N and C subdomains using the double biotin variants, $95_{N}172_{M}$-BtnPyr$_2$ and $172_{M}267_{C}$-BtnPyr$_2$, respectively, in micelles and bicelles were color-coded (*Top*) as a heat map on the structure.

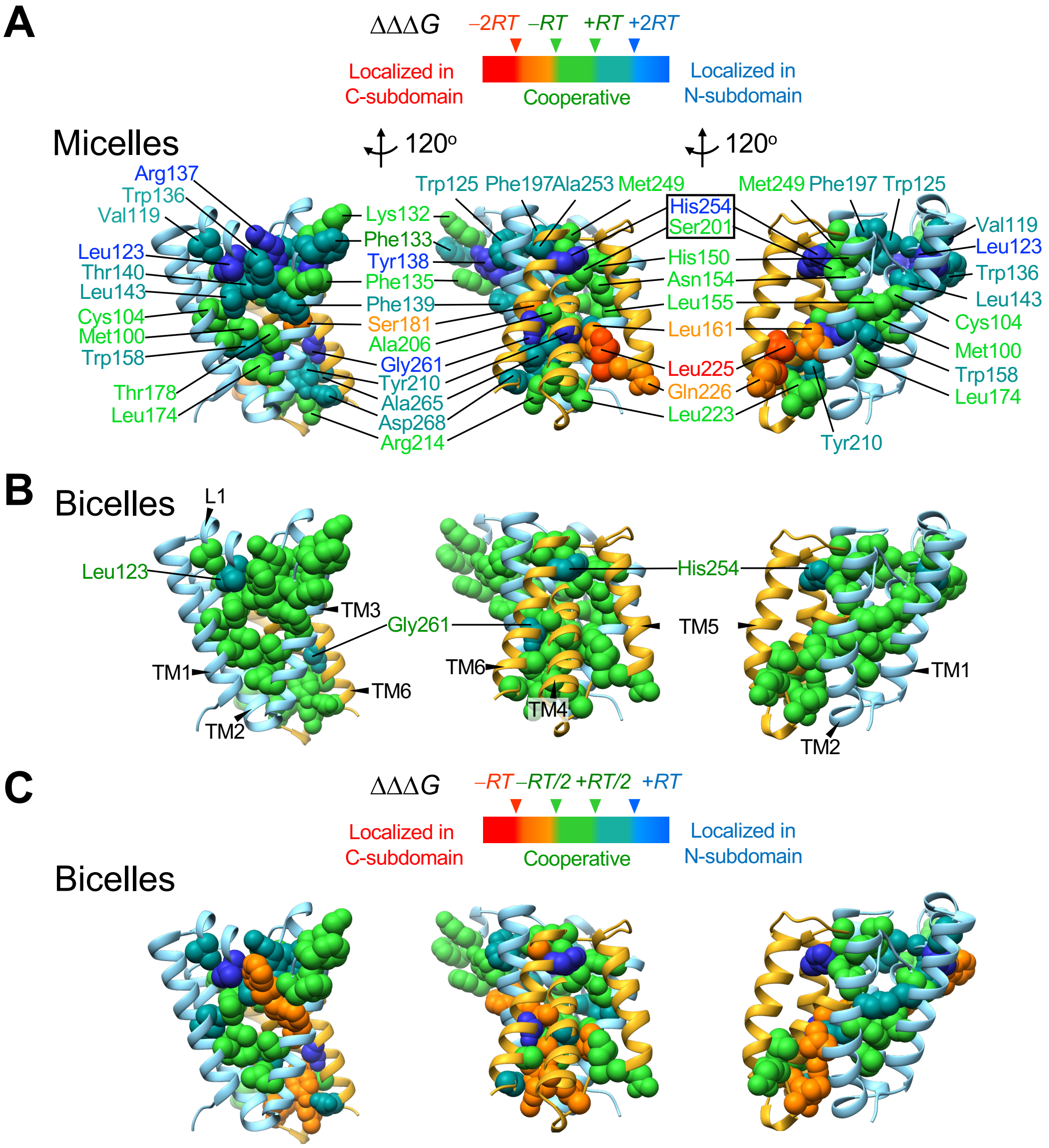


**Fig. S14. The features of cooperativity profiles in micelles are partially preserved in bicelles. (A, B)** Comparison of cooperativity profiles between micelles **(A)** and bicelles **(B)** based on the regular cut-off values, –2*RT*, –*RT*, +*RT*, and +2*RT* (i.e., the *RT* scale). **(C)** Cooperativity profiles in bicelles using the smaller cut-off values, –*RT*, –1/2*RT*, +1/2*RT*, and +*RT* (i.e., the 1/2*RT* scale). Except for several residues (Phe135, Phe136, Ala203, Ala206, Leu225, Gln226, and Arg214), the profiles reconstructed using the 1/2*RT* scale in bicelles are overall similar to those using the *RT* scale in micelles. The preserved features include: the cooperative packing core (formed by TM1, TM2 and TM3), the cooperative cluster in the active site (Ser201, His150 and Asn154), the localized cluster in L1, and the overpropagated cluster at the TM4-TM6 interface.

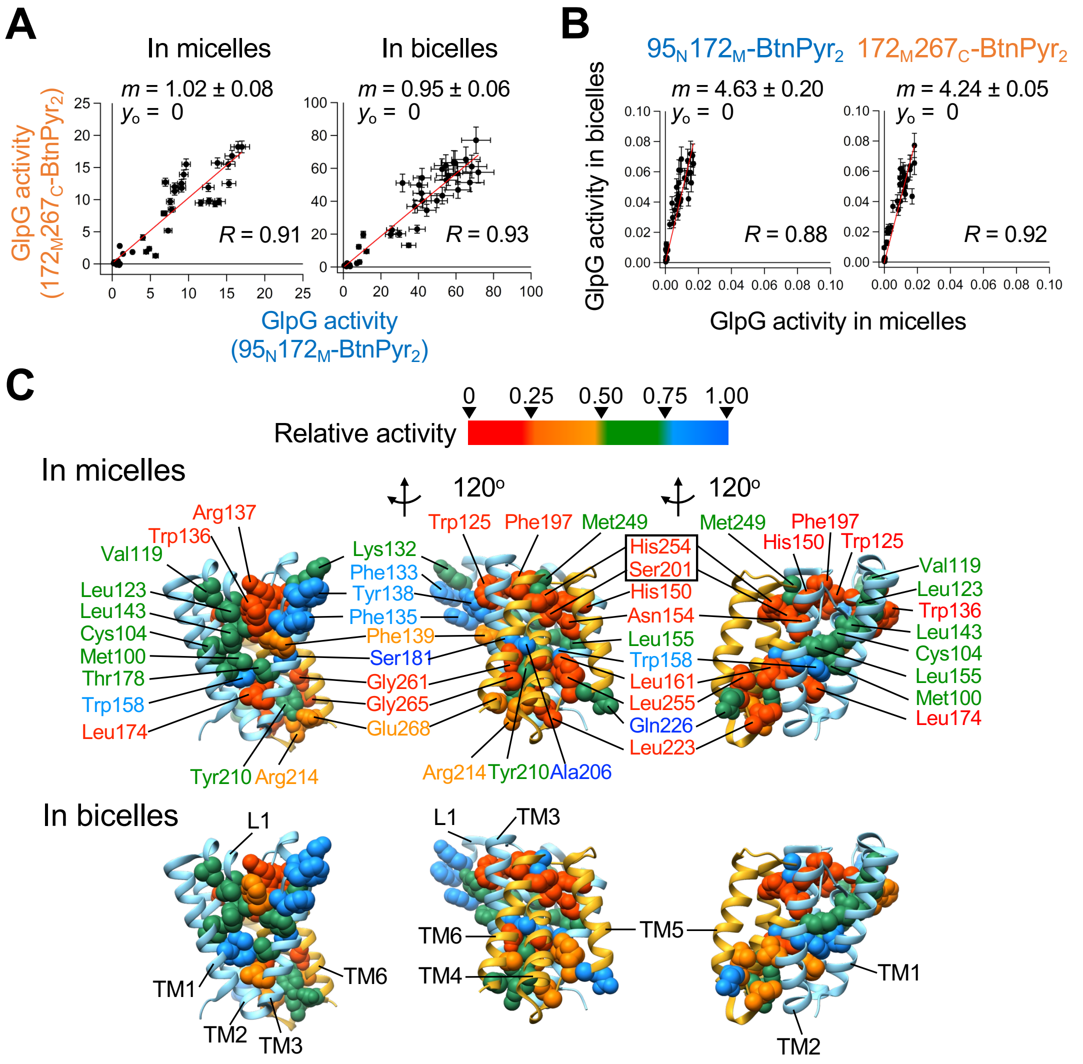


**Fig. S15. Proteolytic activity of GlpG WT and variants for the TM model substrate LYTM2. (A)** The effect of the location of the biotin pair ($95_N172_M$-BtnPyr$_2$ vs $172_M267_C$-BtnPyr$_2$) on GlpG activity. In both micelles and bicelles, the correlation slopes (*m*) are close to unity, indicating that the location of the biotin pair does not affect GlpG activity. Errors denote ± SEM ($N = 3$). **(B)** The effect of the hydrophobic environment (micelles vs bicelles) on GlpG activity. All activity values correspond to the fractional substrate turnover rate (min$^{-1}$) normalized to the initial substrate concentration (10 µM) in DDM micelles (5 mM) or DMPC:CHAPS bicelles (3 w/v-%, $q = 1.5$) as measured by NBD fluorescence (**fig. S4**). Errors denote ± SEM ($N = 3$). **(C)** Mapping of the mutation-induced activity changes onto the structure of GlpG. For each mutation, the activities measured in the background of $95_N172_M$-BtnPyr$_2$ and $172_M267_C$-BtnPyr$_2$ were normalized to the activity of WT ($95_N172_M$-BtnPyr$_2$ and $172_M267_C$-BtnPyr$_2$, respectively, without additional mutation), and then averaged for structural mapping.

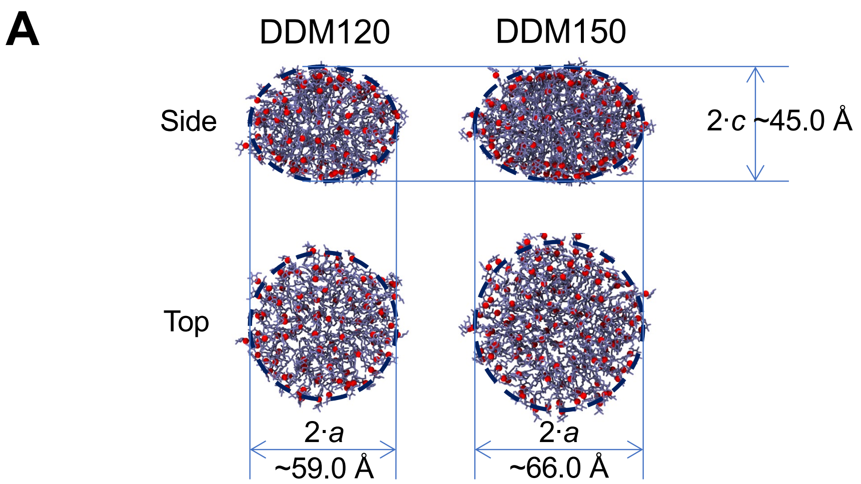


**B**

| Oblate structure | DDM120 | DDM150 | SAXS ($N_{A,DDM}$ = 135-149) |
|---|---|---|---|
| Radius, *a* | 29.5 ± 0.8 Å | 32.9 ± 0.8 Å | 33.7 ± 0.4 Å |
| Radius, *c* | 22.6 ± 1.1 Å | 22.5 ± 0.9 Å | 21.5 ± 0.4 Å |
| Oblateness | 0.23 ± 0.06 | 0.31 ± 0.04 | 0.36 ± 0.01 |
| Area per DDM | 77.3 ± 1.6 Å$^2$ | 72.1 ± 1.7 Å$^2$ | 76.3 ± 4.0 Å$^2$ |

**Fig. S16. Modelling of the micelles for MD simulation. (A)** Two micellar systems in this study, one with 120 DDM molecules (DDM120) and the other with 150 DDM molecules (DDM150). The shapes of both micellar systems were oblate spheroids with the axial dimension (2·*c*), which remained constant at $c$ = 22.5 Å. The shape of micelles was assessed for DDM120 and DDM150 without protein to understand the overall packing of DDM molecules. The structure of a DDM micelle was approximated by a spheroid as below:

$$\frac{x^2}{a^2}+\frac{y^2}{a^2}+\frac{z^2}{c^2}=1$$

, where $x$, $y$, and $z$ are the cartesian coordinates in the 3D-space. The semi-axes $a$ and $c$ are aligned along each symmetry axis, each indicating the equatorial radius in the $xy$-plane and the distance from the spheroid center to the pole along the symmetry axis of $z$, where $a>c$ forms an oblate spheroid, while $a<c$ a prolate. All coordinates of DDM 2O4 atoms in a micelle were utilized to describe the spheroidal shape of the micelle, which then were subjected to a parametric fitting for obtaining the semi-axes, $a$ and $c$. We found that both DDM120 and DDM150 create oblate spheroidal shapes (i.e., $a>c$), from which the effective cross-sectional area per DDM molecule was evaluated by using the equation, $A_{\text{oblate}} = 2pa^2 + pc^2/e \cdot \ln[(1+e)/(1-e)]$ (the eccentricity, $e$, is defined by $e = [1 - c^2/a^2]^{1/2}$) (*12*). **(B)** As the number of DDM molecules increases from 120 to 150, the equatorial dimension increases from $a$ = 29.5 Å to 32.9 Å. The area per DDM at the micellar surface is larger in DDM120 providing room for each DDM molecule to relax fast in the micelles relative to that in DDM150. The experimental values obtained from small-angle X-ray scattering (SAXS) are also shown for comparison (*27*). $N_{\text{A,DDM}}$: the aggregation number of DDM. In the SAXS data, the $a$ and $c$ values are obtained by adding the core radius and the shell thickness based on the two-component core-shell models (*27*).

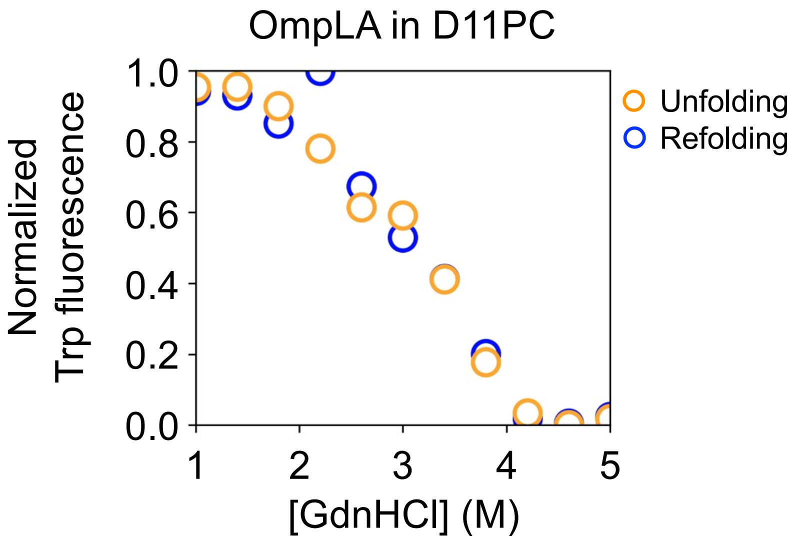


**Fig. S17. The unfolding and folding titrations of OmpLA show no hysteresis indicating the reversibility of folding.**

**Table S1. Fitted parameters of DEER data for native and sterically denatured GlpG in bicelles and micelles.** The background-subtracted dipolar evolution data were fit with the model-free, non-negative regularization. $r_{Prob}$: the most probable distance; $r_{Med}$: the median distance; $r_{Mean}$: the mean distance; $\sigma_{Mean}$: the standard deviation of the mean. $\chi^2$ indicates the goodness of fit. Modulation depth, which ranges from 0 to 1, represents the fraction of interacting spin pairs that lie within the detectable distance limit of the experiment. Distance limit is the interspin distance below which modeled distances can be estimated with confidence, for a given DEER experiment. Different DEER experimental parameters, often dictated by sample quality and amount, result in different distance limits.

| Bicelles (DMPC:CHAPS, $q$ = 1.5) | | | | | | | |
|---|---|---|---|---|---|---|---|
| | $r_{Prob}$ (Å) | $r_{Median}$ (Å) | $r_{Mean}$ (Å) | $\sigma_{Mean}$ (Å) | $\chi^2$ | Modulation depth | Distance limit (Å) |
| $95_N172_M$ | 24.4 | 28.2 | 32.7 | 12.2 | 1.15 | 0.352 | 59 |
| $95_N172_M \cdot mSA_2$ | 56.9 | 42.4 | 37.5 | 14.9 | 1.30 | 0.063 | 57 |
| $172_M267_C$ | 24.4 | 29.4 | 33.8 | 14.5 | 1.18 | 0.250 | 60 |
| $172_M267_C \cdot mSA_2$ | 55.6 | 48.4 | 41.2 | 14.9 | 1.31 | 0.208 | 60 |

| Micelles (DDM)* | | | | | | | |
|---|---|---|---|---|---|---|---|
| | $r_{Prob}$ (Å) | $r_{Median}$ (Å) | $r_{Mean}$ (Å) | $\sigma_{Mean}$ (Å) | $\chi^2$ | Modulation depth | Distance limit (Å) |
| $95_N172_M$ | 26.5 | 27.2 | 28.7 | 5.9 | 1.45 | 0.015 | 54 |
| $95_N172_M \cdot mSA_2$ | 53.4 | 47.4 | 43.6 | 12.4 | 1.43 | 0.008 | 54 |
| $172_M267_C$ | 27.8 | 25.2 | 25.2 | 4.1 | 3.19 | 0.032 | 54 |
| $172_M267_C \cdot mSA_2$ | 53.0 | 52.6 | 50.9 | 7.5 | 1.72 | 0.019 | 54 |

*The data in micelles was adapted from the previous publication (*26*).

**Table S2. The mutation-induced changes in thermodynamic stability ($\Delta\Delta G^{o}_{WT\text{-}Mut}$) and the activities relative to wild type in DMPC:CHAPS bicelles.** $f_{ASA}$: the fraction of residue buried area. In the "Location" column, "N" and "C": N- and C-subdomains, respectively. In the "Cooperativity profile" column, "Moderate/N", "Moderate/C", and "Moderate/Over": moderately localized in N and C subdomains, and moderately overpropagated, respectively. Errors denote ± SD from fitting ($N$ = 1–3). The $\Delta\Delta G^{o}_{WT\text{-}Mut}$ and $\Delta\Delta\Delta G^{o}$ values are in kcal/mol.

| Secondary structure | Mutation | Location | $f_{ASA}$ | N-subdomain ($95_N172_M$) | | C-subdomain ($172_M267_C$) | | $\Delta\Delta\Delta G^{o}$ | Cooperativity profile |
|---|---|---|---|---|---|---|---|---|---|
| | | | | $\Delta\Delta G^{o}_{WT\text{-}Mut}$ | Rel.Activity | $\Delta\Delta G^{o}_{WT\text{-}Mut}$ | Rel.Activity | | |
| TM1 | M100A | N | 0.14 | 1.4 ± 0.2 | 0.86 ± 0.09 | 1.4 ± 0.2 | 0.74 ± 0.08 | 0.0 ± 0.2 | Cooperative |
| | C104A | N | 0 | 0.5 ± 0.3 | 0.84 ± 0.09 | 0.1 ± 0.4 | 0.58 ± 0.06 | 0.4 ± 0.4 | Cooperative |
| L1 | V119A | N | 0.05 | 1.6 ± 0.2 | 0.59 ± 0.07 | 1.1 ± 0.1 | 0.70 ± 0.07 | 0.5 ± 0.2 | Cooperative |
| | L123A | N | 0.23 | 1.4 ± 0.1 | 0.57 ± 0.06 | 0.7 ± 0.2 | 0.65 ± 0.07 | 0.7 ± 0.2 | Moderate/N |
| | W125A | N | 0.09 | 3.9 ± 0.1 | 0.17 ± 0.02 | 3.5 ± 0.1 | 0.12 ± 0.01 | 0.5 ± 0.1 | Cooperative |
| | K132A | N | 0.56 | -0.1 ± 0.1 | 0.79 ± 0.09 | 0.0 ± 0.1 | 0.72 ± 0.08 | -0.1 ± 0.2 | Cooperative |
| | F133A | N | 0.80 | 0.6 ± 0.1 | 0.83 ± 0.10 | 0.8 ± 0.2 | 0.82 ± 0.09 | -0.2 ± 0.2 | Cooperative |
| | F135A | N | 0.73 | 0.4 ± 0.1 | 1.02 ± 0.12 | 0.5 ± 0.1 | 0.75 ± 0.09 | -0.1 ± 0.2 | Cooperative |
| | W136A | N | 0.46 | 1.0 ± 0.3 | 0.22 ± 0.02 | 1.3 ± 0.1 | 0.30 ± 0.03 | -0.3 ± 0.3 | Cooperative |
| | R137A | N | 0.04 | 4.7 ± 0.1 | 0.01 ± 0.01 | 4.3 ± 0.2 | 0.01 ± 0.01 | 0.4 ± 0.2 | Cooperative |
| | Y138F | N | 0.25 | 0.9 ± 0.1 | 0.75 ± 0.08 | 0.8 ± 0.2 | 0.56 ± 0.07 | 0.1 ± 0.2 | Cooperative |
| | F139A | N | 0.38 | 0.7 ± 0.2 | 0.63 ± 0.07 | 1.3 ± 0.1 | 0.45 ± 0.05 | 0.5 ± 0.2 | Cooperative |
| | T140A | N | 0.20 | 1.4 ± 0.1 | 0.59 ± 0.07 | 0.9 ± 0.2 | 0.62 ± 0.07 | 0.5 ± 0.2 | Cooperative |
| | L143A | N | 0.25 | 1.3 ± 0.1 | 0.78 ± 0.09 | 1.5 ± 0.2 | 0.68 ± 0.08 | -0.2 ± 0.2 | Cooperative |
| TM2 | H150A | N | 0.01 | 0.3 ± 0.2 | 0.10 ± 0.01 | 0.5 ± 0.2 | 0.03 ± 0.00 | -0.2 ± 0.3 | Cooperative |
| | N154A | N/Interface | 0 | 0.5 ± 0.1 | 0.12 ± 0.01 | 0.6 ± 0.1 | 0.04 ± 0.09 | -0.1 ± 0.1 | Cooperative |
| | L155A | N | 0.15 | 1.4 ± 0.1 | 0.64 ± 0.07 | 0.9 ± 0.1 | 0.79 ± 0.08 | 0.5 ± 0.2 | Cooperative |
| | W158F | N/Interface | 0 | 0.3 ± 0.1 | 0.84 ± 0.10 | 0.3 ± 0.1 | 0.82 ± 0.09 | 0.0 ± 0.2 | Cooperative |
| | L161A | N/Interface | 0 | 1.2 ± 0.1 | 0.36 ± 0.04 | 1.5 ± 0.3 | 0.26 ± 0.03 | -0.3 ± 0.3 | Cooperative |
| TM3 | L174A | N/Interface | 0 | 4.5 ± 0.2 | 0.49 ± 0.05 | 4.8 ± 0.1 | 0.17 ± 0.02 | -0.3 ± 0.2 | Cooperative |
| | T178A | N | 0.11 | 0.7 ± 0.1 | 0.77 ± 0.09 | 0.9 ± 0.2 | 0.66 ± 0.07 | -0.2 ± 0.2 | Cooperative |
| | S181A | N/Interface | 0 | 0.1 ± 0.2 | 0.92 ± 0.11 | 0.5 ± 0.1 | 0.85 ± 0.09 | -0.4 ± 0.2 | Cooperative |
| | F197A | N | 0.01 | 0.6 ± 0.2 | 0.02 ± 0.00 | 0.5 ± 0.2 | 0.00 ± 0.00 | 0.1 ± 0.2 | Cooperative |
| TM4 | S201T | C/Interface | 0 | 0.0 ± 0.2 | 0.05 ± 0.01 | 0.0 ± 0.2 | 0.01 ± 0.00 | 0.0 ± 0.3 | Cooperative |
| | A206G | C | 0 | 0.3 ± 0.2 | 1.07 ± 0.12 | 0.9 ± 0.1 | 0.92 ± 0.10 | -0.6 ± 0.2 | Cooperative |
| | L207A | C/Interface | 0 | 5.0 ± 0.2 | 0.12 ± 0.01 | 4.7 ± 0.1 | 0.16 ± 0.02 | 0.3 ± 0.2 | Cooperative |
| | Y210F | C | 0 | 0.5 ± 0.2 | 0.86 ± 0.10 | 0.9 ± 0.1 | 0.61 ± 0.07 | -0.4 ± 0.2 | Cooperative |
| | R214A | C/Interface | 0.10 | 0.2 ± 0.2 | 0.70 ± 0.08 | 0.7 ± 0.2 | 0.52 ± 0.05 | -0.5 ± 0.3 | Cooperative |
| TM5 | I223A | C/Interface | 0.00 | 0.1 ± 0.2 | 0.56 ± 0.06 | 0.6 ± 0.2 | 0.30 ± 0.03 | -0.6 ± 0.2 | Moderate/C |
| | L225A | C/Interface | 0.03 | 0.2 ± 0.1 | 0.42 ± 0.05 | 0.4 ± 0.1 | 0.26 ± 0.03 | -0.3 ± 0.1 | Cooperative |
| | Q226A | C | 0.57 | 0.0 ± 0.1 | 0.95 ± 0.11 | -0.1 ± 0.2 | 0.67 ± 0.07 | 0.1 ± 0.2 | Cooperative |
| TM6 | M249A | C/Interface | 0.01 | 0.3 ± 0.1 | 0.97 ± 0.11 | 0.4 ± 0.2 | 0.79 ± 0.09 | -0.1 ± 0.2 | Cooperative |
| | A253V | C/Interface | 0 | 0.0 ± 0.1 | 0.03 ± 0.01 | 0.3 ± 0.2 | 0.03 ± 0.00 | -0.3 ± 0.2 | Cooperative |
| | H254A | C | 0 | 0.7 ± 0.1 | 0.05 ± 0.01 | -0.2 ± 0.2 | 0.01 ± 0.00 | 0.9 ± 0.2 | Moderate/Over |
| | G261A | C | 0 | 4.9 ± 0.1 | 0.05 ± 0.01 | 4.0 ± 0.1 | -0.02 ± 0.01 | 1.0 ± 0.2 | Moderate/Over |
| | A265V | C | 0 | 1.6 ± 0.3 | 0.37 ± 0.04 | 1.4 ± 0.2 | 0.29 ± 0.03 | 0.2 ± 0.4 | Cooperative |
| | D268A | C/Interface | 0.15 | 1.8 ± 0.2 | 0.54 ± 0.06 | 1.5 ± 0.1 | 0.48 ± 0.05 | 0.3 ± 0.2 | Cooperative |

**Table S3. The mutation-induced changes in thermodynamic stability ($\Delta\Delta G^{o}_{WT\text{-}Mut}$) and the activities relative to wild type in DDM micelles.** $f_{ASA}$: the fraction of residue buried area. In the "Location" column, "N" and "C": N- and C-subdomains, respectively. In the "Cooperativity profile" column, "Moderate/N", "Moderate/C", and "Moderate/Over": moderately localized in N and C subdomains, and moderately overpropagated, respectively. The stabilities of the mutants marked with asterisks have previously been reported (*1*). The reproducibility of the data has been confirmed. Errors denote ± SD from fitting. The $\Delta\Delta G^{o}_{WT\text{-}Mut}$ and $\Delta\Delta\Delta G^{o}$ values are in kcal/mol.

| Secondary structure | Mutation | Location | $f_{ASA}$ | N-subdomain ($95_N172_M$) | | C-subdomain ($172_M267_C$) | | $\Delta\Delta\Delta G^{o}$ | Cooperativity profile |
|---|---|---|---|---|---|---|---|---|---|
| | | | | $\Delta\Delta G^{o}_{WT\text{-}Mut}$ | Rel.Activity | $\Delta\Delta G^{o}_{WT\text{-}Mut}$ | Rel.Activity | | |
| TM1 | M100A* | N | 0.14 | 3.0 ± 0.3 | 0.55 ± 0.06 | 2.2 ± 0.3 | 0.64 ± 0.05 | 0.5 ± 0.4 | Cooperative |
| | C104A* | N | 0 | 1.2 ± 0.3 | 0.69 ± 0.04 | 0.9 ± 0.1 | 0.70 ± 0.05 | 0.3 ± 0.3 | Cooperative |
| L1 | V119A | N | 0.05 | 2.0 ± 0.2 | 0.57 ± 0.06 | 0.9 ± 0.1 | 0.76 ± 0.05 | 1.1 ± 0.2 | Moderate/N |
| | L123A | N | 0.23 | 2.1 ± 0.2 | 0.55 ± 0.06 | 0.9 ± 0.1 | 0.68 ± 0.05 | 1.2 ± 0.2 | Local/N |
| | W125A | N | 0.09 | 2.8 ± 0.3 | 0.04 ± 0.10 | 1.7 ± 0.2 | 0.00 ± 0.01 | 1.1 ± 0.3 | Moderate/N |
| | K132A | N | 0.56 | 0.2 ± 0.3 | 0.71 ± 0.06 | 0.4 ± 0.1 | 0.52 ± 0.05 | -0.2 ± 0.3 | Cooperative |
| | F133A | N | 0.80 | 1.3 ± 0.3 | 0.84 ± 0.06 | 0.5 ± 0.2 | 0.87 ± 0.05 | 0.8 ± 0.3 | Cooperative |
| | F135A | N | 0.73 | 0.4 ± 0.2 | 0.93 ± 0.06 | 0.2 ± 0.1 | 0.69 ± 0.05 | 0.2 ± 0.3 | Cooperative |
| | W136A | N | 0.46 | 2.7 ± 0.2 | 0.00 ± 0.02 | 1.7 ± 0.1 | 0.00 ± 0.03 | 1.0 ± 0.2 | Moderate/N |
| | R137A | N | 0.04 | 4.1 ± 0.2 | 0.01 ± 0.01 | 2.8 ± 0.1 | 0.01 ± 0.01 | 1.3 ± 0.2 | Local/N |
| | Y138F* | N | 0.25 | 1.8 ± 0.2 | 0.95 ± 0.06 | 0.6 ± 0.1 | 0.93 ± 0.05 | 1.2 ± 0.2 | Local/N |
| | F139A | N | 0.38 | 2.0 ± 0.2 | 0.47 ± 0.06 | 1.0 ± 0.1 | 0.47 ± 0.05 | 1.0 ± 0.2 | Moderate/N |
| | T140A* | N | 0.20 | 1.6 ± 0.2 | 0.85 ± 0.06 | 0.7 ± 0.1 | 0.60. ± 0.03 | 0.9 ± 0.2 | Moderate/N |
| | L143A* | N | 0.25 | 2.3 ± 0.2 | 0.76 ± 0.06 | 1.4 ± 0.1 | 0.65 ± 0.05 | 0.9 ± 0.2 | Moderate/N |
| TM2 | H150A | N | 0.01 | 0.0 ± 0.3 | 0.05 ± 0.08 | 0.3 ± 0.2 | 0.02 ± 0.13 | -0.3 ± 0.3 | Cooperative |
| | N154A* | N/Interface | 0 | 1.2 ± 0.2 | 0.01 ± 0.04 | 1.2 ± 0.3 | 0.01 ± 0.02 | 0.0 ± 0.4 | Cooperative |
| | L155A | N | 0.15 | 2.2 ± 0.2 | 0.75 ± 0.05 | 1.6 ± 0.2 | 0.60 ± 0.03 | 0.6 ± 0.3 | Cooperative |
| | W158F* | N/Interface | 0 | 1.0 ± 0.2 | 0.92 ± 0.06 | 0.1 ± 0.1 | 0.85 ± 0.05 | 0.9 ± 0.2 | Moderate/N |
| | L161A* | N/Interface | 0 | 2.0 ± 0.3 | 0.16 ± 0.06 | 2.7 ± 0.3 | 0.10 ± 0.06 | -0.7 ± 0.4 | Moderate/C |
| TM3 | L174A* | N/Interface | 0 | 3.7 ± 0.2 | 0.35 ± 0.06 | 3.3 ± 0.1 | 0.07 ± 0.07 | 0.4 ± 0.2 | Cooperative |
| | T178A* | N | 0.11 | 0.6 ± 0.2 | 0.77 ± 0.06 | 0.3 ± 0.1 | 0.66 ± 0.07 | 0.3 ± 0.2 | Cooperative |
| | S181A* | N/Interface | 0 | -0.6 ± 0.2 | 1.03 ± 0.06 | 0.6 ± 0.1 | 1.00 ± 0.05 | -1.2 ± 0.2 | Moderate/Over |
| | F197A | N | 0.01 | 1.7 ± 0.2 | 0.01 ± 0.03 | 0.6 ± 0.1 | 0.00 ± 0.07 | 1.1 ± 0.2 | Moderate/N |
| TM4 | S201T* | C/Interface | 0 | 0.4 ± 0.2 | 0.02 ± 0.01 | 0.8 ± 0.2 | 0.00 ± 0.03 | -0.4 ± 0.3 | Cooperative |
| | A206G | C | 0 | 0.4 ± 0.2 | 0.09 ± 0.09 | 0.6 ± 0.1 | 0.09 ± 0.06 | -0.2 ± 0.2 | Cooperative |
| | L207A* | C/Interface | 0 | 4.1 ± 0.3 | 0.12 ± 0.01 | 2.7 ± 0.1 | 0.16 ± 0.02 | 1.4 ± 0.3 | Local/N |
| | Y210F* | C | 0 | 1.9 ± 0.2 | 0.50 ± 0.07 | 1.2 ± 0.1 | 0.66 ± 0.05 | 0.8 ± 0.2 | Moderate/N |
| | R214A | C/Interface | 0.10 | 0.9 ± 0.2 | 0.41 ± 0.06 | 0.6 ± 0.1 | 0.43 ± 0.05 | 0.3 ± 0.3 | Cooperative |
| TM5 | I223A | C/Interface | 0.00 | 1.0 ± 0.3 | 0.24 ± 0.06 | 0.5 ± 0.1 | 0.23 ± 0.11 | 0.5 ± 0.3 | Cooperative |
| | L225A* | C/Interface | 0.03 | -0.7 ± 0.2 | 0.27 ± 0.07 | 1.0 ± 0.1 | 0.10 ± 0.06 | -1.6 ± 0.2 | Local/C |
| | Q226A* | C | 0.57 | 0.2 ± 0.2 | 0.82 ± 0.06 | 0.8 ± 0.2 | 0.51 ± 0.05 | -0.6 ± 0.3 | Moderate/C |
| TM6 | M249A | C/Interface | 0.01 | 0.3 ± 0.2 | 0.59 ± 0.06 | 0.5 ± 0.2 | 0.85 ± 0.05 | -0.2 ± 0.3 | Cooperative |
| | A253V* | C/Interface | 0 | 1.5 ± 0.2 | 0.06 ± 0.01 | 0.9 ± 0.1 | 0.00 ± 0.06 | 0.6 ± 0.3 | Moderate/Over |
| | H254A | C | 0 | 1.5 ± 0.2 | 0.05 ± 0.01 | -0.3 ± 0.1 | 0.01 ± 0.05 | 1.8 ± 0.3 | Over |
| | G261A* | C | 0 | 4.0 ± 0.2 | 0.05 ± 0.01 | 2.7 ± 0.1 | -0.01 ± 0.06 | 1.3 ± 0.2 | Over |
| | A265V* | C | 0 | 2.3 ± 0.2 | 0.30 ± 0.06 | 1.3 ± 0.1 | 0.13 ± 0.05 | 1.0 ± 0.2 | Moderate/Over |
| | D268A* | C/Interface | 0.15 | 2.4 ± 0.2 | 0.44 ± 0.07 | 1.3 ± 0.1 | 0.28 ± 0.05 | 1.1 ± 0.2 | Moderate/Over |

**Table S4. Fitted parameters of the time-dependent contact autocorrelation data to a triple exponential decay function for the whole (A), headgroup (B), and tail (C) regions of the lipid (Lip) or detergent (Det) molecules on GlpG (Prot) and on themselves.** $A$: % amplitude; $\tau_R$: residence time; $<\tau_R>$: the amplitude-weighted average residence time; Adj-$R^2$: the adjusted $R$-square; $\Delta G^o_{Solv}$: the solvation free energy of an amphiphile molecule on the protein. $\tau_{R,1/e}$ denotes the resident time at which the contact autocorrelation decays to 1/$e$ of the initial value. Errors denote ± SD from fitting.

**A**

| Whole | $A_1$ (%) | $\tau_{R,1}$ (ns) | $A_2$ (%) | $\tau_{R,2}$ (ns) | $A_3$ (%) | $\tau_{R,3}$ (ns) | $A_\infty$ (%) | $<\tau_R>$ (ns) | Adj-$R^2$ | $\tau_{R,1/e}$ (ns) |
|---|---|---|---|---|---|---|---|---|---|---|
| Lip–Lip | 14.2 ± 0.4 | 1.3 ± 0.1 | 39.8 ± 0.5 | 21 ± 1 | 46.0 ± 0.5 | 82 ± 1 | 0.0 ± 0.0 | 46 ± 1 | 0.999 | 36 ± 1 |
| Prot–Lip | 29.4 ± 0.6 | 19 ± 1 | 54.3 ± 0.5 | 112 ± 2 | 11.1 ± 0.6 | 527 ± 30 | 1.3 ± 0.1 | 132 ± 7 | 0.999 | 83 ± 1 |
| $\Delta G^o_{Solv,Lip}$ = –0.62 ± 0.03 kcal/mol from $<\tau_R>$ | | | | | | $\Delta G^o_{Solv,Lip}$ = –0.50 ± 0.02 kcal/mol from $\tau_{R,1/e}$ | | | | |
| Det–Det120 | 30.5 ± 0.9 | 4.1 ± 0.2 | 50.3 ± 1.3 | 28 ± 1 | 18.1 ± 1.8 | 85 ± 4 | 0.0 ± 0.0 | 31 ± 2 | 0.998 | 22 ± 1 |
| Prot–Det120 | 23.8 ± 0.4 | 4.1 ± 0.1 | 35.8 ± 0.3 | 58 ± 1 | 39.7 ± 0.3 | 302 ± 2 | 0.4 ± 0.0 | 143 ± 2 | 0.999 | 92 ± 1 |
| $\Delta G^o_{Solv,Det120}$ = –0.90 ± 0.05 kcal/mol from $<\tau_R>$ | | | | | | $\Delta G^o_{Solv,Det120}$ = –0.85 ± 0.03 kcal/mol from $\tau_{R,1/e}$ | | | | |
| Det–Det150 | 16.6 ± 1.5 | 3.2 ± 0.4 | 39.4 ± 1.2 | 18 ± 1 | 43.7 ± 1.5 | 66 ± 1 | 0.0 ± 0.0 | 37 ± 2 | 0.998 | 29 ± 1 |
| Prot–Det150 | 25.8 ± 0.3 | 6.2 ± 0.1 | 48.6 ± 0.4 | 83 ± 1 | 24.1 ± 0.5 | 307 ± 4 | 0.1 ± 0.0 | 118 ± 3 | 0.999 | 81 ± 1 |
| $\Delta G^o_{Solv,Det150}$ = –0.69 ± 0.03 kcal/mol from $<\tau_R>$ | | | | | | $\Delta G^o_{Solv,Det150}$ = –0.61 ± 0.02 kcal/mol from $\tau_{R,1/e}$ | | | | |

**B**

| Headgroup | $A_1$ (%) | $\tau_{R,1}$ (ns) | $A_2$ (%) | $\tau_{R,2}$ (ns) | $A_3$ (%) | $\tau_{R,3}$ (ns) | $A_\infty$ (%) | $<\tau_R>$ (ns) | Adj-$R^2$ | $\tau_{R,1/e}$ (ns) |
|---|---|---|---|---|---|---|---|---|---|---|
| Lip–Lip | 37.0 ± 0.5 | 2.2 ± 0.1 | 31.9 ± 2.4 | 27 ± 1 | 31.1 ± 2.6 | 64 ± 2 | 0.0 ± 0.0 | 29 ± 3 | 0.999 | 22 ± 1 |
| Prot–Lip | 33.1 ± 0.4 | 8.9 ± 0.2 | 51.8 ± 0.3 | 104 ± 1 | 9.8 ± 0.3 | 725 ± 39 | 0.5 ± 0.1 | 135 ± 6 | 0.998 | 64 ± 1 |
| $\Delta G^o_{SolvEx,LipHead}$ = –0.91 ± 0.06 kcal/mol from $<\tau_R>$ | | | | | $\Delta G^o_{SolvEx,LipHead}$ = –0.63 ± 0.03 kcal/mol from $\tau_{R,1/e}$ | | | | | |
| Det–Det120 | 15.0 ± 2.7 | 1.5 ± 0.3 | 45.0 ± 2.4 | 6.1 ± 0.3 | 40.1 ± 0.7 | 34 ± 1 | 0.0 ± 0.0 | 17 ± 1 | 0.999 | 11 ± 1 |
| Prot–Det120 | 31.3 ± 0.8 | 3.3 ± 0.1 | 38.7 ± 0.7 | 28 ± 1 | 28.6 ± 0.9 | 110 ± 2 | 1.5 ± 0.0 | 44 ± 2 | 0.998 | 27 ± 1 |
| $\Delta G^o_{SolvEx,Det120Head}$ = –0.58 ± 0.03 kcal/mol from $<\tau_R>$ | | | | | $\Delta G^o_{SolvEx,Det120Head}$ = –0.53 ± 0.06 kcal/mol from $\tau_{R,1/e}$ | | | | | |
| Det–Det150 | 13.3 ± 1.1 | 0.1 ± 0.3 | 44.5 ± 1.1 | 6.4 ± 0.2 | 41.9 ± 0.6 | 38 ± 1 | 0.0 ± 0.0 | 19 ± 1 | 0.998 | 12 ± 1 |
| Prot–Det150 | 21.0 ± 0.7 | 0.9 ± 0.1 | 40.5 ± 0.5 | 18 ± 1 | 38.2 ± 0.5 | 106 ± 1 | 0.3 ± 0.0 | 48 ± 1 | 0.999 | 30 ± 1 |
| $\Delta G^o_{SolvEx,Det150Head}$ = –0.56 ± 0.02 kcal/mol from $<\tau_R>$ | | | | | $\Delta G^o_{SolvEx,Det150Head}$ = –0.54 ± 0.02 kcal/mol from $\tau_{R,1/e}$ | | | | | |

**C**

| Tail | $A_1$ (%) | $\tau_{R,1}$ (ns) | $A_2$ (%) | $\tau_{R,2}$ (ns) | $A_3$ (%) | $\tau_{R,3}$ (ns) | $A_\infty$ (%) | $<\tau_R>$ (ns) | Adj-$R^2$ | $\tau_{R,1/e}$ (ns) |
|---|---|---|---|---|---|---|---|---|---|---|
| Lip–Lip | 25.2 ± 0.5 | 1.4 ± 0.1 | 30.8 ± 0.5 | 19 ± 1 | 44.1 ± 0.5 | 81 ± 1 | 0.0 ± 0.0 | 42 ± 1 | 0.999 | 30 ± 1 |
| Prot–Lip | 33.8 ± 0.4 | 5.5 ± 0.1 | 44.3 ± 0.5 | 79 ± 1 | 17.2 ± 0.5 | 323 ± 8 | 1.6 ± 0.0 | 97 ± 3 | 0.999 | 58 ± 1 |
| $\Delta G^o_{SolvEx,LipTail}$ = –0.50 ± 0.02 kcal/mol from $<\tau_R>$ | | | | | $\Delta G^o_{SolvEx,LipTail}$ = –0.39 ± 0.02 kcal/mol from $\tau_{R,1/e}$ | | | | | |
| Det–Det120 | 48.5 ± 0.7 | 1.6 ± 0.1 | 29.4 ± 0.8 | 17 ± 1 | 21.9 ± 1.0 | 67 ± 2 | 0.0 ± 0.0 | 21 ± 1 | 0.997 | 9 ± 1 |
| Prot–Det120 | 32.1 ± 0.4 | 3.6 ± 0.1 | 29.8 ± 0.3 | 61 ± 1 | 36.5 ± 0.3 | 388 ± 3 | 0.5 ± 0.0 | 164 ± 2 | 0.999 | 87 ± 1 |
| $\Delta G^o_{SolvEx,Det120Tail}$ = –1.22 ± 0.03 kcal/mol from $<\tau_R>$ | | | | | $\Delta G^o_{SolvEx,Det120Tail}$ = –1.34 ± 0.07 kcal/mol from $\tau_{R,1/e}$ | | | | | |
| Det–Det150 | 36.2 ± 1.7 | 0.9 ± 0.1 | 28.7 ± 1.7 | 8.8 ± 1.0 | 34.6 ± 1.2 | 55 ± 2 | 0.5 ± 0.1 | 22 ± 1 | 0.998 | 11 ± 1 |
| Prot–Det150 | 37.0 ± 0.3 | 4.6 ± 0.1 | 36.6 ± 0.3 | 76 ± 1 | 23.9 ± 0.4 | 314 ± 3 | 0.2 ± 0.0 | 107 ± 2 | 0.999 | 58 ± 1 |
| $\Delta G^o_{SolvEx,Det150Tail}$ = –0.94 ± 0.04 kcal/mol from $<\tau_R>$ | | | | | $\Delta G^o_{SolvEx,Det150Tail}$ = –0.98 ± 0.05 kcal/mol from $\tau_{R,1/e}$ | | | | | |

**Table S5.** Equilibrium folding data of OmpLA. The fitted stabilities ($\Delta G^o$) and *m*-values (the dependence of $\Delta G^o$ on GdnHCl concentration) were obtained from a three-state model. For the titrations with DC12PC, we used the fixed *m* values previously determined by global fitting on WT and many variants ($m_{N\text{-}I}$ = 2.0 kcal·mol$^{-1}$·M$^{-1}$ and $m_{I\text{-}U}$ = 7.2 kcal·mol$^{-1}$·M$^{-1}$). For the titrations with DC11PC, we floated both *m* values during fitting.

| | $\Delta G^o_{N\text{-}I,l,w}$ (kcal/mol) | $m_{N\text{-}I}$ (kcal/mol·M$^{-1}$) | $\Delta G^o_{I\text{-}U,l,w}$ (kca/mol) | $m_{I\text{-}U}$ (kcal/mol·M$^{-1}$) | $\Delta G^o_{N\text{-}U,l,w}$ (kcal/mol) |
|---|---|---|---|---|---|
| DC12PC | 5.4 ± 0.5 | 2.0 | 26.4 ± 0.1 | 7.2 | 31.8 ± 0.6 |
| DC11PC | 9.3 ± 4.8 | 4.7 ± 2.6 | 16.4 ± 1.4 | 4.2 ± 0.4 | 25.7 ± 6.2 |

### References


1. R. Guo *et al.*, Steric trapping reveals a cooperativity network in the intramembrane protease GlpG. *Nat Chem Biol* **12**, 353-360 (2016).
2. M. Howarth *et al.*, Monovalent, reduced-size quantum dots for imaging receptors on living cells. *Nat Methods* **5**, 397-399 (2008).
3. D. E. Hyre, I. Le Trong, S. Freitag, R. E. Stenkamp, P. S. Stayton, Ser45 plays an important role in managing both the equilibrium and transition state energetics of the streptavidin-biotin system. *Protein Sci* **9**, 878-885 (2000).
4. Y. C. Chang, J. U. Bowie, Measuring membrane protein stability under native conditions. *Proc Natl Acad Sci U S A* **111**, 219-224 (2014).
5. T. Parasassi, G. De Stasio, G. Ravagnan, R. M. Rusch, E. Gratton, Quantitation of lipid phases in phospholipid vesicles by the generalized polarization of Laurdan fluorescence. *Biophys J* **60**, 179-189 (1991).
6. M. Beaugrand *et al.*, Lipid concentration and molar ratio boundaries for the use of isotropic bicelles. *Langmuir* **30**, 6162-6170 (2014).
7. K. S. Mineev, K. D. Nadezhdin, S. A. Goncharuk, A. S. Arseniev, Characterization of Small Isotropic Bicelles with Various Compositions. *Langmuir* **32**, 6624-6637 (2016).
8. M. Kieber *et al.*, The Fluidity of Phosphocholine and Maltoside Micelles and the Effect of CHAPS. *Biophys J* **116**, 1682-1691 (2019).
9. G. C. Chow, Tests of Equality Between Sets of Coefficients in Two Linear Regressions. *Econometrica* **28**, 591-605 (1960).
10. Y. Wang, Y. Zhang, Y. Ha, Crystal structure of a rhomboid family intramembrane protease. *Nature* **444**, 179-180 (2006).
11. S. Jo, T. Kim, V. G. Iyer, W. Im, CHARMM-GUI: a web-based graphical user interface for CHARMM. *J Comput Chem* **29**, 1859-1865 (2008).
12. D. M. Kruger, S. C. L. Kamerlin, Micelle Maker: An Online Tool for Generating Equilibrated Micelles as Direct Input for Molecular Dynamics Simulations. *ACS Omega* **2**, 4524-4530 (2017).
13. R. B. Best *et al.*, Optimization of the additive CHARMM all-atom protein force field targeting improved sampling of the backbone phi, psi and side-chain chi(1) and chi(2) dihedral angles. *J Chem Theory Comput* **8**, 3257-3273 (2012).
14. S. Pall *et al.*, Heterogeneous parallelization and acceleration of molecular dynamics simulations in GROMACS. *J Chem Phys* **153**, 134110 (2020).
15. S. K. McDonald, K. G. Fleming, Aromatic Side Chain Water-to-Lipid Transfer Free Energies Show a Depth Dependence across the Membrane Normal. *J Am Chem Soc* **138**, 7946-7950 (2016).
16. C. P. Moon, K. G. Fleming, Side-chain hydrophobicity scale derived from transmembrane protein folding into lipid bilayers. *Proc Natl Acad Sci U S A* **108**, 10174-10177 (2011).
17. P. J. Fleming, J. A. Freites, C. P. Moon, D. J. Tobias, K. G. Fleming, Outer membrane phospholipase A in phospholipid bilayers: a model system for concerted computational and experimental investigations of amino acid side chain partitioning into lipid bilayers. *Biochim Biophys Acta* **1818**, 126-134 (2012).
18. C. P. Moon, N. R. Zaccai, P. J. Fleming, D. Gessmann, K. G. Fleming, Membrane protein thermodynamic stability may serve as the energy sink for sorting in the periplasm. *Proc Natl Acad Sci U S A* **110**, 4285-4290 (2013).

19. E. L. Wu *et al.*, E. coli outer membrane and interactions with OmpLA. *Biophys J* **106**, 2493-2502 (2014).
20. H. J. Snijder *et al.*, Structural evidence for dimerization-regulated activation of an integral membrane phospholipase. *Nature* **401**, 717-721 (1999).
21. N. Michaud-Agrawal, E. J. Denning, T. B. Woolf, O. Beckstein, MDAnalysis: a toolkit for the analysis of molecular dynamics simulations. *J Comput Chem* **32**, 2319-2327 (2011).
22. N. Bernhardt, J. D. Faraldo-Gomez, MOSAICS: A software suite for analysis of membrane structure and dynamics in simulated trajectories. *Biophys J* **122**, 2023-2040 (2023).
23. D. Min, R. E. Jefferson, J. U. Bowie, T. Y. Yoon, Mapping the energy landscape for second-stage folding of a single membrane protein. *Nat Chem Biol* **11**, 981-987 (2015).
24. W. Lu, N. P. Schafer, P. G. Wolynes, Energy landscape underlying spontaneous insertion and folding of an alpha-helical transmembrane protein into a bilayer. *Nat Commun* **9**, 4949 (2018).
25. H. K. Choi *et al.*, Watching helical membrane proteins fold reveals a common N-to-C-terminal folding pathway. *Science* **366**, 1150-1156 (2019).
26. K. A. Gaffney *et al.*, Lipid bilayer induces contraction of the denatured state ensemble of a helical-bundle membrane protein. *Proc Natl Acad Sci U S A* **119**, (2022).
27. R. C. Oliver *et al.*, Dependence of micelle size and shape on detergent alkyl chain length and head group. *PLoS One* **8**, e62488 (2013).
28. T. A. Caldwell *et al.*, Low- q Bicelles Are Mixed Micelles. *J Phys Chem Lett* **9**, 4469-4473 (2018).
